# The adaptive landscapes of three global *Escherichia coli* transcriptional regulators

**DOI:** 10.1101/2024.11.11.623025

**Authors:** Cauã Antunes Westmann, Leander Goldbach, Andreas Wagner

## Abstract

The evolution of gene regulation is a major source of evolutionary adaptation and innovation, particularly when organisms encounter new or changing environments. Central to this process is the emergence of new transcription factor binding sites (TFBSs). Adaptive landscapes provide a powerful framework to study such emergence by linking regulatory DNA sequences to their transcriptional outputs. Although several landscapes have been characterized for DNA, RNA, and proteins, large-scale in vivo adaptive landscapes for bacterial TFBSs remain scarce. Here, we address this gap by experimentally mapping the first comprehensive in vivo regulatory landscapes for three global transcription factors in *Escherichia coli*: CRP, Fis, and IHF. Using a massively parallel reporter assay, we quantify the regulation strength of more than 30,000 TFBS variants for each factor and reconstruct their adaptive landscapes. All three landscapes are highly rugged and exhibit pervasive epistasis, with thousands of local peaks distributed broadly across sequence space. This ruggedness contrasts sharply with the much smoother TFBS landscapes of eukaryotes. It suggests greater constraints on the evolution of prokaryotic gene regulation. Nonetheless, evolutionary simulations show that ∼10% of evolving populations can reach a peak of strong regulation, a proportion that is significantly greater than in comparable random landscapes. Adaptive evolution starting from the same DNA sequence can attain different high peaks, and some peaks are reached more frequently than others. Together, our results show that de novo adaptive evolution of new gene regulation in bacteria is feasible, but subject to a blend of chance, historical contingency, and evolutionary biases.

## INTRODUCTION

Transcriptional regulation controls the ability of RNA polymerase to initiate the transcription of a gene into mRNA^1^. It is crucial in the life of all organisms, orchestrating the expression of genes in response to environmental cues and cellular states^2,3^. Transcriptional regulation is mediated by transcription factors (TFs), proteins that bind DNA near a gene at short DNA words known as transcription factor binding sites (TFBSs). The binding of a TF to its binding site on DNA can either hinder or facilitate transcription initiation by RNA polymerase^1,4,5^. The stronger the binding of a TF to its TFBS is, the more strongly the TF can regulate a nearby gene^6–9^. In bacteria, TFs operate within a hierarchically organized gene regulatory network. At the bottom of this hierarchy are *local* TFs that regulate the expression of one or few genes and modulate specific biological processes ^10–12^. At the top are *global* TFs that may regulate hundreds of genes and control many cellular functions ^11,13–15^. Consistent with their broader role, global regulators are typically more highly expressed and bind to a broader range of TFBSs than local regulators^12^.

A genotype-phenotype map is a conceptual analog of a physical landscape, where each location corresponds to one genotype in sequence space, and is associated with a quantitative phenotype. If the phenotype is related to gene expression, the map is also called a regulatory landscape^6,16–19^. Another special case of such a map is a fitness landscape or adaptive landscape, in which the phenotype is fitness and is interpreted as an elevation^20,21^. Over the last decade, genotype–phenotype (GP) maps and fitness landscapes have become central tools for understanding how molecular systems evolve under mutation and selection^22–25^. Such maps and landscapes have been experimentally studied for DNA^6,8,18,19,26,27^, protein^28–32^ and RNA^33–35^ molecules, revealing key topographical properties that shape evolutionary outcomes, including epistasis^24,36^—the non-additive effects of multiple mutations on phenotype—landscape ruggedness, reflected in the number and distribution of fitness peaks, and constraints on adaptive evolution. For example, one large-scale study adopted CRISPR-Cas9 technology to measure the fitness of more than 200,000 *E. coli* genotypes that encode variants of the bacterial antibiotic resistance gene dihydrofolate reductase (DHFR). It showed that this landscape is highly rugged (multi-peaked)^31^.

For transcription factor binding sites, most pertinent large-scale studies are based on in vitro binding assays, such as protein-binding microarrays (PBMs), and they focus predominantly on eukaryotic transcription factors^6^. While these studies have been instrumental in characterizing transcription factor binding preferences, they typically do not measure regulatory output in a native cellular context. In contrast, comprehensive in vivo data for bacterial TFBSs remain extremely rare. To our knowledge, only two high-resolution in vivo landscapes have been previously mapped for bacterial regulators, those of the local regulators TetR^18^ and LacI^27^. As a result, it remains unclear whether principles inferred from protein landscapes, eukaryotic TFBSs, or in vitro binding assays generalize to transcriptional regulation in bacteria, particularly for global regulators^11^ that integrate multiple physiological signals.

Both TFs and their TFBSs evolve, but TFBSs evolve more rapidly. The reason is that a mutation in a TF can affect the expression of many genes, whereas a mutation in a TFBS may affect only one gene and is thus less likely to be deleterious ^13,37–39^. A special case of TFBS evolution is the evolution of a strong TFBS from a DNA word with weak or no regulatory activity. Such *de novo* evolution of TFBSs can create new regulatory interactions and change the structure of gene regulatory networks^40–43^. Unfortunately, we know little about the ability of Darwinian evolution to create TFBSs de novo. A strong TFBS may have to arise from a non-binding site through an evolutionary path of multiple mutational steps. Darwinian evolution can favor this process only if strong binding is adaptive and if each mutational step in a path increases binding strength, i.e., if the path is *accessible* to Darwinian evolution. To find whether such paths exist may require the analysis of multiple paths. This is challenging because sequence space contains an astronomical number of potential TFBSs for any one TF. The number of evolutionary paths to strong transcriptional regulation is thus also astronomical. For each such path, the strength of each TFBS along the path has to be measured experimentally ^44–47^.

In principle, one could attempt to construct such landscapes in silico using commonly employed models of TF–DNA interactions, such as position weight matrices (PWMs)^48–50^. However, PWMs are derived from a limited set of naturally occurring binding sites and primarily reflect sequence conservation rather than quantitative regulatory output. Moreover, PWMs assume independent and additive contributions of individual nucleotide positions to DNA binding ^48–50^. They therefore cannot capture the influences of epistatic interactions between positions, which can dramatically affect landscape topography and evolutionary accessibility^24^. Lastly, PWMs do not account for important biological effects that modulate gene regulation such as DNA shape^51,52^, cooperative interactions^53,54^, and chromosomal context^55,56^. Thus, experiments are necessary not only to obtain quantitative measurements of gene regulatory activity, but also to refine and inform PWM-based models using thousands of experimentally characterized sequences.

Building on our previous work on a local TF^18^, we address this challenge for three global regulators in *Escherichia coli* by performing three independent and massively parallel experiments^44^ for each TF. The experiments use a synthetic biology platform to quantify how strongly each of more than 30,000 binding sites for a TF can regulate the expression of a nearby reporter gene.

The first TF we study is the cAMP receptor protein (CRP). In the absence of glucose, CRP modulates the expression of genes mostly involved in carbon metabolism. It allows *E. coli* to efficiently switch between sources of carbon and energy ^57–60^. The second TF is the factor for inversion stimulation (Fis), which helps to regulate the expression of genes involved in growth phase transitions. It also modulates the supercoiling of DNA^61–63^ and influences DNA replication, recombination, and repair^61–63^. The third factor is the integration host factor (IHF). IHF regulates genes involved in stress responses and stationary phase survival^64,65^. Similar to Fis, it is involved in DNA compaction, replication, and recombination, but also in the assembly of complex nucleoprotein structures^64,65^. We chose these factors because they are the most global regulators in *E. coli*, and they are diverse, belonging to different protein families.

We use our experimental data for each TF’s binding sites to map the relationship between binding site genotype and gene expression. Our first aim is to characterize the resulting regulatory landscapes for global bacterial regulators, and to find out whether these landscapes are different or similar. When strong regulation of a gene is adaptive, a regulatory landscape becomes a fitness landscape ^20,21,25,66^. Our second aim is to understand how populations would evolve on each of our three landscapes when they are viewed as fitness landscapes. Specifically, we study how evolving populations would traverse each landscape through individual mutational steps that change a TFBS its ability to regulate a gene via its cognate TF. A peak in such a landscape is a TFBS conveying stronger regulation than all neighboring TFBSs in sequence space. If such a landscape is rugged (has multiple peaks), natural selection alone may not enable a population to discover the highest peaks, i.e., the strongest TFBSs. The reason is that the peaks may be separated by valleys of low regulation strength that cannot be traversed by natural selection alone ^67–69^ ^25^. In other words, high peaks may not be easily accessible through Darwinian evolution – they may be reachable by few or no evolutionary paths of single DNA mutations in which each mutational step increases TFBS strength^6^. One factor that can reduce peak accessibility is epistasis, which can reduce the predictability of evolutionary trajectories towards a peak ^24,36^.

To accomplish both aims, we first studied the topography of the three landscapes, and then simulated the dynamics of evolving populations on them. All three landscapes are highly rugged. They contain more than 2,000 peaks that are scattered through genotype space and are rife with epistatic interactions, in striking contrast to the comparatively smooth TFBS landscapes described for eukaryotic systems^6^. Despite these features, evolving populations can reach the strongest TFBSs in all three landscapes. Which of several high peak is reached is contingent on chance events during adaptive evolution.

## RESULTS

### Landscape mapping

We constructed a modular plasmid system based on the common backbone plasmid pCAW-Sort-Seq-V2^18^ (**Supplementary Table S1; Supplementary Figures S1–S2; Supplementary Methods 2–3**). This backbone contains all shared regulatory and reporter elements required for fluorescence-based measurements, but it encodes neither a TF nor its binding site(s) for any of the regulators studied here. From this backbone, we generated three TF-specific plasmid derivatives. Each of these plasmids encodes one of the global transcription factors CRP, Fis, or IHF under inducible control (hereafter pCAW-Sort-Seq-V2-CRP, pCAW-Sort-Seq-V2-Fis, and pCAW-Sort-Seq-V2-IHF; **Supplementary Figure S2**). In each of the TF-specific plasmids, a TFBS insertion site is positioned immediately upstream of the *gfp* reporter gene, such that TF binding represses the transcription of *gfp*. Consequently, stronger TF–DNA binding results in lower GFP expression, which enables a quantitative readout of a binding site’s regulation strength.

For each TF-specific plasmid, we constructed three kinds of variants. The first carries a wild-type (WT) TFBS for the corresponding TF upstream of *gfp*. It serves as a reference conferring wild-type regulation of the reporter gene. The second contains a complete TFBS library, in which we randomized the eight most information-rich base-pair positions of the respective binding site (**Supplementary Methods 4**; **Supplementary Table S3**), as determined from alignments of experimentally characterized binding sites curated in RegulonDB^70^. Each library comprised 4⁸ = 65,536 unique TFBS sequences, and we constructed three independent biological replicate plasmid libraries from them per TF (**Supplementary Methods 4–5**). The third variant lacks a promoter upstream of *gfp* and serves as a negative control that allows us to quantify cellular autofluorescence during fluorescence measurements (**Supplementary Table S1**).

We introduced each TF-specific plasmid into an *E. coli* host strain in which the corresponding chromosomal TF gene had been deleted (*Δcrp*, *Δfis*, or *Δihf*; **Supplementary Table S2**). This genetic background ensures that the focal transcription factor is expressed exclusively from the plasmid. Although the mutant strains grow more slowly than the WT, they reached similar cell densities during late exponential or early stationary phase, the growth phase at which we performed all measurements (**Supplementary Figure S3**). TF expression is controlled by a tetracycline-inducible promoter and can be precisely tuned using anhydrotetracycline (aTc), allowing us to regulate TF abundance independently of growth conditions and to isolate the effects of TF–DNA binding on transcriptional regulation (**Figure 1a**).

**Figure 1.**
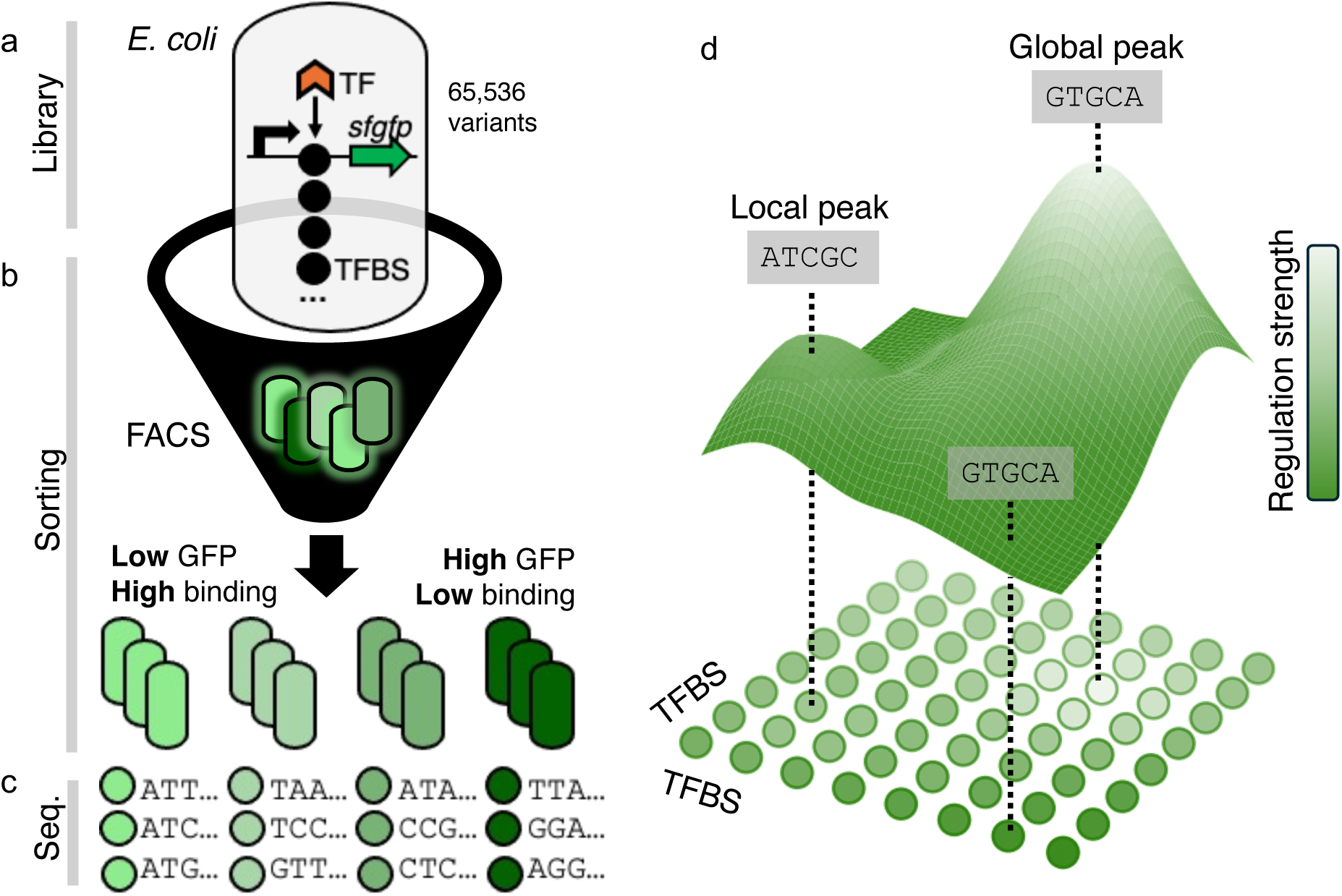
Mapping TFBS regulatory landscapes. **a-c) Sort-Seq procedure.** We utilized a plasmid-based fluorescence reporter system followed by sort-seq to map TFBS regulatory landscapes. **a. Library generation.** For generating our *E. coli* libraries, we cloned TFBS sequence variants (4^8^ = 65,536), represented as black circles, into our plasmid, between a σ^70^ constitutive promoter (shown as a black right-facing arrow) and a *gfp* gene (shown as a green right-facing arrow), to measure transcriptional repression through fluorescence intensity. When a TF binds to the TFBS, it blocks RNA polymerase activity through steric hindrance, reducing *gfp* transcription. Thus, lower fluorescence levels (light green) represent stronger TF-TFBS binding. The TFBS variants produce a range of regulation strengths resulting in variable green fluorescence intensities in bacterial cells (green-colored rounded rectangles). **b. Sorting procedure.** We sorted libraries into expression bins based on fluorescence intensity using a fluorescence-activated cell sorting (FACS, **Supplementary Figures S4-S6**). **c. Sequencing and phenotyping.** We sequenced TFBS variants from each fluorescence bin and used these data to calculate a continuous regulation strength for each genotype. Regulation strength (S) was computed as a weighted average of fluorescence across bins, based on the distribution of sequencing reads (see Methods). Regulation strength is visualized using a color gradient from dark green (low-affinity TFBSs, high fluorescence) to light green (high-affinity TFBSs, low fluorescence). **d) Genotype-phenotype mapping.** To construct a regulatory landscape, we connected TFBS genotypes (colored circles) that differed by a single nucleotide via edges, thereby establishing an interconnected genotype network. Each genotype conveys a regulatory phenotype (strength of regulation, heatmap colors), which can be viewed as the elevation dimension (z-axis) in a landscape.

We then mapped these TFBS genotypes to their respective regulatory phenotypes using a well-established technique known as sort-seq^16,44,71–73^ (**Figure 1b**), which combines fluorescence-activated cell sorting (FACS) with high-throughput sequencing (**Figure 1c**). In sort-seq, one first sorts cells into multiple fluorescence “bins” depending on the level of GFP expression. A cell’s GFP fluorescence serves as a proxy for GFP expression levels^74^ and quantifies how strongly the TFBS library member in this cell can regulate GFP expression (**Methods**, **Supplementary Methods 5**). For each TFBS genotype, we quantified regulation strength (S) as a weighted average of its sequencing counts across the different fluorescence bins, yielding a single continuous measure of regulatory activity (see **Methods**).

The results of our sort-seq experiments are three maps – one for each transcription factor – from each of more than 30,000 genotypes (TFBS variants) to regulatory phenotypes (regulation strength). One can view each map as a regulatory landscape^8,9,16,18,75–79^ (**Figure 1d**) that becomes a fitness landscape whenever strong regulation entails high fitness^6,25^. For the purpose of analyzing the landscape quantitatively, we represent it as a network of genotypes (TFBS variants). Each node in this network corresponds to a genotype. Edges link neighboring genotypes, which differ in a single nucleotide (**Figure 1d**).

### Landscapes exhibit diverse regulation strengths and distribution breadths

To evaluate the ability of our library to regulate gene expression, we first measured the distribution of GFP fluorescence intensities across the bins in two conditions, i.e., in the presence or absence of the TF (with or without the atc inducer, **Supplementary figures S4-S6**). In the presence of the TF, GFP expression was lower on average and showed a broader distribution than in the absence of the TF (**Supplementary figures S4-S6**). This indicates that the TFBSs in each library can indeed downregulate GFP expression, but some library variants convey stronger regulation than others, hence the broader fluorescence distribution.

We then pooled barcoded DNA sequences extracted from cells in each bin and biological replicate. We sequenced at least 250 unique TFBS genotypes from each bin, with an average of 3992.3±4293.9 unique sequences per bin for the three TFs (as detailed in **Methods**, **Supplementary Methods 7**, and **Supplementary Figures S4-S6**). The resulting sequences covered 95%, 90%, and 93% of the total library sizes (N = 65,536 genotypes) for CRP, Fis and IHF, respectively. To ensure the reliability of our data, we excluded sequences with low read coverage and sequences that were not present in all triplicates (**Methods, Supplementary Methods 7.3**). This quality filtering step resulted in library sizes of 31,975 genotypes for CRP (49% of all 4^8^ genotypes), 43,222 genotypes for Fis (66%), and 41,325 genotypes for IHF (63%). The correlation in read counts for each variant across replicates was very high for this quality-filtered data (Pearson’s R= 0.98-0.99) (**Supplementary Figures S7-S9**). We used this data for all further analyses.

The fluorescence and sequence data from each bin allowed us to quantify the variant’s ability to regulate gene expression for each TFBS library variant. We refer to the resulting metric as the *regulation strength S* conveyed by the variant (**Methods**, **Supplementary Methods 7.2**, **Figure 1d**). It ranges between S=0 (no regulation) to S=1 (strongest regulation among all variants in the library). Although our experiments directly quantify expression regulation, and not the affinity or binding strength of a TF to a TFBS variant, we also use *binding strength* as a proxy for regulation strength, because TF-DNA binding is necessary for regulation^1,4,11,54,80,81^. To each of our three TFs, we also assigned a reference TFBS that is naturally occurring and conveys strong regulation by the TF, as proven by previous experimental work^60,82–84^. We refer to this TFBS as the wild-type (WT ^82,85^ WT ^83,86^ and WT ^84,87^, **Supplementary Table S4**). We refer to TFBSs with regulation strengths below and above the wild-type (WT_CRP_: S_WT_=0.71, WT_Fis_: S_WT_=0.97, WT_IHF_: S_WT_=0.95) as weak and strong, respectively.

We observed a broad range of regulation strengths S for each TF landscape, with varying dispersions. The CRP landscape exhibited the lowest average regulation strength and a narrower distribution of S compared to Fis and IHF, which had similar distributions (mean±s.d., 0.37±0.1 for CRP, 0.57±0.13 for Fis, and 0.52±0.14 for IHF; see **Figure 2a, Supplementary Figure S10**). Next, we analyzed the strongly regulating TFBSs to identify nucleotides that may be particularly frequent, and thus potentially important for strong regulation (**Figure 2b**). We discovered a moderate association between the most frequent nucleotides (highlighted in yellow in **Figure 2b**) and the most informative nucleotides from the available position weight matrices (PWMs) for these TFs^70^. (Pearson correlation coefficients: R=0.51, R=0.43, R=0.47 for Fis, CRP and IHF, respectively, with p-values smaller than 10^-16^, rejecting the null hypothesis of an absence of association). This observation suggests that our logos capture different information compared to available PWMs. This is expected, because our approach allows us to filter sequences by regulation strength thresholds to construct our matrices, unlike the traditional method of aligning genomic TFBSs without considering their regulation strengths^48,49^.

**Figure 2.**
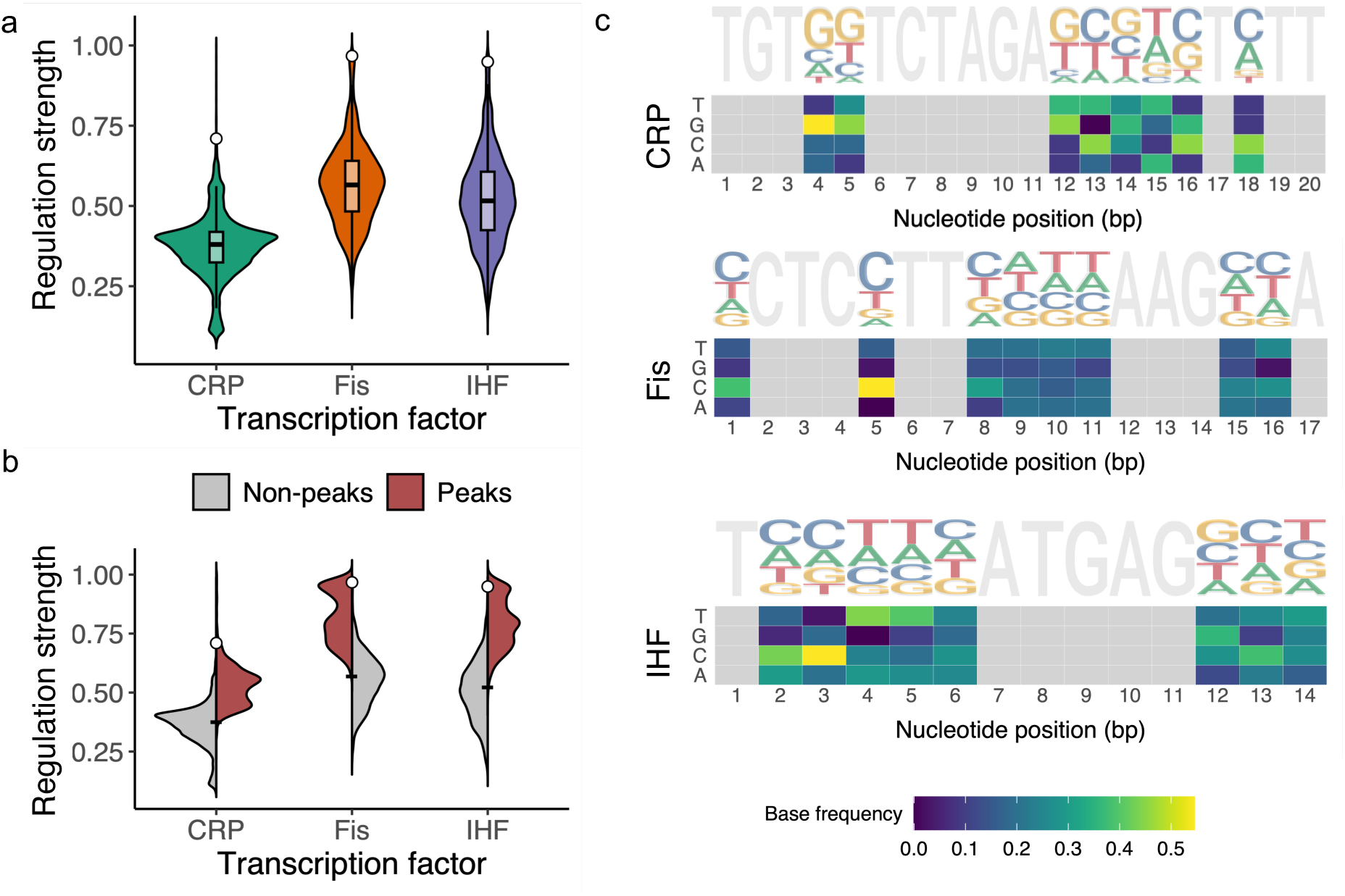
a. Genotypes in each landscape vary broadly in their regulation strength. The violin plots show the distribution of regulation strengths S (vertical axis) for each TF landscape (horizontal axis). The width of a plot at a given value of S represents the frequency of TFBSs at this value. The vertical length of the box in each violin plot covers the range between the first and third quartiles (IQR). The horizontal line within the box represents the median value, and whiskers span 1.5 times the IQR. The white circle shows the regulation strength of the wild type for each landscape (CRP: 0.71, Fis: 0.97, IHF: 0.95). The landscape sizes are as follows. CRP: N=31,975 TFBS variants; Fis: N= 43,222; IHF: N= 41,325. **b. Peak genotypes are stronger regulators than non-peak genotypes.** Dual violin plots show the distribution of regulation strength (vertical axis) for the three TF landscapes (horizontal axis), stratified by non-peak genotypes (grey, CRP, N=29,821, Fis, N= 40,910, IHF, N= 38,872), and peak genotypes (red, CRP, N= 2,154, Fis, N= 2,312, IHF, N= 2,453). The black tick-mark in each plot indicates the mean regulation strength of both non-peak and peak genotypes taken together (mean ± standard deviation, CRP: 0.37 ±0.1, Fis: 0.57±0.13, IHF: 0.52±0.14). The white circle on each plot marks the regulation strength of the wild-type (CRP: 0.71, Fis: 0.97, IHF: 0.95). **c. Sequence logos and nucleotide frequency matrices for strong CRP, Fis and IHF binding sites.** Each sequence logo ^48,49^ is based on an alignment of TFBSs with greater regulation strength than the wild-type for each of the three TFs IHF, Fis, and CRP. Each logo also shows the non-varying position (grey) of the TFBS genotype from which our libraries were created. In each logo, the height of each letter at each TFBS position indicates the information content at that nucleotide position – the taller the letter, the more frequent the nucleotide is in strongly regulating TFBSs ^48,49^. Similar information is conveyed by the frequency matrices displayed as heat maps below each logo. They represent the variability of each nucleotide at each position (horizontal axis) through a color gradient (see color legend). Tall letters in the sequence logo and yellow letters in the frequency matrix indicate frequent, and thus likely important nucleotides for strong regulation.

To further validate the data from our sort-seq experiments, we isolated cells harboring 10 different TFBS variants from each of 13 bins of each library (i.e., 10×13=130 variants per library), and determined their regulation strength more directly by quantifying GFP expression with a microplate reader (**Methods** and **Supplementary Figure S11**). This comparison validates the sort seq approach by revealing a strong association of the two independent quantifications of regulation strength (Pearson’s R=-0.81 for CRP, R=-0.74 for Fis, and R=-0.73 for IHF). (**Methods** and **Supplementary Figure S11**).

### All three regulatory landscapes are highly rugged

The study of our landscapes in a network framework (**Figure 1d**) can help to quantify different aspects of landscape topography (**Supplementary Table S5)**. One of them is the ruggedness of each landscape. It can be quantified by the number of peaks^31,88–90^. In the network framework, a peak is a TFBS whose neighbors are all weaker regulators (with lower S) than the TFBS itself. We find that all three landscapes are highly rugged, with 2,154, 2,312, and 2,453 peaks for the CRP, Fis, and IHF landscapes, respectively (**Figure 2c, Supplementary Figure S12, Supplementary Table S5**). Not surprisingly, peak genotypes generally are stronger regulators than non-peak genotypes (**Figure 2c**). Only a small fraction of peak genotypes regulate expression more strongly than the wild-type (**Figure 2c**; 61 for CRP, 172 for Fis, and 199 for IHF). We refer to such peaks as high peaks and distinguish them from low peaks (S<S_WT_).

The prevalence of sign epistasis (**Supplementary Table S5**) supports the notion that our landscapes are indeed rugged (see **Supplementary Methods 7.5** for further details on epistasis and its evolutionary consequences). Independent evidence for landscape ruggedness comes from comparing the ruggedness of our landscapes with that of a well-established theoretical model of uncorrelated random landscapes, in which each sequence is assigned a fitness at random from the same fitness distribution, and neighbors have uncorrelated fitness values^90–92^. Such landscapes are maximally rugged ^90–92^. We created 10^3^ uncorrelated random landscapes for each of our three TF landscapes by randomly shuffling the measured regulation strengths among all genotypes (**Supplementary Methods 7.6**). The number of peaks in our TF landscapes lies within 93%, 96%, and 98% percent of that of a maximally rugged random landscape, which, on average [mean±s.d.], has 2,308±133, 2,405±89 and 2,373±97 peaks for CRP, Fis and IHF, respectively. This analysis underlines that our landscapes are indeed highly rugged.

Because we use only quality-filtered genotype data, our landscapes lack regulatory information for about 40% of the 4^8^ genotypes. While we cannot exclude that this undersampling of genotypes has led to systematic biases in our determination of landscape ruggedness, we note that the sampling of landscape genotypes by our experiments was not strongly biased with respect to regulation strength. Specifically, when we analyzed the relative connectivity of genotypes in our landscapes – the fraction of each genotype’s 24 possible neighbors for which our experiments yielded regulatory data – we found that it is only weakly correlated with regulation strength (R=-0.1, -0.1, 0.01 for the CRP, Fis, and IHF landscapes, **Figures S13-S15**). Similarly, the relative connectivity of peak genotypes is only weakly correlated with their regulation strength (R=-0.05, -0.04, 0.06 for the CRP, Fis, and IHF landscapes).

### Landscape peaks are widely scattered in genotype space

In a rugged adaptive landscape, reaching high fitness peaks can be challenging, because such peaks are separated from other genotypes by valleys of low fitness that cannot be traversed by natural selection alone^25^. If natural selection favors strong gene regulation, the ruggedness of our regulatory landscapes may thus present a challenge for adaptive evolution. To better understand the potential magnitude of this challenge, we next analyzed our landscapes’ topography in greater detail. We began by studying the distribution of peaks in genotype space. If a landscape’s highest peaks are widely scattered through genotype space, then they may be accessible via fitness-increasing evolutionary paths from diverse non-peak genotypes. This may facilitate adaptive evolution compared to a landscape where peaks are clustered in a small region of genotype space.

For each of our three landscapes, we determined the genetic distance between peaks, i.e., the minimum number of mutations needed to transition from one peak to another, regardless of their effect on regulation strength. We compared the distribution of these distances to the distribution of genetic distances for an equal number of randomly selected non-peak variants. In all three landscapes, the distances between peaks are almost indistinguishable from those of random genotypes, differing on average by fewer than 0.1 mutational steps. In other words, peaks are about as widely dispersed in each landscape as random genotypes (**Supplementary Figure S16**). This is the case for both low and high peaks (**Supplementary Figures S17-S18**). A principal component analysis further underscores this dispersion (**Supplementary Methods 7.7, Supplementary Figures S19-S21**).

### Accessible paths to a peak are often not the shortest possible paths

Next, we focused on mutational paths to peaks that are evolutionarily accessible, i.e., a series of mutational steps where each step increases regulation strength. Specifically, we enumerated all accessible paths that exist from each non-peak genotype to each peak genotype for all three landscapes. We found that these paths are generally longer than the shortest genetic distance between a non-peak and the peak genotypes. Specifically, the mean length of accessible paths exceeded the shortest distance by 1.5, 1.8 and 1.8 steps for the CRP, Fis and IHF landscapes (**Supplementary Figure S22**). In other words, accessible mutational paths are often not the most direct paths.

The existence of indirect paths implies that some mutational steps are evolutionarily prohibited because they decrease regulation strength. The reason is closely linked to non-additive (epistatic) effects of two or more mutations on regulation strength. (see **Supplementary Methods 7.5** for further details on epistasis). More specifically, such inaccessible steps are a result of sign epistasis. In this kind of epistasis, a double mutant of a TFBS regulates expression more strongly than the TFBS itself, even though one or both constituent single mutants regulate expression more weakly than the TFBS^24,93–95^. Indeed, epistasis is prevalent in all three landscapes **(Table S5)**. Specifically, we observe sign epistasis in 62%, 66%, and 65% percent of interactions between single mutant pairs in the CRP, Fis, and IHF landscapes.

The existence of accessible paths alone does not tell us how easily a high peak can be found through Darwinian evolution. The reason is that only a very small fraction of all paths to that peak may be accessible, and an evolving population may not find any one of those paths. To quantify the fraction of accessible paths, we determined the total number of paths from each non-peak genotype to each high fitness peak, and computed the fraction of those paths that are accessible. **Figure 3a** shows how this fraction depends on the number of mutational steps in a path. It behaves similarly for all three landscapes (**Figure 3a**). The majority of two-step paths (a fraction greater than 80%) are accessible in all three landscapes (**Figure 3a**), but the fraction of accessible paths dwindles rapidly with an increasing number of mutational steps. It reaches a minimum below 1% for all three landscapes at eight mutational steps (**Figure 3a**).

**Figure 3.**
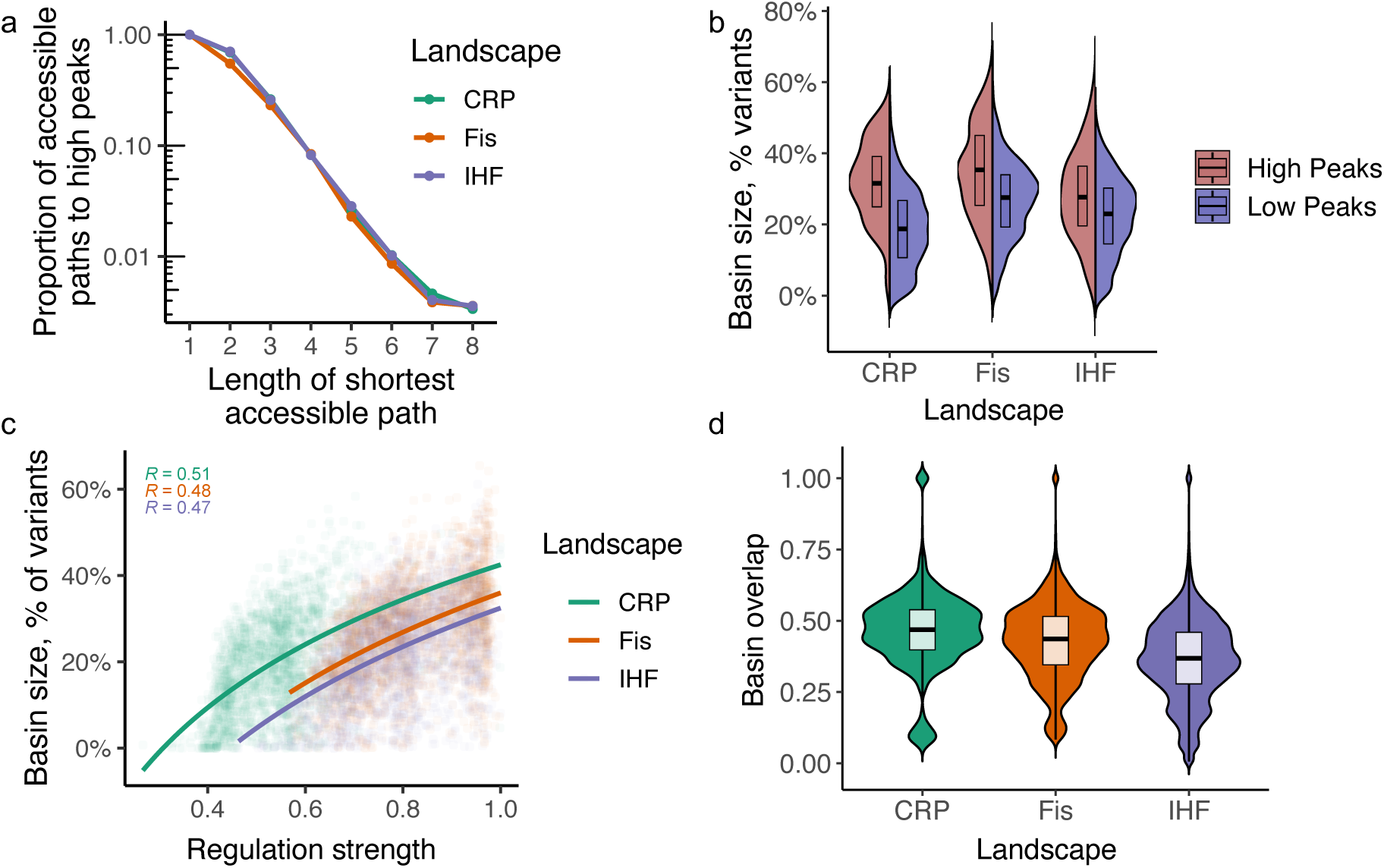
Peaks and their basins of attractions. **a. The fraction of accessible paths declines with increasing path length.** The figure illustrates how the fraction of accessible paths to high peaks (vertical axis, logarithmic scale) decreases with the length of the shortest accessible path (horizontal axis). Accessible paths are defined as paths where each step increases regulation, starting from a specified initial genotype. Each colored line corresponds to data from a different TF landscape: CRP (green), Fis (orange), and IHF (blue). Circles indicate the mean fraction of accessible paths for a given path length. **b. Higher peaks have larger basins of attraction.** The split-violin plots display the distribution of basin sizes (vertical axis) for high (red) and low (blue) peaks in the three TF landscapes CRP, Fis, and IHF (horizontal axis). High peaks have significantly larger basins of attraction (Welch Two Sample t-tests; CRP: t-value = 9.0898, df = 63.509, and p-value = 4.22 × 10^-13^, with mean basin sizes of x_high_ = 9528.426 and x_low_ = 5655.035. Fis: t-value = 7.617, df = 172.35, and p-value = 1.645 × 10^-12^, with mean basin sizes of x_high_ = 14206.70 and x_low_ = 10931.94. IHF: t-value = 6.1777, df = 201.99, and p-value = 3.521 × 10^-9^, with mean basin sizes of x_high_ = 10965.872 and x_low_ = 8664.953.). The violin plots show the distribution of basin sizes for each of the two kinds of peaks. Their width represents the frequency of a given basin size. The vertical length of the box in each plot covers the range between the first and third quartiles (IQR). The horizontal line within the box represents the median value, and whiskers span 1.5 times the IQR. **c. Peak genotypes with larger basins of attraction regulate expression more strongly.** The regulation strength of peak genotypes (horizontal axis) is plotted against the size of their basins of attraction (vertical axis, shown as a percentage of all (non-peak genotypes) for peaks in all three landscapes (color legend). Color-coded values of R at the top of the graph indicate Pearson correlation coefficients for each landscape (color legend). Curved lines are based on a linear regression analysis (in linear space), with grey shading indicating 95% confidence intervals (CRP: R^2^=0.28, N = 2,154; Fis: R^2^=0.24, N = 2,312; IHF: R^2^=0.23, N = 2,434). **d. Basins of attractions share many TFBS genotypes.** The violin plots with embedded boxplots illustrate the fraction of genotypes shared between basins of attractions (vertical axis) for all pairs of high peaks in the CRP (green), Fis (orange), and IHF (purple) TF landscapes (horizontal axis). Specifically, we computed basin overlap as one minus the Jaccard coefficient (**Supplementary Methods 7.5**). A value of one (zero) indicates that the basin of attraction of two peaks share all (no) genotypes (CRP: 61 high peaks, N=1,830 comparisons, Fis: 172 high peaks, N=14,706 comparisons, IHF: 199 high peaks, N=19,701 comparisons). Their width represents the frequency of a given basin overlap. The vertical length of the box in each box covers the range between the first and third quartiles (IQR). The horizontal line within the box represents the median value, and whiskers span 1.5 times the IQR.

### High peaks have large basins of attraction that share many TFBS genotypes

In the following analysis, we computed the basin of attraction of each peak, defined as the set of non-peak genotypes from which accessible paths to the peak exist^31,96,97^. In other words, a peak’s basin of attraction comprises all genotypes from which adaptive evolution can access the peak. The peaks in our landscapes have widely different basin sizes. They include peaks accessible from a mere fraction of genotypes to those accessible by a majority of genotypes (**Figure 3b**). Notably, high peaks generally had larger basins of attraction than low peaks (**Figure 3c, Supplementary Figure S23**). In addition, we found that many genotypes are part of multiple basins of attraction. Adaptive evolution may reach different high peaks starting from any such genotype. For all pairs of high peaks, we computed the pairwise overlap (intersection) of the basins of attraction, i.e., the fraction of genotypes that are part of both basins of attraction. (**Figure 3d, Supplementary Figures S24-S25**). The number of genotypes in this intersection varies widely, but the basins of attraction of high peaks generally share a substantial fraction of genotypes (mean shared genotypes: 48%, 42%, and 36% among all pairs of high peaks for the CRP, Fis, and IHF landscapes, **Figure 3d**)

### Genetic drift facilitates and clonal interference reduces the attainment of high peaks

Taken together, our analyses thus far indicate that high peaks in the CRP, Fis, and IHF landscapes are more accessible than low peaks (**Figure 3b**), and their basins of attraction also share more genotypes (**Figure 3d**). However, the accessible evolutionary paths to high peaks are often indirect. In addition, the fraction of accessible paths to any one high peak dwindles rapidly with the distance from that peak (**Figure 3a**).

To understand how these topographical features jointly affect adaptive evolution, we simulated the evolutionary dynamics on our three landscapes under the assumption that natural selection would favor strong regulation and that fitness is proportional to regulation strength.

Because high peaks constitute only a small fraction of all peaks (2.8%, 0.4%, and 0.5% in the CRP, Fis, and IHF landscapes), we hypothesized that most evolving populations would reach only low peaks. To test this hypothesis, we took advantage of the fact that *E.coli* has a low genomic mutation rate of 2.2 × 10^−10^ per base pair per generation ^67^, and that we study evolution only within an 8-nucleotide segment of a TFBS. In addition, *E. coli* has a large effective population size (>10^8^ individuals^67^), which means that genetic drift is weak and even minor differences in fitness are visible to natural selection ^67,98^. These conditions imply that a population on our landscape would evolve in the well-studied strong-selection weak-mutation regime (SSWM)^16,99–101^. In this regime, a population is monomorphic most of the time, until a beneficial mutation arises, which usually sweeps rapidly to fixation. In other words, adaptive evolution can be modeled as an adaptive walk, in which each mutational step is taken with a fixation probability that has been derived by Kimura ^98^. We thus call the resulting adaptive walks Kimura walks (**Supplementary Methods 8**). We simulated 10^3^ such adaptive walks starting from each of 15,000 randomly and uniformly selected starting (non-peak) TFBS genotypes from each landscape. Each random walk comprised up to 25 mutational fixation steps, unless it reached a fitness peak earlier. Each adaptive walk also accounted for known *E. coli* mutational biases ^102–104^ (**Supplementary Methods 8**).

As we hypothesized, most adaptive walks indeed reach only low peaks (90%, 85%, and 87% percent for the CRP, Fis, and IHF landscapes.). However, the percentage of walks that terminate at high peaks is several-fold greater than the proportion of high peaks. Specifically, 10 percent of walks terminate at high peaks in the CRP landscape, which is 3.6-fold higher than the percentage (2.8%) of high peaks in this landscape. In the Fis and IHF landscape, adaptive walks are 2 and 1.6-fold more likely to terminate at high peaks than expected from the proportion of these peaks among all peaks (7.4% and 8.1% percent) (**Figure 4a**).

**Figure 4.**
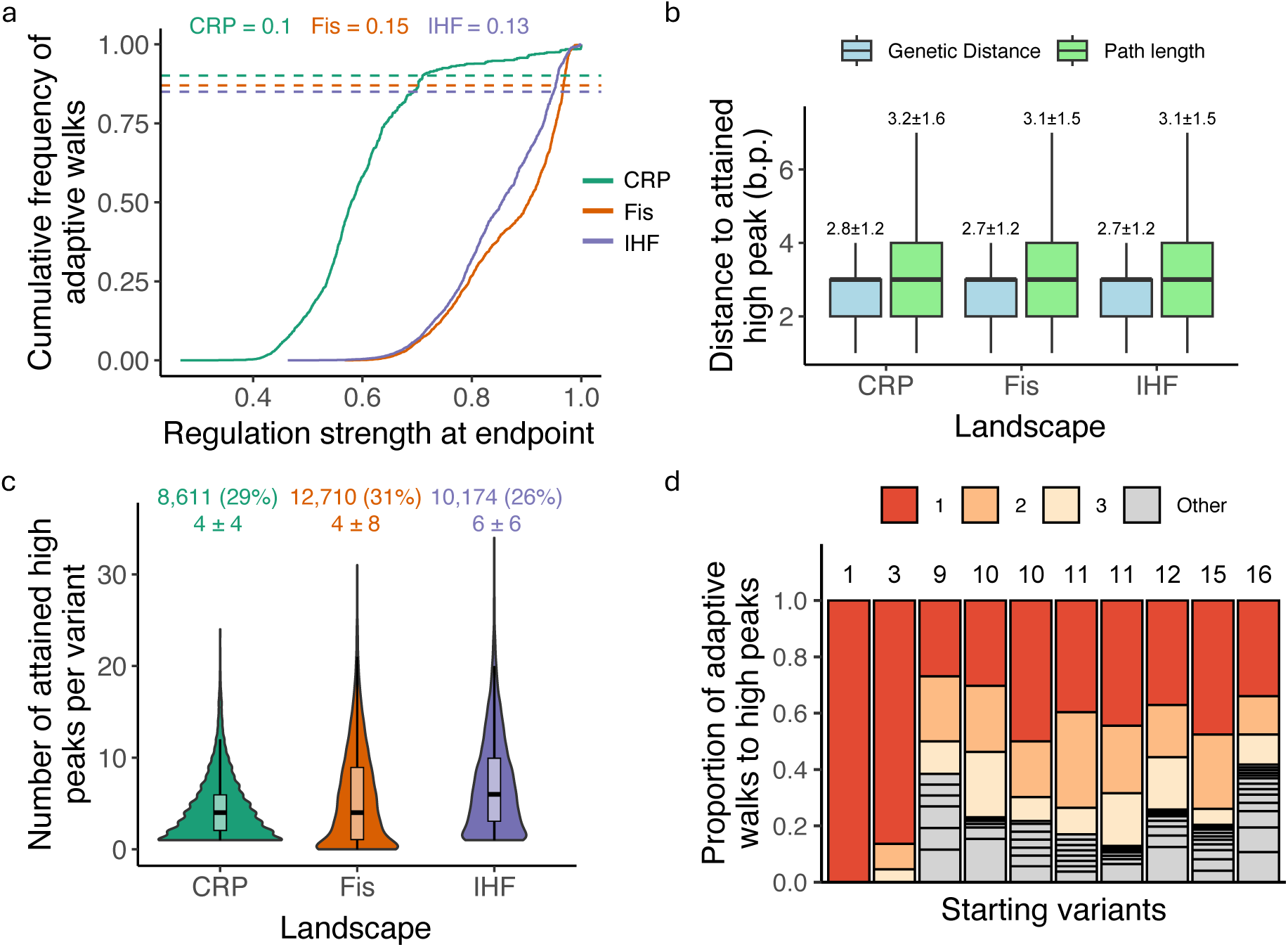
Peak accessibility, contingency and biases in adaptive walks. **a. More than 10% percent of adaptive walks lead to high peaks.** The graph shows the cumulative distribution function (CDF, vertical axis) of regulation strength (horizontal axis) attained at the end of 15 million adaptive walks (15,000 starting genotypes × 1,000 adaptive walks each) for each TF landscape (color legend, CRP: green; Fis: orange; IHF: purple). Dashed lines intersect the CDF at points equivalent to the wild-type regulation strength for each TF (color legend). They help to infer the proportion y_TF_ of walks that terminate at high peaks (S>S_WT_), which is indicated by the numerical values at the graph’s top for each of the landscapes (color legend). **b. Paths to high peaks are not much longer than genetic distances from the starting genotype.** Colored boxplots summarize, for each of the three landscapes (horizontal axis), the distribution of shortest genetic distances (blue) and actual path lengths (green) for adaptive walks terminating at high peaks. Path lengths are typically less than one mutational step longer than genetic distances. Each box covers the range between the first and third quartiles (IQR). The horizontal line within each box represents the median value, and whiskers span 1.5 times the IQR. Numbers atop each plot indicate means ± one standard deviation. **c. Different numbers of high peaks are reached from different starting variants.** Violin plots integrated with boxplots show the distribution in the number of high peaks reached (vertical axis) from each starting variant that attained any high peak during 1,000 adaptive walks.

To determine whether high regulatory peaks are more accessible due to chance alone, we compared the empirical landscapes to randomized “null” landscapes. Specifically, we generated these randomized landscapes by permuting regulation strengths across genotypes while preserving the sampled genotype network (**Supplementary Methods 7.6**). On each randomized landscape, we then performed adaptive-walks (15,000 walks for each of 10^3^ starting genotypes) with the same parameters as for the biological landscape. For all three landscapes, the fraction of adaptive walks reaching high regulatory peaks in the empirical landscapes exceeds that for the randomized landscapes by almost three-fold, a difference that is statistically significant (**Supplementary Figure S30**, **Supplementary Methods 9**). In sum, rugged regulatory landscapes strongly constrain evolutionary trajectories, yet do not render the evolution of strong regulation vanishingly rare. Instead, strong regulatory phenotypes remain evolutionarily attainable at levels that exceed null expectations, even though they are reached by only a minority of evolving populations.

The adaptive walks that reached a high peak were short (**Figure 4b**). On average, they were also no more than half a mutational step longer than the shortest genetic distance between each starting genotype and peak (**Figure 4b**, CRP landscape: path length 3.2 ± 1.6 [mean±s.d.] vs. genetic distance 2.8 ± 1.2; Fis landscape: path length 3.1 ± 1.5 vs. genetic distance 2.7 ± 1.2; IHF landscape: path length 3.1 ± 1.5 vs. genetic distance 2.7 ± 1.2).

These short paths can be explained by our previous observation that peaks are widely distributed across the genotype spaces of the three landscapes. This distribution makes it easier for populations starting from any (non-peak) genotype to find a local peak through few mutations. Additionally, path lengths realized during adaptive walks tend to be shorter than the average lengths of accessible paths. (**Figure 4b**).

Because genetic drift can cause evolving populations to escape a low fitness peak and attain a higher fitness peak, we also asked how small population sizes affect the likelihood that a population attains a high fitness peak (**Supplementary Material 8, Supplementary Figures S26-28**). When simulating adaptive evolution in populations of N=10^2^ individuals, we found that the likelihood that a population reaches a high peak increased by at least 10% percent (to 18%, 20%, and 21% for the CRP, Fis, and IHF landscapes, **Supplementary Figures S26-S28**).

We also studied how deviations from the strong selection weak mutation regime would affect evolutionary dynamics. In large populations or at high mutation rates, populations tend to be polymorphic and are subject to clonal interference, where the most advantageous of several co-occurring mutations typically dominates and becomes fixed ^105–107^. To approximate this dynamic, we conducted simulations using “greedy” adaptive walks (**Supplementary Methods 8**) ^25,108^. In such an adaptive walk, it is always a genotype’s mutational neighbor with the highest increase in regulation strength that becomes fixed ^100,108–110^. In other words, a greedy walk starting from a given genotype is deterministic. We, therefore, simulated only one walk for each of the 15,000 randomly chosen (non-peak) starting genotypes. We found that the fraction of greedy walks reaching high regulatory peaks is slightly lower than in the SSWM regime, with 1%, 2%, and 5% fewer walks achieving such peaks in the CRP, Fis, and IHF landscapes, respectively (**Supplementary Figure S29**). This outcome is expected, because greedy walks prioritize immediate fitness gains at the expense of the ability to discover higher fitness peaks^108^.

### The attainment of peaks is highly contingent on chance events

Because different basins of attraction share many TFBS genotypes (**Figure 3d, Supplementary Figures S24-S25**), it is possible that adaptive evolution starting from any one genotype can reach different peaks, depending on which mutational paths it takes. In other words, the structure of the landscapes we study may lead to evolutionary contingency ^111,112^ – the dependence of an outcome of evolution on unpredictable chance events^47,111,113^. Indeed, non-peak genotypes can access on average around 30% of the total number of high peaks in each landscape (**Supplementary Figure S25).** To assess the prevalence of evolutionary contingency, we determined how many different peaks are attained by 10^3^ adaptive walks starting from the same randomly chosen genotype. We performed this analysis on 15,000 starting genotypes, focusing on the subset of those starting genotypes from which at least one high peak is reached during the 10^3^ walks. Specifically, 28.6%, 23%, and 24.5% of starting genotypes for the CRP, Fis, and IHF landscapes, reached at least one high peak. Notably, these percentages represent the fraction of starting genotypes that can access at least one high peak, not the total fraction of adaptive walks that lead to a high peak, which is lower (as shown in **Figure 4a**). For instance, while adaptive walks starting from 28.6% of genotypes reached at least one high peak in the CRP landscape only 10% of all adaptive walks in this landscape end at a high peak (**Figure 4a**). Importantly, for any one starting genotype where random walks reached at least one high peak, they typically reached more than one high peak (**Figure 4c**, median±IQR, 4±4, 4±8, 6±6 for the CRP, Fis, and IHF landscapes, respectively). The number of high peaks attained from any one starting genotype was as large as 21, 30, and 33 for the three respective landscapes. Thus, evolutionary contingency is pervasive in all three landscapes. In addition, not only the end point of evolution but also the evolutionary trajectories leading to this end point show contingency. That is, adaptive walks starting from a specific genotype and ending at a specific high peak often take multiple paths to this peak (**Supplementary Figure S31**).

Biases towards specific evolutionary outcomes have been documented in both empirical and computational studies ^18,31,47,111,114–116^. They also exist in our landscapes (**Figure 4d, Supplementary Figure S31**). **Figure 4d** illustrates both contingency and bias for 10^3^ adaptive walks starting from each of 10 different genotypes in the CRP landscape (thus a total of 10^4^ adaptive walks). Depending on the starting variant, the adaptive walks reached between 1 and 16 high peaks. Whenever different peaks are reached, they are reached by markedly different proportions of adaptive walks. The same holds in the Fis and IHF landscapes (**Supplementary Figure S33**). Additionally, while multiple routes to each peak can be traversed (**Supplementary Figure S31**), we observed in all three landscapes biases towards specific paths that are taken more frequently than others (**Supplementary Figure S34**).

The number of these variants is shown as an integer on top of the panel, and its percentage among all 15,000 starting variants is shown in parentheses (color legend). Underneath it, the panel shows the mean ± 1 s.d. of the number of attained high peaks. **a. d. Some high peaks are reached more often than others.** We randomly and uniformly sampled 10 starting genotypes from the CRP landscape, started 10^3^ adaptive walks from each, and recorded the number and frequency of distinct high peaks attained in these random walks. Results for each starting genotype are symbolized by a vertical bar. The number of stacks within each bar (delineated by horizontal lines, also indicated by an integer above each bar) indicates the number of high peaks reached by the 10^3^ adaptive walks. Starting variants are ordered in ascending order based on this number of attained peaks. Stack height indicates the fraction of walks that reached the same peak, and is indicated in red, orange, and yellow for the three most frequently attained peaks. **See Supplementary Figure S33** for the Fis and IHF landscapes.

Lastly, because sort-seq measurements are subject to experimental uncertainty^71,117,118^, an important question is whether such noise affects the inferred structure of a regulatory landscape and the resulting evolutionary dynamics. To address this issue we explicitly incorporated empirically estimated measurement uncertainty into our fitness comparisons and repeated pertinent analyses under two conditions: a noise-free landscape (𝒢*^S^*) and an “uncertainty-aware” landscape 𝒢^S,τ^ using genotype-specific noise estimates (τ, see **Supplementary Figure S35**, **Supplementary Methods 10**). We examined whether noise-induced changes in landscape structure influence evolutionary trajectories. To do so, we compared adaptive walk dynamics between the two kinds of landscapes using identical population genetic parameters. Despite differences in peak counts and local topology, the overall pattern of genotype visitation during adaptive walks was highly similar between the two kinds landscapes. Specifically, the visitation frequency profiles of genotypes were strongly correlated between 𝒢^*S*^ and 𝒢^*S,τ*^ landscapes (Spearman’s ρ<0.001 shown in **Supplementary Figure S35**). This means that genotypes frequently accessed in the noise-free landscape remain frequently accessed in the noisy landscape.

## DISCUSSION

We evaluated the ability of three *E. coli* global regulators to control gene expression through each of more than 30,000 TFBSs for each regulator. To this end, we utilized a synthetic plasmid-based system that facilitates high-throughput fluorescence measurements. This system allowed us to quantify the regulation strength of individual TFBSs by measuring gene expression through GFP fluorescence ^74^. Additionally, the system insulates the control of transcription from any direct effects library sequences might have on transcription, such as the transcriptional and translational impacts of 5’ untranslated regions on gene expression ^119,120^. Recent studies have investigated large empirical datasets of cis-regulatory genotypes, examining the impact of sequence variation on gene regulation in both eukaryotes and prokaryotes^6,16,72,81,121–124^. However, few studies have focused on these interactions from an evolutionary perspective and studied adaptive landscapes of gene regulation, as we do here^6,16,123^.

We showed that the regulatory landscapes of all three TFs are highly rugged and have multiple peaks. The ruggedness of all three landscapes is also supported by the prevalence of epistasis between pairs of TFBS mutations (**Supplementary Table S5**). A particularly important form of epistasis is sign epistasis^24,93,94^, because it can lead to multiple adaptive peaks ^24,93,94^ (see **Supplementary Methods 7.5**). Our landscapes contain up to 65% of mutation pairs with sign epistasis, a value that is especially high compared to the almost exclusively additive interactions of mutations in eukaryotic TFs^6,125^. However, the TFs we study are not exceptions among other prokaryotic TFs. Recent smaller-scale studies have shown that epistatic interactions are common in binding sites of local prokaryotic regulators, such as AraC (sign epistasis in more than 50% of 20 mutants^126^) and Cl (sign epistasis in 85% of 113 mutants^127^).

A possible reason for this greater incidence of epistasis lies in the nature of prokaryotic TFBSs. Specifically, prokaryotic TFBSs are at approximately 20bps twice as long as eukaryotic TFBSs^80,128^ and exhibit symmetries that reflect the dimeric state of their cognate TFs^129–131^. These factors may increase the likelihood of intramolecular epistasis. Our observations raise important questions for future work, such as why the landscapes of prokaryotic TFBSs differ so dramatically from those of eukaryotic ones. And what do these differences imply for the evolutionary dynamics of gene regulation?

Despite the high ruggedness and pervasive epistasis of these landscapes, we found that a modest fraction of evolving populations can still access the highest regulatory peaks. Specifically, our evolutionary simulations show that 10% of populations with a size typical of *E. coli* reach one of the highest peaks. This percentage is significantly higher than in randomized landscapes (**Supplementary Methods 9**; **Supplementary Figure S30**), which shows that the structure of our regulatory landscapes facilitates access to stronger regulation than expected by chance. We speculate that this property reflects the biological role of global regulators, which coordinate the expression of many target genes^11^ and thus must operate across a wide spectrum of regulation strengths, while still permitting the evolution of strong regulation when necessary.

The clonal interference that may occur in even larger populations or in populations with high mutation rates reduces the accessibility of high peaks from 10% to 5% of evolving populations (**Supplementary Figure S29**). Conversely, in small populations where genetic drift can help a population escape from a low peak, this percentage increases to 18%. Numbers like these render the de novo evolution of strong transcription factor binding sites plausible for our three global regulators. Once such binding sites have originated in one population, they can also spread to others through horizontal gene transfer^132,133^.

In addition to increasing the ruggedness of a landscape, epistasis can also influence the evolution of populations on the landscape^24,94,134^. The regulation strength of a TFBS is partially determined by the combined interactions of the nucleotides within the TFBS^50,135–137^, which in turn affects TF-TFBS binding affinities. Different combinations of mutations can result in similar regulation strengths and create multiple evolutionary pathways to achieve optimal or near-optimal regulation^6,138^ (**Supplementary Figure S31**). This can lead to overlapping basins of attraction among different peak TFBSs conveying strong regulation, such that even evolution starting from the same genotypes can take different paths and reach different peaks with similar regulation strengths. Indeed, we observed such overlapping basins in our landscapes (**Figure 3d and Supplementary Figure S24**). Moreover, epistasis can create plateaus of regulation strength where various combinations of nucleotides yield TFBSs with intermediate regulation strengths. These plateaus further contribute to the overlapping basins of attraction we observe, and may serve as common evolutionary intermediates that multiple starting sequences can traverse on their way to different peaks.

In a landscape with substantial epistasis, the sequence of mutations that occurs in an evolving population can also render adaptive evolution highly contingent on this sequence^111,112^. For example, some beneficial mutations within a TFBS might only be accessible after other specific mutations have occurred. Different starting sequences or early mutations can also change the spectrum of accessible mutations, leading to different peaks with similarly strong regulation ^111,112^. Indeed, we observed in all three landscapes that different evolving populations starting from the same genotypes in the landscape attain different peaks (**Figures 3d, 4d and Supplementary Figure S33**) ^111,113,139,140^. Such contingency reduces the predictability of evolution^47,111,141^.

Despite the prevalence of contingency in all three landscapes, evolving populations starting from the same genotype more often attain some peaks than others (**Supplementary Figure S33)**. Moreover, for each attained peak with multiple possible evolutionary routes, we also observed that some paths are more frequently transversed than their alternatives (**Supplementary Figure S34)**. That is, evolution is biased towards traversing some paths and attaining some peaks more often than others^142,143^. These observations emphasize the complex interplay between chance, contingency, and evolutionary biases in shaping the outcomes of adaptive evolution.^47,111,113,115^.

The three TFs we studied here belong to different protein families, yet they have regulatory landscapes with similar topography. All three landscapes are highly rugged, highly epistatic, and harbor multiple peaks that are widely scattered, and this holds for peaks of low, intermediate, and high regulation strength. In addition, peak accessibility in all three landscapes increases with peak height (**Supplementary Figure S32**). On the one hand, these commonalities may be caused by common biological or biochemical properties of the three TFs. For example, CRP has been suggested to possess nucleoid-associated protein properties similar to Fis and IHF, due to its ability to bend and loop DNA^144^. On the other hand, the commonalities may reflect general characteristics of global gene regulation. One of them is that global TFs often bind unspecifically to multiple TFBSs^11,13^. Also, they may bind DNA with a broad range of different affinities centered around intermediate affinity, rather than bind few sites but very strongly ^13^. This possibility is supported by comprehensive analyses of in vitro eukaryotic TF binding affinities for thousands of TFBS variants, showing essentially continuous binding affinity distributions for hundreds of sites^6^.

Despite broad similarities among the three landscapes we study, we also found some differences. Most notable is the narrower distribution of regulation strengths for the binding sites of CRP compared to those of Fis and IHF (**Figure 2a**). This is not unexpected, given that CRP binding sites are known for being quasi-symmetric and less degenerate than those of Fis and IHF^57,60^. Previous studies have also shown that Fis and IHF binding sites are less conserved in their nucleotide sequence and more biased toward AT-rich content^43,145,146^. Differences in the distribution of regulation strengths may also result from differences in functions among the three proteins. CRP’s function is mostly restricted to gene regulation. As a result, it may have evolved to finely tune its regulatory output, resulting in a narrower distribution of regulation strengths. In contrast, Fis and IHF are involved in functions beyond gene regulation, such as DNA replication and genomic structural maintenance^147,148^, which might impose additional constraints or require a broader range of DNA binding strengths.

One limitation of our work stems from our rigorous quality filtering of sort-seq data. As a result, we lack regulatory data for some 40% of the TFBSs in each landscape. This limited diversity of reliable data is a common feature in mutational library studies. It has several technical causes, such as biases in library synthesis ^149^, PCR amplification ^150^, cloning, and loss of sequence diversity after cell sorting ^122^. In future work, this limitation could be overcome by a combination of strategies, such as subsampling complete genotype spaces or combining different molecular methods to overcome biases and diversity loss during PCR amplification^151,152^, sorting^71,117,153^, and high-throughput sequencing^123^.

Importantly, although undersampling of genotype space is a limitation, standard approaches such as random subsampling or predictive modeling are not straightforward remedies. Several of our core analyses—including peak identification, quantification of epistasis, and assessment of evolutionary accessibility—rely on combinatorially complete local neighborhoods in genotype space. Random subsampling of genotypes would remove mutational neighbors, and thereby confound pairwise comparisons and the interpretation of landscape topology. Predictive modeling could be used to infer missing genotypes and reconstruct more complete landscapes^154^, but it requires additional assumptions that introduce their own limitations. In addition, developing, validating, and benchmarking such models would be beyond the scope of this study, which is focused on empirical landscape mapping, and can serve as a starting point for future modeling work.

A second limitation comes from our use of the sort-seq method, which is best-suited for our work, because it allows the high-throughput measurement and sorting of millions of individual cells in a standardized and straightforward manner. However, the method’s accuracy depends on the binning procedure. Other studies have used between two and several dozen bins, with or without unbiased sampling ^71,73,81,117,121,122,155^. Recommendations emerging from this work include sorting of cells into at least four logarithmically (log_2_) equally-spaced bins, each covering approximately 12.5-15% of the fluorescence distribution, ^81,122,156^. We followed these recommendations, using 13 bins. In addition, we computed a weighted average to calculate expression values from sequences appearing in multiple bins, a straightforward method validated by robust studies in transcriptional regulation analysis ^16,72^. In addition, we validated individual regulation strengths with an independent method to demonstrate its reliability.

A third limitation of our study is that we only examined variation at eight positions for each TF. We selected these positions based on their importance for DNA binding and regulation by our TFs, as evidenced by their high information content. We cannot exclude the possibility that selecting a different set of positions could yield different landscape topographies. However, we speculate that less information-rich position would not reduce but rather increase the potential for the de novo evolution of strong TFBSs. For example, they may facilitate landscape navigability through extradimensional bypasses^96,157^ or provide small-incremental changes in regulation strengths. These small changes help populations reach peaks via diminishing returns effects, meaning that as a population gets closer to a peak, each subsequent mutation contributes progressively smaller improvements to regulation strength ^158,159^. Investigating a wider array of positions and larger regulatory landscapes remains an important task for future work.

Fourth, we use a simplified empirical system for the study of gene regulation—a promoter followed by a single TFBS. Although this regulatory architecture exists for some genes^160^, global regulators often form part of more complex, combinatorial architectures^4,160,161^ that are influenced by environmental factors^2,162,163^, concentrations of active TFs^164–166^, chromosome structure^167–171^, methylation states^172^, and more. Studies on more complex promoter architectures may reveal regulatory landscapes with different topographies.

Lastly, we also analyzed the adaptive evolution of TFBSs in a simplified manner, performing data-driven evolutionary simulations rather than experimental evolution. Although many studies assume that regulation strength and fitness are correlated, this is not always the case. Low binding affinities can be adaptive during development ^173^, and many genes exhibit a nonlinear fitness-expression function ^174^ with a plateau of maximal fitness across a wide range of expression levels ^175–177^. For instance, most mutations and polymorphisms in the promoter of the yeast gene TDH3 do not significantly affect fitness in a glucose-rich medium ^176^. For these reasons we focused our observations and interpretations on regulatory phenotypes (regulation strength). Our simplified approach allowed us to model population evolution in a large genotype space and avoid monitoring thousands of evolving regulatory sequences simultaneously in vivo^23,178^. However, it cannot replace experimental evolution of TFBSs, which also remains an important challenge for future work.

Transcription factor binding sites are among the simplest units of biological organization. Our work provides the first large-scale analysis of the regulatory landscapes formed by such sites for global transcriptional regulators. It shows that strong binding sites can readily evolve de novo, even though prokaryotic transcription factor binding sites are much larger than their eukaryotic counterparts and have more rugged regulatory landscapes. In addition, the evolution of these simple sequences also displays phenomena that have been characterized in much more complex systems, such as evolutionary contingency and evolutionary biases.

## MATERIALS AND METHODS

The Supplementary Materials contain extended details of experimental procedures and data analysis.

### Strains and plasmids

Bacterial strains and plasmids used in this work are listed in Table 1. We obtained electrocompetent *E. coli* cells of strain SIG10-MAX^®^ from Sigma Aldrich (CMC0004). We used this strain for molecular cloning and library generation due to its high transformation efficiency. The genotype of this strain (**Supplementary Table S4**) is similar to DH5α (Sigma Aldrich commercial information, see **Supplementary Table S4**). The strain is resistant to the antibiotic streptomycin.

We amplified plasmid libraries in SIG10-MAX^®^, extracted their DNA, and transformed them into mutants derived from *E. coli* K-12 strain BW25113 that harbor chromosomal deletions of the *crp*, *fis*, or *ihfa* gene. We obtained these mutant strains from the KEIO collection ^179^ and used them for sort-seq experiments. The design, genetic parts, and assembly of the plasmid vectors we used in this study are available in the Supplementary material, as are all primers, TFBS sequences/libraries, strains, and plasmids.

### Sort-Seq procedure

To explore the regulatory effects of each transcription factor on binding sites in the corresponding library, we constructed three plasmids, each of which enables the inducible expression of one of our three TFs. These are plasmids pCAW-Sort-Seq-V2-CRP, pCAW-Sort-Seq-V2-Fis, and pCAW-Sort-Seq-V2-IHF (**Supplementary Methods 3** and **Supplementary Table S1**). We cloned the TFBS libraries (**Supplementary Table S3**) into their respective plasmids and then transformed them into mutant strains lacking the corresponding TFs (*Δcrp*, *Δfis*, and *Δihf*, as listed in **Supplementary Table S2**). We induced TF expression using anhydrotetracycline (Atc) and, after overnight growth, performed cell sorting for cell populations. During sorting, we distributed cells into 13 equally-spaced logarithmic bins based on their fluorescence levels. We replicated each sort-seq experiment three times from three separate library transformations for each TF. To mitigate the impact of extrinsic noise (gene expression variation among cells ^180^), we adhered to standard protocols by normalizing our GFP fluorescence measurements against mScarlet-I fluorescence values obtained from flow-cytometry assays ^7,71,181,182^. We subsequently recovered the sorted cells from each bin in 50mL Falcon tubes containing 10 mL of LB medium supplemented with chloramphenicol and incubated them overnight at 37°C with shaking at 220 rpm. After this growth period, we re-sorted cell cultures from each bin to eliminate potential contaminants and ensure that the cell populations had preserved their fluorescence distributions. Following re-sorting, we extracted plasmids from the cell population of each bin. We amplified and barcoded the TFBS region from each population through a polymerase chain reaction (PCR). Lastly, we sequenced barcoded amplicons containing TFBS sequences, and used the sequencing results to calculate the regulation strengths of each TF to its TFBSs in the corresponding library. More details are provided in the Supplementary Material.

### Regulation strengths

Due to gene expression and measurement noise, individual TFBS variants in a sort-seq experiment usually appear in more than a single bin, and their read count (frequency) varies among bins^71,72,121,153^. Following established practice^18,72,123^, we used a weighted average of these frequencies for each variant to represent the mean expression level caused by the variant. To facilitate the interpretation of this quantity, we converted this expression level into a regulation strength relative to the highest observed regulation strength for a given TF, to which we assigned a value of one. From each library, we selected a single naturally occurring binding site for each TF that was previously characterized in the literature as a strong binder. We called this TFBS the WT sequence and used it as a baseline to separate weakly (higher GFP expression) from strongly (lower GFP expression) regulating TFBSs.

### Validating regulation strengths with plate reader measurements

To further validate our regulation strength data, we chose 10 DNA binding sites from each bin (10 variants ×13 bins = 130 variants in total per TF library, **Supplementary File 2**), covering a wide range of measured regulation strengths. We cloned these sequences into the appropriate vector pCAW-Sort-Seq-V2-TF (TF: CRP, Fis or IHF), and transformed them into the appropriate mutant strain. We picked individual colonies and grew them overnight (16 hours, 37°C, 220 rpm) in liquid LB supplemented with 50 μg/mL of chloramphenicol and anhydrotetracycline. We diluted the cultures to 1:10 (v/v) in cold Dulbecco’s PBS (Sigma-Aldrich #D8537) to a final volume of 1 mL. We transferred 200 μl of the diluted cultures into individual wells in 96-well plates and measured GFP fluorescence (emission: 485nm/excitation: 510nm, bandpass: 20nm, gain: 50), as well as the optical density at 600nm (OD_600_) as an indicator of cell density. We then normalized fluorescence by the measured OD_600_ value to account for differences in cell density among cultures and compared the obtained ratios to the previously inferred regulation strengths for the 130 selected variants. We performed all such measurements in biological and technical triplicates (three colonies per sample, and three wells per colony, respectively).

## Code Availability and Data Analysis

All code used for processing data and plotting, as well as the final processed data, plasmid sequences, and primer sequences are available in our GitHub repository.

**Supplementary Information** is linked to the online version of the paper

## Acknowledgements

We gratefully acknowledge financial support from the Swiss National Science Foundation grant 310030_208174. We also extend our thanks to the UZH University Priority Research Program in Evolutionary Biology, the UZH flow cytometry facility, and the Functional Genomics Center Zurich for their technical support. We are especially thankful to Andrei Papkou for his guidance with computational analysis and for engaging in theoretical discussions.

## Author contributions

C.A.W. and A.W conceived the study and designed experiments. C.A.W. carried out experiments. C.A.W and A.W. analyzed data. C.A.W wrote computer code to carry out bioinformatic work and analysis. C.A.W generated figures. L.G. wrote computer code to carry out simulations. C.A.W. and A.W. wrote the paper, which was edited by all authors.

## Data availability

Sequencing data has been deposited in the NCBI database under the BioProject accession code: PRJNA1162449. The data generated in this study have been deposited in the Zenodo public repository and are accessible via the following DOI: 10.5281/zenodo.13838265

## Computer code

The computer code generated in this study has been deposited in the Zenodo public repository and is accessible via the following DOI: 10.5281/zenodo.13838265

## SUPPLEMENTARY METHODS

### 1. General procedures

Although all general procedures have been previously described ^18^, we describe them again below for completeness.

#### 1.1 Media and reagents

To prepare SOB medium, we mixed 25.5g of solid medium stock (VWR J906) with 960 ml of purified water and subjected the resulting suspension to autoclaving. To prepare the SOC medium, we dissolved 20 ml of 1 M D-glucose (Sigma G8270) and 20 ml of 1 M magnesium sulfate (Sigma 230391) in 960 ml of pre-prepared SOB solution. For the LB medium, we combined 25g of solid medium stock (Sigma-Aldrich L3522) with 1 liter of purified water and then autoclaved it. To prepare the M9 minimal medium, we diluted M9 minimal salt sourced from Sigma (M6030) in distilled water as per the manufacturer’s guidelines, autoclaved the solution, and added 0.4% glucose (Sigma G8270), 0.2% casamino acid (Merk Millipore, 2240), 2 mM magnesium sulfate (Sigma 230391), and 0.1 mM calcium chloride (Sigma C7902). Where required, we supplemented growth media with chloramphenicol (50 µg/mL), anhydrotetracycline (100 ng/mL, Cayman-chemicals #10009542), and/or glucose (0.4% w/v final concentration). We prepared anhydrotetracycline by diluting the dried chemical in absolute ethanol, from a stock concentration (1000X) of 100 µg/mL to a working concentration of 100 ng/mL.

#### 1.2 Overnight incubation of cultures in liquid and solid medium

We cultivated bacteria in liquid LB medium (using either 15mL or 50mL Falcon tubes), enriched with chloramphenicol at a concentration of 50 µg/mL. We incubated these cultures for a period of 16 hours at a temperature of 37°C, with a shaking speed of 200rpm and 50 mm orbital motion, using an Infors HT Multitron Incubator Shaker. Similarly, for cultures in solid medium, we grew bacterial colonies on LB-agar plates (using sterile plastic Petri dishes of 90mm × 15mm dimensions), also supplemented with chloramphenicol at a concentration of 50 µg/mL, and incubated them for the same time and at the same temperature.

#### 1.3 PCR Reactions

Except where specifically mentioned, we amplified DNA fragments through a polymerase chain reactions (PCR), employing Q5^®^ high-fidelity polymerase (NEB #M0491L) to minimize mutation introduction into the amplicons. We followed the protocol recommended by NEB, aiming for a final reaction mixture of 50uL. We conducted each PCR reaction twice, and combined the products from these duplicates after the completion of the reaction. We determined the primer melting temperatures (Tm) using the NEB Tm calculator (accessible at https://tmcalculator.neb.com/#!/main), with primers at a concentration of 500nM.

#### 1.4 Verifying PCR products through gel electrophoresis

Unless indicated otherwise, we verified successful PCR amplification via gel electrophoresis to ensure the presence of singular-band amplicons and the absence of non-specific bands. This process involved separating PCR products in a 0.8% agarose Tris-EDTA (TAE) gel. We conducted electrophoresis for 45 minutes at 120V, or until the bands had progressed beyond the halfway point of the gel’s total length.

#### 1.5 DNA purification with commercial kits

Once we had confirmed a successful PCR through gel electrophoresis, we purified the PCR products utilizing the Monarch^®^ DNA PCR/Gel Extraction Kit (NEB #T1020L), adhering to the manufacturer’s protocol. In situations requiring further purification (such as the occurrence of non-specific bands post-PCR), we performed gel purification. This involved mixing 10 μL of 6x NEB DNA dye with each 50 μL PCR product, then loading all 60 μL onto a 1% agarose gel. We carried out electrophoresis for 45 minutes at 120V, or until the bands had moved more than halfway through the gel. We then excised the DNA band corresponding to the amplified sequence using a scalpel. For extracting DNA from the gel, we used the Monarch^®^ DNA Gel Extraction Kit (NEB #T1020L).

#### 1.6 Gibson assembly^183^

We assembled PCR-amplified fragments using the NEBuilder-HiFi^®^ DNA Assembly Master Mix kit (NEB #E2621L). We determined the molarity required for assembly based on the guidelines outlined by the Barrick Lab (details available at https://barricklab.org/twiki/bin/view/Lab/ProtocolsGibsonCloning). We incubated the assembly mix for one hour at 50°C in a dry bath incubator, followed by cooling on ice for subsequent steps.

#### 1.7 Preparation of electrocompetent cells

We utilized glycerol/mannitol step centrifugation to prepare electrocompetent cells ^184^. To this end, we first cultured the appropriate *E. coli* strain (**Supplementary Table S4**) in 5 mL SOB medium at 37°C with a shaking speed of 250 rpm overnight. The next day, we transferred 3 mL of the culture to 300 mL SOB medium, and incubated under the same conditions until the OD600 reached a value between 0.4 and 0.6 (measured at an optical path length of 1 cm), which took approximately 2-4 hours. After cooling the culture on ice for 15 minutes, we centrifuged the cells at 4°C and 1,500 g for 15 minutes. We then resuspended the cells in 60 mL of ice-cold distilled water (dH_2_O) and divided them into three 50 mL tubes. Gradually, we added 10 mL of an ice-cold glycerol/mannitol solution (consisting of 20% glycerol (w/v) and 1.5% mannitol (w/v)) to each tube using a 10 mL pipette. We centrifuged the tubes at 1,500 g and 4°C for 15 minutes in an Eppendorf 5810/5810 R centrifuge with acceleration/deceleration set to zero. After discarding the supernatant, we resuspended cells in 3.0 mL of the same glycerol/mannitol solution. We transferred these suspensions to pre-cooled 1.5 mL tubes, and incubated them in a dry ice-ethanol bath for about 1 minute. Finally, we stored the suspensions at -80°C for future transformation experiments.

#### 1.8 Electroporation

In all transformation experiments described in this study, we used 100 µL of electrocompetent cells for electroporation, employing 0.2 cm cuvettes (EP202, Cell Projects, UK) and a Micropulser electroporator (Bio-Rad) set to the EC3 setting (15k V/cm). Post-electroporation, we recovered cells in 1 mL of SOC media, warmed in advance in 15 mL Falcon tubes. This recovery step lasted for 1.5 hours at 37°C with a shaking speed of 220 rpm. Unless specified otherwise, we spread 300 µL of each recovered culture on an LB agar plate that contained 50 μg/mL chloramphenicol. We then incubated this plate overnight for 16 hours at 37 °C. Subsequently, we confirmed the identity of the clones on the plate via Sanger sequencing.

### 2. The design of plasmid pCAW-Sort-Seq-V2

The plasmid pCAW-Sort-Seq-V2 is a derivative of the plasmid pCAW-Sort-Seq^18^. Briefly, plasmid pCAW-Sort-Seq harbours a pBBR1 replication origin, which ensures a broad host range and a low copy number—typically between 5 to 10 copies per cell^185^.It also harbours a chloramphenicol resistance gene, a TetR repression system^186^ and a TFBS measuring module. The latter consists of an interchangeable TFBS that lies between a constitutive promoter and a superfolder GFP (*sfgfp*^187^) reporter gene. In this system, TF-TFBS interactions can decrease GFP production by physically obstructing the bacterial RNA polymerase, a phenomenon known as steric hindrance. The reporter gene *sfgfp* is insulated by a transcriptional insulator named RiboJ, a synthetic ribozyme that removes 5’UTR interferences from variable TFBS sequences in the mRNA by self-cleavage^188^. More details and features have been previously described in ref. ^18^. Here, we have integrated a bicistronic expression cassette into pCAW-Sort-Seq to enable the regulated expression of a TF of choice. This cassette harbors the gene encoding one of our focal TFs situated upstream of a *mscarlet-I* ^189^ reporter gene to monitor the expression of this bicistronic operon via fluorescence. Regulation is achieved through the *pLtetO-1* ^186^ promoter, a synthetic promoter tightly repressed by TetR. Expression is initiated by the addition of anhydrotetracycline (Cayman Chemicals, catalog #10009542), which relieves TetR-mediated repression and activates the expression of the entire cassette.

### 3. Construction of the plasmid pCAW-Sort-Seq-V2 and its variants

We initiated the construction of pCAW-Sort-Seq-V2 by synthesizing (Twist Biosciences, Calfornia, USA) and cloning a codon-optimized *mscarlet-I* gene expression cassette into the pCAW-Sort-Seq plasmid. This gene is oriented antiparallel to the tetracycline resistance gene (*tetr*). This cloning step resulted in the formation of the intermediate pCAW-Sort-Seq-V2t plasmid (**Supplementary Figure S1)**. Subsequently, we amplified for each of our three TFs the encoding gene (*crp*, *fis* and *ihf*) from the genome of the *E. coli* strain BW25113 and inserted it upstream of the *mscarlet-I* gene in a bicistronic operon configuration using Gibson assembly. This process produced TF-specific expression plasmid variants (**Supplementary Figure S2)**. The final phase involved removing the core promoter to create negative controls for each TF and cloning libraries into each plasmid variant upstream of the sf*gfp* gene. This yielded the plasmids used in our experiments (**Supplementary Table S1, Supplementary Figure S2**). The specific sets of primers used for each PCR reaction are listed in **Supplementary Table S6**.

### 4. Library design, synthesis, and cloning

We based the design of the TFBS library for each TF on consensus sequences available in the RegulonDB database^70^, the most comprehensive database for *E. coli* transcriptional studies. We designed each library by randomizing the eight most important TFBS positions, such that each of the four nucleotides had an equal probability (0.25) to occur at each position. The expected library size for this approach is (4^8^) ^=^ 65,536 sequences. Library compositions can be found in **Supplementary Table S3**. We designed libraries in the Snapgene^®^ software (snapgene.com), and had them synthesized by IDT (Coralville, USA) as single-stranded DNA Ultramers^®^ of 140bp (4nmol). We resuspended each library in nuclease-free distilled water and serially diluted it to a concentration of 50ng/uL. We used Ultramers^®^ as templates in a PCR reaction for the formation of dsDNA fragments and library amplification. We amplified the Ultramer^®^ DNA molecules with the following PCR program: 98°C/30 s; 25 cycles of 98°C/10 s, 60°C/15 s and 72°C/80 s; and 1 cycle of 72°C/5 min. We opted for a maximum of 20 cycles to reduce amplification biases. We analyzed amplification products through gel electrophoresis to confirm that only a single product band with the expected size of around 140bp was present for each PCR reaction. After confirming the presence of single bands, we purified the products using the Monarch^®^ DNA gel extraction kit (NEB #T1020L). Whenever unspecific bands appeared during electrophoresis, we gel-purified the PCR products using the same kit.

For the construction of each plasmid-based library, we digested 1μg of the purified library with HindIII-HF (NEB #R3104) and BamHI (NEB #R3136) restriction enzymes in a 100μL reaction, followed by overnight incubation at 37°C. We isolated the appropriate cloning plasmid from its host strain using the QIAprep spin miniprep kit (Qiagen, Germany), and digested it with the same enzymes. Post-digestion, we added 3μL of Quick CIP (calf intestinal alkaline phosphatase) to prevent self-ligation by dephosphorylating DNA ends. We purified the ligated DNA using the Monarch^®^ DNA gel extraction kit (NEB #T1020L).

We performed the ligation with a 10:1 molar ratio of insert-to-vector, using 100ng of vector, 10 units of T4 DNA ligase (NEB #M0202L), and 2 µL of 10X ligation buffer in a 20 µL reaction. We incubated the mixture at 20-22°C for approximately 16 hours, followed by 15-minute inactivation of the ligase at 65°C. We purified the ligation product, resulting in 10 µL of DNA resuspended in dH_2_O. We transformed *E. coli* SIG10-MAX® cells by electroporation with this purified product.

Post-transformation, we plated 50 µL of each recovered culture on LB agar for colony-forming unit counting (cfu) and transformation efficiency estimation. From the agar plates, we selected 30 colonies for colony PCR and Sanger sequencing to assess library diversity (NightSeq^®^ service, Microsynth, Switzerland). Our mean transformation efficiency was 10^6^ cells per transformation. We diluted the remaining 950 µL of transformants in 9 mL of LB medium with chloramphenicol and cultured it overnight. We then aliquoted the overnight culture into 1 mL cryotubes with 20% glycerol and stored at -80°C.

After transforming plasmid libraries into the SIG10-MAX® strains for reasons of transformation efficiency and plasmid maintenance, we extracted the plasmids using a QIAprep spin miniprep kit (Qiagen, Germany), and transformed them into the appropriate host mutant host strains ***Δ****crp, **Δ**fis and **Δ**ihf* (see the strain genotypes in **Supplementary Table S2**). We cultured each library-transformed strain overnight, aliquoted it in 1 mL cryotubes with 20% glycerol, and stored it at -80°C for further experimentation.

### 5. Analysing and sorting cells

In preparation for cell sorting, we cultivated cells harboring each library in liquid LB medium enriched with chloramphenicol. Specifically, we cultured 1mL of transformed cell aliquots and a streak of cells with a control plasmid (a promoterless pCAW-Sort-Seq-V2 plasmid lacking sfGFP expression) in 50ml Falcon tubes with 9mL LB medium (containing 50 µg/mL chloramphenicol) overnight. Subsequently, we diluted these overnight cultures at a 1:100 ratio (v/v) in LB medium with chloramphenicol and divided them into two aliquots. We supplemented one aliquot with the anhydrotetracycline (Atc) inducer to express the plasmid-encoded TF. The other aliquot remained unchanged. We incubated both cultures for 5 hours until late-exponential/early-stationary phase (200RPM, 37°C). Subsequently, we diluted 20µL of the cultures in 1mL of cold filtered Dulbecco’s PBS (Sigma-Aldrich #D8537) in 15 mL FACS tubes.

We performed FACS-sorting on a FACS Aria III flow cytometer (BD Biosciences, San Jose, CA) using a 70 µm nozzle. We utilized a 488 nm laser for detecting forward scatter (FSC) and side scatter (SSC) with a 488nm/10nm band-pass filter. We set the flow rate to 1.0, adjusting sample dilution as necessary to achieve no more than ≈10,000 events/second. Considering the small size of bacterial cells, we reduced the particle detection threshold to the lowest feasible setting (200 arbitrary units on FSC and SSC channels), increasing it to a maximum of 500 units if background noise was excessive. We then adjusted FSC-H and SSC-H for cells with the negative control plasmid to center the bacterial population in the cytometer’s software (BD FACSDiva™ Software v9.0). We based the sorting and binning of cells on both mScarlet-I (PE-Texas-Red channel, excitation laser: YellowGreen 561nm, LP filter: 600 nm, BP filter: 610/20nm) and sfGFP fluorescence (FITC channel, excitation laser: 488nm, LP filter: 502nm, BP filter: 530nm/30nm), setting both the PE-Texas-Red and FITC channels voltages so that the median fluorescence of the negative control was between 0 and 100 (arbitrary units) on the FITC-H and PE-Texas-Red-H axes.

Initially, the sorting process involved establishing a gate for identifying cells that are red-fluorescence-positive, i.e., reporter and thus TF-expressing. To establish this gate, we began by measuring the autofluorescence of our negative control culture on the PE-Texas-Red-H axis. Thereafter, we examined the fluorescence of a positive control that had been induced with Atc and was expressing the mScarlet-I protein. We then configured a gate around the positive mScarlet-I-expressing population, ensuring that all cells sorted through this gate exhibited a level of reporter expression higher than the negative control.

Next, we established the green-fluorescent sorting gates. For setting sorting gates on the FITC-H axis, we first recorded the autofluorescence of the negative control culture. This median autofluorescence defined the upper boundary of the lowest bin (B1) for the experimental population. We then analyzed the fluorescence of 10^6^ cells expressing sfGFP and containing the library, without sorting, in order to establish the boundaries of the next binning gates. We took the lower bound of the highest bin (B13) for the experimental population to correspond to the 95th percentile of the fluorescence distribution of this population. We chose boundaries between the remaining 11 intermediate bins with equidistant spacing on a binary logarithmic (log_2_) scale. After we had determined the gates in this way, we calculated the fractions of the previously 10^6^ recorded cells that fell inside each bin.

To maintain statistical robustness and minimize sampling error in our downstream analyses, we wanted to ensure that each of the 65,536 unique sequences in our library was represented by at least 30 cells after sorting. In addition, it was crucial to account for an anticipated diversity loss of up to 70% of the cell population during sorting, due to factors such as cell death and the dilution of low-frequency and lower-fitness genotypes during the post-sorting recovery phase^122^. To establish the number of cells required to be sorted initially, we thus determined a multiplicative factor 𝑓 based on the minimally needed number of 30 cells per sequence and the expected cell retention rate post-sorting of 30%, i.e., 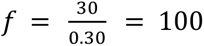 cells/sequence. We thus multiplied our initial library size by 100 for sorting purposes, i.e., we sorted total of 6,553,600 cells. This calculation aims to ensure that despite a substantial reduction in cell numbers post-sorting, each unique sequence remains adequately represented in the surviving cell population. We sorted cells into 1.5 mL Eppendorf tubes, each containing 500 µL of LB medium, and kept at 4 °C to prevent growth during sorting and sample processing. We replicated the entire sorting procedure three times, each based on independent library transformations.

After sorting, we added 1mL of LB without antibiotics to each tube and removed 20 uL of the resulting volume for serial dilutions. We transferred the remaining liquid culture (980 uL) to 50 mL falcon tubes and allowed each culture to recover for 2 hours (37°C, 220 rpm). After recovery, we added 9mL of LB supplemented with chloramphenicol, and grew the cultures overnight for freezing part of them in glycerol stock aliquots, for validating our binning procedure (see below), and for extracting plasmid DNA for subsequent PCR and sequencing steps. We used the 20 uL initially removed from each 1mL culture for preparing two serial dilutions (10^-4^ and 10^-6^) that we plated on LB-Cm agar plates (200uL per plate) to estimate the post-sorting viability through colony forming unit (cfu) counting. By knowing how many cells were sorted into each bin, we can estimate how many cells would be expected in our dilutions and compare this number with the cfu counts we observed. In this way, we estimated that on average (across bins) 77% of cells remained viable (standard deviation: 18.15%). We also estimated the genetic diversity of the library through Sanger sequencing of DNA from individual colonies (NightSeq^®^ service, Microsynth, Switzerland).

To validate our binning procedure, we re-grew binned cultures from either overnight recovered cultures or frozen aliquot stocks. We then measured their expression distributions by flow cytometry, which reproduced the original expression measurements. This approach allowed us to compare the fluorescence distributions of the re-grown cultures with the pre-sorting distributions. Specifically, we assessed whether the geometric mean of the fluorescence distribution for each sorting bin matched those recorded during the initial sorting procedure. Next, we re-sorted cells derived from each bin in order to eliminate cell cross-contaminations and other factors that could alter the bin distributions. The geometric mean is often preferred over the arithmetic mean in this type of analysis, because it is less affected by the presence of outliers in the data ^190,191^. It is also a more accurate representation of the central tendency of data that is log-normally distributed, which is often the case with flow cytometry fluorescence measurements^190,191^.

### 6. DNA extraction and sequencing

We diluted 500 uL of individual glycerol stocks of each replicate subpopulation of sorted cells (i.e., cells from each bin of fluorescence intensity) in 5mL of LB supplemented with chloramphenicol in 15mL Falcon tubes, and grew the resulting cell culture overnight (16 hours, 37°C, 220 rpm). On the next day, we isolated plasmids from each culture using a QIAprep^®^ spin miniprep kit (Qiagen, Germany). In order to allow the sequencing of multiple pooled samples (multiplexing), we barcoded our regulatory region through PCR with specific HPLC-purified primers (**Supplementary Table S6**) provided by Eurofins (Konstanz, Germany). We added barcodes to the 5’region of the amplicon through a PCR reaction. We performed this PCR reaction with the Q5 high-fidelity polymerase, and did so in triplicate for each miniprep-isolated plasmid library. To calculate primer melting temperatures (Tm), we used the NEB Tm calculator (https://tmcalculator.neb.com/#!/main) for a primer concentration of 500nM. We performed the PCR with the following program: 98°C/30 s; 25 cycles of 98°C/10 s, 64°C/30 s and 72°C/30 s; and 1 cycle of 72°C/2 min.

After PCR amplification, we digested the reaction products with the restriction enzymes DpnI (NEB #R0176L) and Exonuclease I (NEB #M0293L) in order to remove traces of genomic DNA, plasmids, and single-stranded DNA that could interfere with sequencing. The Master Mix we used for a single digestion harbored 1µL of 10x CutSmart^®^ Buffer (NEB #B6004S), 1µL of Exonuclease I (NEB #M0293L), 1µL of DpnI restriction enzyme (NEB #R0176L), and 7µL of distilled nuclease-free water. For each PCR product, we added 10µL of the Master Mix. We incubated the reaction for 1 hour at 37°C, following 15 minutes at 80°C for deactivation of the enzymes.

We purified the digestion products using the Monarch^®^ DNA PCR/Gel Extraction Kit (NEB #T1020L) and fractionated them through gel electrophoresis to confirm that only a single band with a size ≈150bp was present. After this confirmation, we pooled the purified PCR products from the different bins of each replicate sorting equimolarly to a total mass of 2,600 ng and a volume of 100 µL (26ng/µL of DNA) in 1.5mL Eppendorf tubes. We then sent the pooled purified amplicons for adapter ligation and sequencing at Eurofins (NGSelect Amplicons^®^ on Illumina HiSeq), obtaining 15 million paired-end reads (2 × 150 bp) for all samples.

We quantified the total number of reads (*t*) required for sequencing based on several factors, including the number of bins (*b)*, biological replicates (*r)*, strains (*s)*, genotypes (*g)*, and the reads per genotype (*p)*. Using these variables, we first calculated the total number 𝑛_’(_ of needed barcodes as follows:

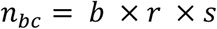

For each of our three TFs, the values of these parameters are *b*=13 bins, *r*=3 biological replicates, and *s*=1 strains, yielding *n_bc_*=39. The total number of required sequence reads can be estimated through the equation:

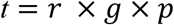

where we aimed for *p*=30 paired-reads per genotype. The number *g* of expected genotypes per library is the same (65,536) for each TF. These values lead to a total of *t* = 3 × 65,536 × 30 = 7,667,712 required paired-end reads for each library.

### 7. Data analysis

#### 7.1 Filtering and preparing sequencing reads

We processed the sequencing data with a blend of custom python and awk scripts, complemented by established bioinformatics utilities. Firstly, we trimmed sequences by computationally excising Illumina adapters, followed by merging paired-end reads. We then organized the paired-end reads into distinct files, each tagged with a sequence barcode identifying the sequence bin from which the corresponding reads originated.

For the removal of Illumina adapter sequences from the paired-end reads, we employed Cutadapt ^192^ with the following parameters:

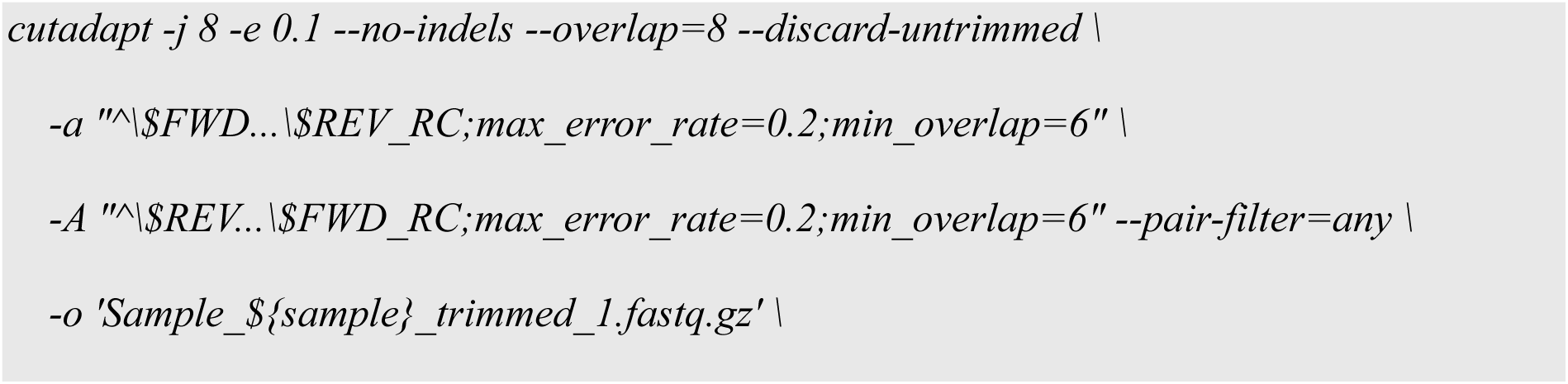
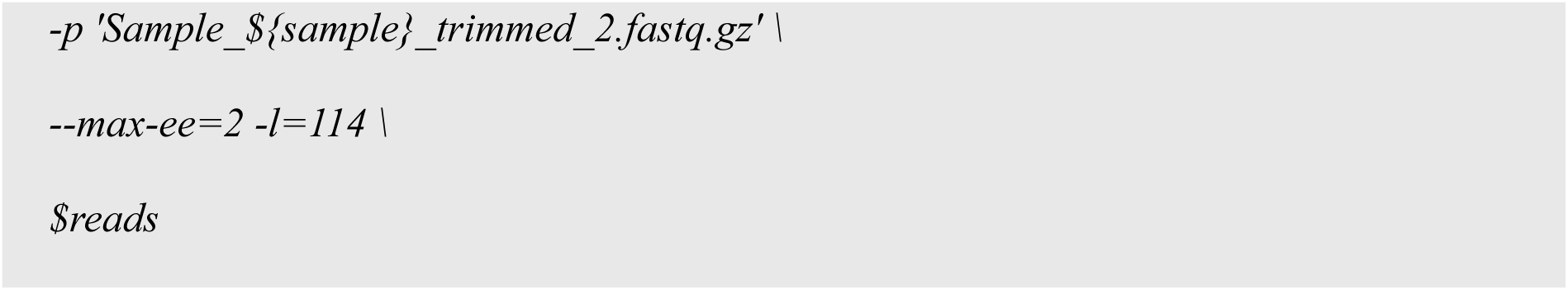

Here, “ADAPTER_FWD” and “ADAPTER_REV_RC” are placeholders for the actual adapter sequences used, which are 5’-AGATCGGAAGAGCACACGTCTGAACTCCAGTCA-3’ for read 1, and 5’-AGATCGGAAGAGCGTCGTGTAGGGAAAGAGTGT-3’ for read 2.

Because our amplicons were short (153 bp) and each sequencing sample was divided into two files with overlapping single-end reads, merging the two files (single-end reads) was necessary, for which we used the FLASH software ^193^. After the filtering and merging of reads, we demultiplexed paired-end reads with the following FLASH parameters:

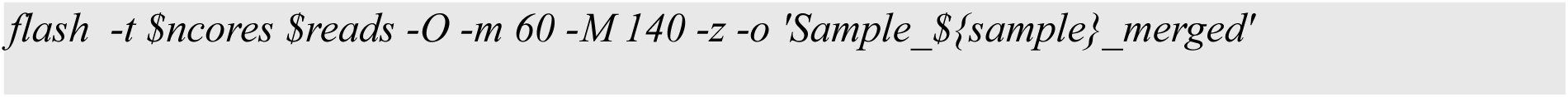

Next, we utilized the FastX toolkit (http://hannonlab.cshl.edu/fastx_toolkit/)^194^ to discern and preserve only those sequences that exceeded a high-quality threshold of Q = 33. Subsequently, we refined our dataset by removing sequences that contained undesirable mutations or insertions/deletions (indels) within the region extending from the start of the core promoter to the end of the variable TFBS library. We performed this filtering step using a custom R ^195^ script. Thereafter, we used custom awk and R^195^ scripts to convert the data into a table. Each row of this table contains data from one TFBS in the library, and each column contains the number of reads for this TFBS from one of the thirteen bins into which we had sorted cells. We estimated the average regulation strength of each TFBS from this data.

#### 7.2 Calculating regulation strengths

We calculated regulation strengths as previously described^18^. Briefly, in sort-seq experiments, sequences often appear in multiple fluorescence bins due to random mis-sorting^71,117^ and the stochastic nature of gene expression^180^. Following methods established in previous research ^16,72,123^, we determined the reporter expression level driven from each TFBS variant by calculating a weighted average of bins in which the variant occurred. This involved multiplying the frequency of each sequence (*x_i_*) in a given bin *i* by a numerical value representing that bin (*w_i_*=1,2,3,…,13), and then averaging these products over the total sequence count. Mathematically, this weighted average calculates as

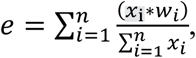

where *e* is the expression level driven by the sequence.

This approach yields a continuous spectrum of expression levels *e* within the range of 1 to 13. High expression values indicate robust GFP expression and, thus, weak binding of a TF to a TFBS variant. To facilitate interpretation, we inverted this scale so that higher values indicate stronger repression and lower GFP expression, i.e., we define the regulation strength *b* (strength of regulation) as follows:

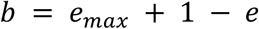

where *e_max_*=13 is the maximal expression level (maximal bin value). We then normalized *b* by the maximal regulation strength among all TFBSs we studied for a given TF. The resulting normalized regulation strength scores *b* vary from 0 to 1, where low scores denote TFBS variants with weak TF binding (weak reporter repression), and high scores indicate variants with strong TF binding (strong reporter repression). A score of *b* =1 signifies the highest regulation strength (repression) calculated among all TFBSs for a given TF.

#### 7.3 Combining data from triplicates

As a quality-filtering step, we eliminated TFBS variants that did not appear in all three replicates or that were represented by less than 30 reads in total (summing their read counts over the thirteen bins). Given that we have 13 bins and most sequences are typically found within an average of 3.4 bins, we wanted a minimum of 10 reads per bin to prevent misinterpretation of repression levels and also guide our threshold’s choice. This procedure reduced the fraction of TFBS variants for which we had sort-seq-based sequence data from 95%, 90%, and 93% of the total library size (65,536 sequences) for CRP, Fis and IHF, respectively, to 49%, 66% and 63%.

Subsequently, we calculated the regulation strengths *b* of the remaining TFBS variants (**Supplementary Materials and Methods 7.2**). Then, we determined the coefficient of variation of regulation strengths across replicates for each TFBS variant, which estimate the consistency of measured regulation strengths among replicates. Notably, most TFBS variants exhibited a coefficient of variation (CV) below 0.5, which was the threshold we set for filtering sequences. Sequences with a CV above 0.5 were excluded from the analysis. This threshold is commonly used in transcriptional studies employing fluorescent reporters to account for transcriptional noise and measurement variability ^180^. Finally, we averaged regulation strengths for each TFBS variant across all replicates and normalized the resulting averages by the maximal observed regulation strength among all TFBSs of a given TF.

#### 7.4 Frequency matrices and sequence logos

We generated frequency matrices of nucleotides that occur in TFBSs from each fluorescence bin by counting the frequency of each nucleotide at each variable position of the TFBS library. From this data, we generated heatmaps and DNA sequence logos representing the frequency matrices graphically. A sequence logo consists of a stack of the letters A, C, G, and T, at each position of a DNA sequence, where the relative size of each letter indicates its frequency in the sequence. The total height of the stack corresponds to the information content of that position, in bits ^49,196^. A sequence logo is a graphical representation of the informational properties of a TFBS. When mutated, nucleotides with high information content are more likely to lead to a loss of binding (repression) than nucleotides with low information content.

#### 7.5 Creation of genotype networks and determining network metrics

We used in-house Python script and the Python package *igraph*^197^ to generate directed genotype networks. These are graphs in which TFBS variants are nodes (vertices, genotypes), and variants that differ in a single nucleotide are connected by an edge. Each node of this network is associated with the corresponding DNA sequence and the associated regulation strength. Each edge is directed, i.e., it corresponds to a binding-score-increasing mutation, and points from a TFBS variant with lower regulation strength to a neighbor with higher regulation strength. We extracted the largest weakly connected subgraph (the“giant component” ^6^) of the network, and used it for all further analyses. This giant component comprises the vast majority (99%) of sequenced genotypes for all of our TF landscapes. We used in-house Python and R scripts for all network analyses described below.

*<u>Epistasis:</u>* Epistasis refers to non-additive interactions between two or more mutations. It can impose severe constraints on molecular evolution, because the mutations that are beneficial in one genetic background may be deleterious in another^24^. Epistasis between two mutations can be classified as magnitude, simple sign, or reciprocal sign epistasis, depending on the sign (i.e., positive or negative) of the fitness effect of individual mutations and their combinations^24^. In magnitude epistasis, the effect of a mutation on regulation strength varies depending on the genetic background but the sign of this effect (increasing or decreasing regulation strength) does not. Simple sign epistasis occurs if one single mutant has a lower regulation strength than both the wild type and the double mutant, while the other single mutant has a regulation strength that is intermediate to the wild type and double mutant. Reciprocal sign epistasis occurs when both mutations independently decrease regulation strength, but their combination increases regulation strength. The presence of reciprocal sign epistasis is a necessary condition for the existence of multiple peaks in an adaptive landscape ^93,94^.

To determine the incidence of epistasis in our landscapes, we employed a method that involves identifying all “squares” in a genotype network with the *motifs* function from the *igraph* library in R. Each square consists of a “wild-type” sequence, a double-nucleotide mutant, and the corresponding two single mutants. We assessed epistasis for each square along a single axis by selecting the highest- regulation strength sequence as the double mutant. Our analysis placed each square into one of three categories: no sign epistasis, simple sign epistasis, and reciprocal sign epistasis. The no sign epistasis category included both magnitude epistasis and additivity (no epistasis) without differentiating between them, because neither affects the accessibility of landscape peaks through only binding-increasing mutations^11,108^. We determined the proportion of all squares that fell into each category (see **Supplementary Table S5**).

*<u>Peaks:</u>* A peak is a genotype (TF binding site variant) whose neighbors all convey lower regulation strength than itself. Two peaks are connected if they are neighbors and convey the same regulation strength. We refer to the genotype with the highest regulation strength as the summit or global peak^6,198^.

*<u>Genotype connectivity:</u>* To evaluate the sparsity of our networks and the likelihood of peak misclassification due to the incompleteness of our landscapes, we used a metric called “genotype connectivity.” This represents the number of immediate (1-mutant) neighbors of a given genotype for which our experiment produced repression strength data. A perfect experiment creating a complete landscape data set would provide repression data for each of our 4^8^ genotype and all of its 24 immediate neighbors. In practice, however, data for some genotypes and some of their neighbors is unavailable. We quantified the “relative connectivity” of any one genotype as the ratio of the actual number of adjacent genotypes for which we have repression data to the theoretical maximum of 24 neighbors. This measure helps us to address questions about the sparsity of our landscape and its implications. Specifically, it allows us to determine if our sample is biased, with some regions or genotypes being more connected than others. This is particularly important for assessing whether peaks and high peaks are less connected than non-peaks, which could potentially lead to peak misassignments.

*<u>Accessible Paths:</u>* In the context of our genotype networks, we call a mutational path accessible if the regulation strength increases with each mutational step ^6,198^. We systematically enumerated all shortest accessible paths, distinguishing them by their path length, i.e., by the number of mutational steps in a path. Notably, there can be multiple alternative accessible paths with the same number of mutational steps from a single starting genotype to a peak.

*<u>Basins of attraction:</u>* The basin of attraction of a peak comprises all TFBS variants from which accessible paths to the peak exist. We refer to the basin’s size as the number of variants in the basin. We determined basin sizes by exhaustive enumeration.

<u>Overlap between basins.</u> The basins of attraction of different peaks may comprise overlapping sets of variants. To determine the overlap between two basins B_1_ and B_2_, we used the Jaccard index *J*^199,200^, which is equal to the size of the intersection between two sets of variants divided by the size of their union:

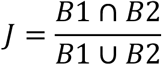

#### 7.6 Generating randomly shuffled landscapes

To generate uncorrelated random landscapes for our three TF landscapes, we performed a random shuffling of regulation strength values from our genotypes, followed by landscape analysis to identify peaks. To this end, we employed an ad-hoc python script to randomly permute experimentally measured regulation strength values among all genotypes in each of our landscape. This procedure preserves the distribution of regulation strengths in each landscape, but also creates an uncorrelated random landscape^91,97^. Following random shuffling, we analyzed each shuffled landscape’s topography with the same methods we had applied to the original (unshuffled) landscape to identify its peaks. To ensure statistical robustness, we repeated the random shuffling and peak identification 10^3^ times. We then compared the number of peaks between the 10^3^shuffled landscapes and the original (unshuffled) landscape.

#### 7.7 Principal component analysis

In order to investigate which nucleotides in a TFBS are most important for changes in regulation strengths, we performed a principal component analysis for each landscape (**Supplementary Figures S19-S21**). To this end, we first one-hot-encoded our data. This encoding represents each categorical value (nucleotide at each position of a DNA string) as a binary vector of length 4. It thus converts an entire DNA string of length L into a 4xL binary matrix. We used this binary matrix to perform PCA with the R base function *prcomp*.

### 8. Simulated adaptive walks

We simulated the adaptive evolution of a population on our landscapes by performing two different types of random walks, “greedy” adaptive random walks, and random walks based on Kimura’s model of fixation probabilities.

For all simulations, we assumed that only point mutations occur and that the time it takes for a point mutation to become fixed in a population is much shorter than the time it takes for a new mutation to appear that will eventually go to fixation. This scenario is also called the strong selection weak mutation (SSWM) scenario ^16,99–101^. Under this scenario, evolving populations are monomorphic most of the time, i.e., all individuals have the same genotype. This scenario is realistic when the product of effective population size *N* and mutation rate *μ* is small (*Nμ* < 1), which is the case for *E. coli* (N=1.8*×10*^8^*, μ=2×*10^-10^) ^67^. It allows us to model adaptive evolution as an adaptive random walk in our landscapes. Our simulations also account for mutation bias, which means that different types of mutation have a different probability of occurring in a population. We use mutation biases that were experimentally determined for *Escherichia coli*^104^.

In a greedy adaptive random walk, starting from any one genotype, only the mutational neighbor that conveys the largest fitness advantage is fixed in a population. We initiated one greedy random walk from each non-peak genotype and terminated the walk once a fitness peak was reached, i.e., once every neighbor of the current genotype had lower fitness than the genotype itself. Because every greedy random walk is deterministic in the absence of neutral mutations, it is sufficient to initiate one greedy random walk per starting genotype.

Another well-established model for random walks allows for genetic drift. It uses fixation probabilities computed by Kimura^98,201,202^ i.e., *f_ij_* = (1 – *e*^-2*s*^) / (1 – *e*^-2*Ns*^), where *f_ij_* is the probability of fixing mutation *j* in the background of genotype *i*, *N* is the effective population size, and *s* is the selection coefficient, i.e. the difference in fitness between genotypes *i* and *j*^98,201^. For a given pair of genotypes, the only parameter of this model is the effective population size *N*, for which we explored values of *N*=10^8^, *N*=10^5^ and *N*=10^2^. Generally, the smaller the population size is, the larger is the probability that neutral or deleterious mutations become fixed in a population.

For these “Kimura” adaptive walks, we chose 15,000 random starting genotypes for each of the three values of *N*, and simulated 1,000 random walks for each of them. At each step, we randomly picked a mutation *j*, generated a random number in the interval [0, 1], and considered the mutation to become fixed if the random number fell within the interval [0, *f_ij_*]. Computationally, we accelerated this process by precomputing all fixation probabilities for all genetic backgrounds, and then used the *python* function *numpy.choice* to generate a random sample from the multinomial distribution of fixation probabilities at each step^203^. Because of genetic drift, Kimura’s random walks do not necessarily terminate when they reach a fitness peak. For this reason, we simulated each such random walk for a maximum of 25 mutational steps.

### 9. Randomized landscape null model for peak accessibility

To evaluate whether the accessibility of strong regulatory peaks observed in the empirical landscapes exceeds random expectations, we constructed randomized “null” landscapes and repeated evolutionary simulations on them under identical conditionsas for the empirical landscapes.

Specifically, for each transcription factor landscape, we generated 10^$^independently shuffled landscapes landscapes see **Supplemtnary Methods 7.6**) by permuting repression-strength values across genotypes while preserving the network of genotypes and its mutational neighborhood structure. This procedure maintains the number of genotypes, their connectivity, and the distribution of repression-strength values, while removing correlations between genotype and phenotype. All other aspects of the analysis were kept identical to those used for the empirical landscapes.

For each randomized landscape, we simulated adaptive evolution using the same Kimura random-walk framework as for the empirical data. We assumed a population size of 10^8^ individuals, initiated adaptive walks from the same number of starting genotypes, and used identical stopping criteria, fixation probabilities, and mutational bias parameters as described in **Supplementary Methods 8** (*“Simulated adaptive walks”)*.

For each randomized landscape, we computed the fraction of adaptive walks that reached a high regulatory peak. This yielded a null distribution of peak-accessibility values for each transcription factor. We tested the null hypothesis that the observed fraction from the corresponding empirical landscape is greater than the mean fraction derived from this null distribution using a one-sided Monte Carlo permutation test (𝑁 = 10^$^ randomized landscapes). For this test, we used Z-scores computed as the difference between the observed accessibility and the mean of the shuffled-landscape distribution, normalized by the standard deviation of that distribution.

### 10. Incorporating experimental uncertainty into adaptive walks

Fluorescence-based sort-seq measurements are subject to experimental uncertainty arising from finite fluorescence bin resolution, variability in fluorescence measurements, and sequencing noise. To account for this uncertainty when defining fitness relationships between genotypes and identifying peaks in the landscape, we incorporated empirically estimated noise into pairwise repression strength comparisons.

For each genotype, we computed repression strength (*S*) as a continuous quantity derived from the distribution of sequencing reads across fluorescence bins (see Methods). We quantified experimental uncertainty in *S* by a genotype-specific noise parameter (*τ*_*G*_), calculated as the standard deviation of repression strength across three biological replicates. This parameter defines an uncertainty interval within which repression strength values cannot be reliably distinguished.

We compared the repression strengths of pairs of genotypes while explicitly accounting for their associated experimental uncertainty. Let *S*_*A*_ and *S_B_* denote the repression strengths of genotypes *A* and *B*, with corresponding uncertainty estimates 𝜏_*A*_ and 𝜏_*B*_. We considered genotype *A* to exhibit significantly stronger repression than genotype *B* if

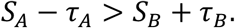

In this case, we assigned a directed edge from *A* to *B* in the genotype network. Conversely, genotype *A* was considered to exhibit significantly weaker repression than genotype *B* if

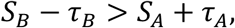

in which case we assigned a directed edge from *B* to *A*. If neither condition was satisfied—i.e., if the uncertainty intervals of the two genotypes overlapped—we considered their repression strengths to be indistinguishable within experimental uncertainty. Such genotype pairs were treated as neutral and connected by two directed edges, one in each direction.

Applying these rules to all genotype pairs yielded the uncertainty-aware genotype network, denoted 𝒢^*S,τ*^. For comparison, we also constructed a noise-free genotype network, denoted 𝒢^*S*^, in which experimental uncertainty was ignored (*τ*_*G*_ = 0) and pairwise comparisons were based solely on point estimates of repression strength *S*. In both networks, peaks were defined as genotypes without outgoing edges.

## SUPPLEMENTARY TABLES

**Table S1.**
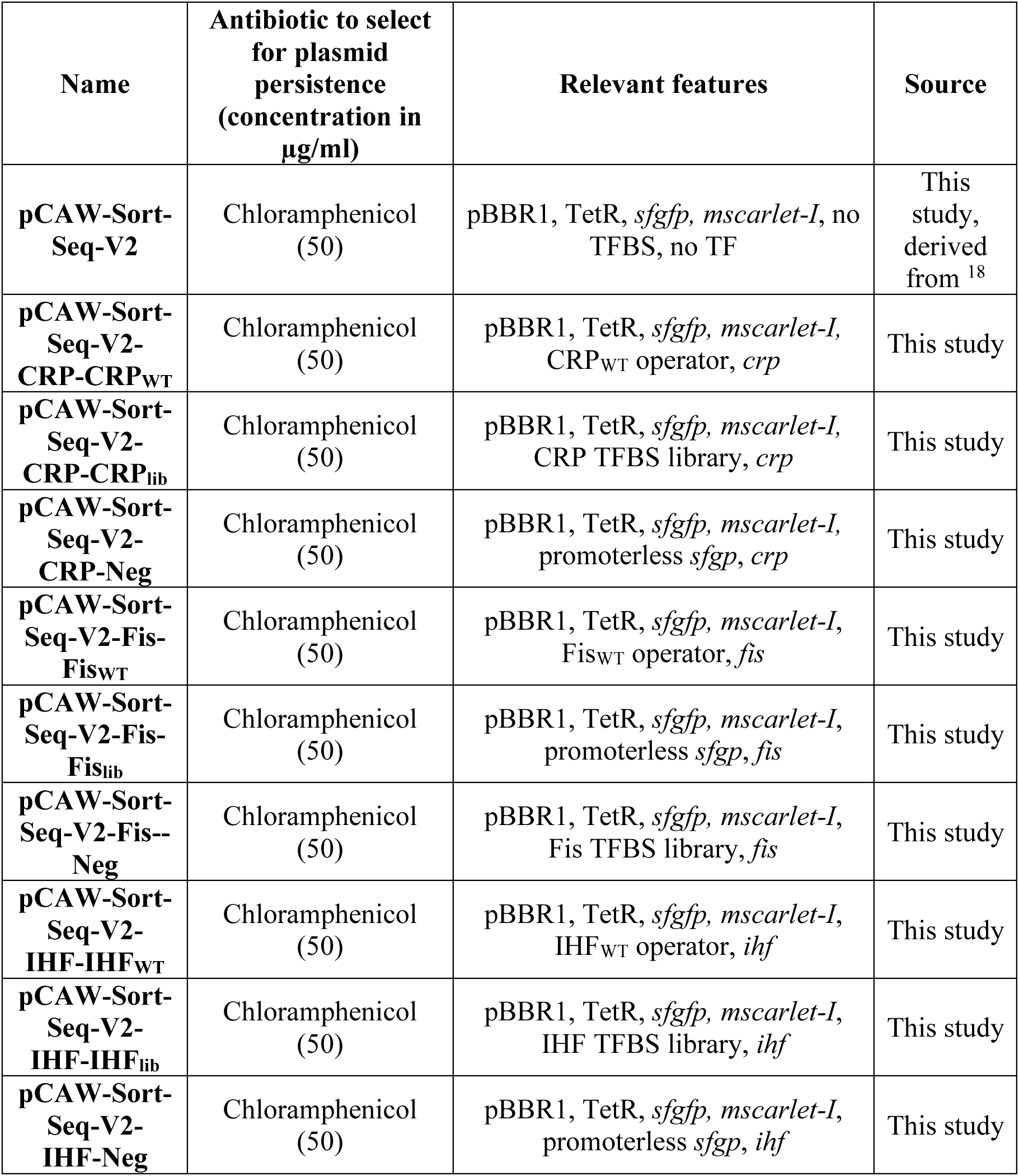
Plasmids used in this study.

**Table S2.** Table of strains.

| Strain | Genotype | Antibiotic resistance | Reference |
| --- | --- | --- | --- |
| SIG10-MAX<br>from Sigma<br>Aldrich<br>*Cloning strain* | F- mcrA $\Delta$(mrr-hsdRMS-mcrBC)<br>endA1 recA1 Φ80dlacZΔM15<br>ΔlacX74 araD139<br>$\Delta$(ara,leu)7697galU galK rpsL<br>nupG λ- tonA (StrR) | Streptomycin | Sigma Aldrich |
| *E. coli* DH5α<br>*Cloning strain* | F- endA1 gln V44 thi-1 recA1<br>relA1 gyrA96 deoR nupG<br>Φ80dlacZΔM15 $\Delta$(lacZYA-<br>argF)U169, hsdR17(rK – mK + ),<br>λ– | None | 204 |
| *E. coli* JW5702-4<br>*Δcrp mutant strain.* | $\Delta$(araD-araB)567,<br>ΔlacZ4787(::rrnB-3), λ–, Δcrp-765::kan, rph-1, $\Delta$(rhaD-rhaB)568, hsdR514. | Kanamycin | 179 |
| *E. coli* JW1702-1<br>*ΔihfA mutant strain.* | $\Delta$(araD-araB)567,<br>ΔlacZ4787(::rrnB-3), λ–,<br>ΔihfA786::kan, rph-1, $\Delta$(rhaD-rhaB)568, hsdR514. | Kanamycin | 179 |
| *E. coli* JW3229-1<br>*Δfis mutant strain* | bΔ(araD-araB)567,<br>ΔlacZ4787(::rrnB-3), λ–, Δfis-779::kan, rph-1, $\Delta$(rhaD-rhaB)568, hsdR514. | Kanamycin | 179 |

**Supplementary Table S3.**
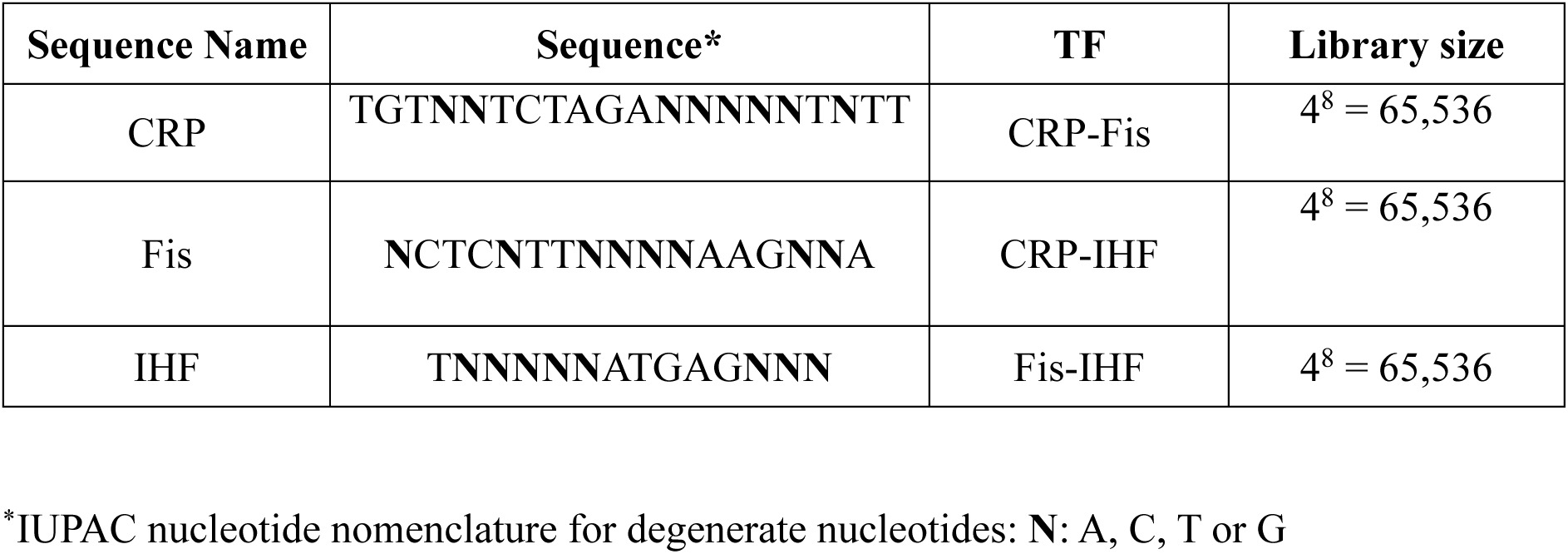
Libraries used in the present study.

**Supplementary Table S4.**
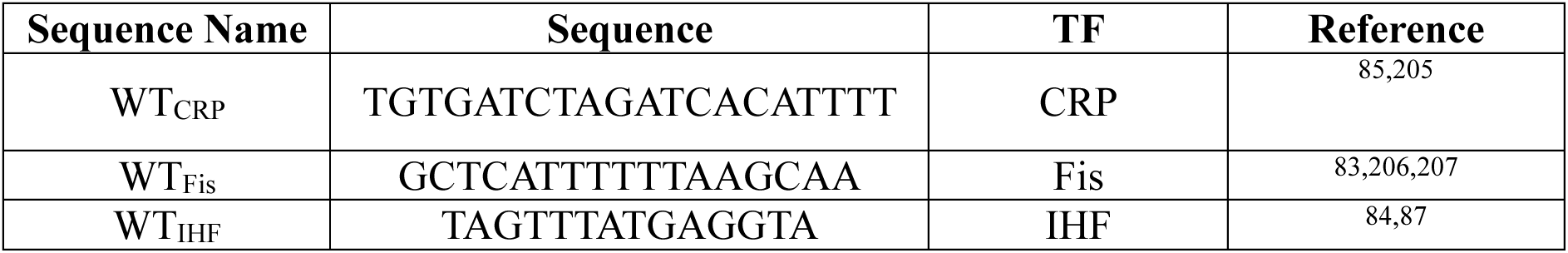
TFBS reference (“wild-type”, WT) sequences used in the present study.

**Table S5.** Global landscape properties.

|  | *Number of*<br>*genotypes<sup>1</sup>* | *Number*<br>*of peaks<sup>2</sup>* | *Number*<br>*of low*<br>*peak<sup>3</sup>* | *Number*<br>*of high*<br>*peaks<sup>4</sup>* | *Number of*<br>*squares<sup>5</sup>* | *Magnitude*<br>*epistasis*<br>*or*<br>*additivity<sup>6,7</sup>* | *Simple*<br>*sign*<br>*epistasis<sup>7</sup>* | *Reciprocal*<br>*sign*<br>*epistasis<sup>7</sup>* |
| --- | --- | --- | --- | --- | --- | --- | --- | --- |
| CRP | 31,975<br>(49%)<sup>1</sup> | 2,154<br>(7%) | 2,093<br>(97.2%) | 61<br>(2.8%) | 580,289 | 37% | 34% | 28% |
| Fis | 43,222<br>(66%)<sup>1</sup> | 2,312<br>(5%) | 2,140<br>(92.6%) | 172<br>(7.4%) | 1,194,144 | 34% | 34% | 32% |
| IHF | 41,325<br>(63%)<sup>1</sup> | 2,453<br>(6%) | 2,254<br>(91.9%) | 199<br>(8.1%) | 1,026,818 | 34% | 33% | 32% |

<sup>1</sup> Percentages indicate the proportion of the studied library relative to its expected size.
<sup>2</sup> Percentages indicate the proportion of peaks relative to the total number of genotypes
<sup>3</sup> Low peaks are peaks with regulation strengths below the wild sequence. Percentages refer to the proportion of all peak genotypes.
<sup>4</sup> High peaks are peaks with regulation strengths above the wild-type sequence. Percentages refer to the proportion of all peak genotypes.
<sup>5</sup> A square represents a quadruplet of sequences containing a focal sequence (ab), one double mutant of this sequence (AB), and the two intermediate single mutants (Ab and aB).
<sup>6</sup> This category includes both magnitude epistasis and additivity (no epistasis) without distinguishing them, because neither of the two subcategories affects peak accessibility<sup>11,108</sup>
<sup>7</sup> Percentages refer to the proportion of all squares.

**Table S6.**
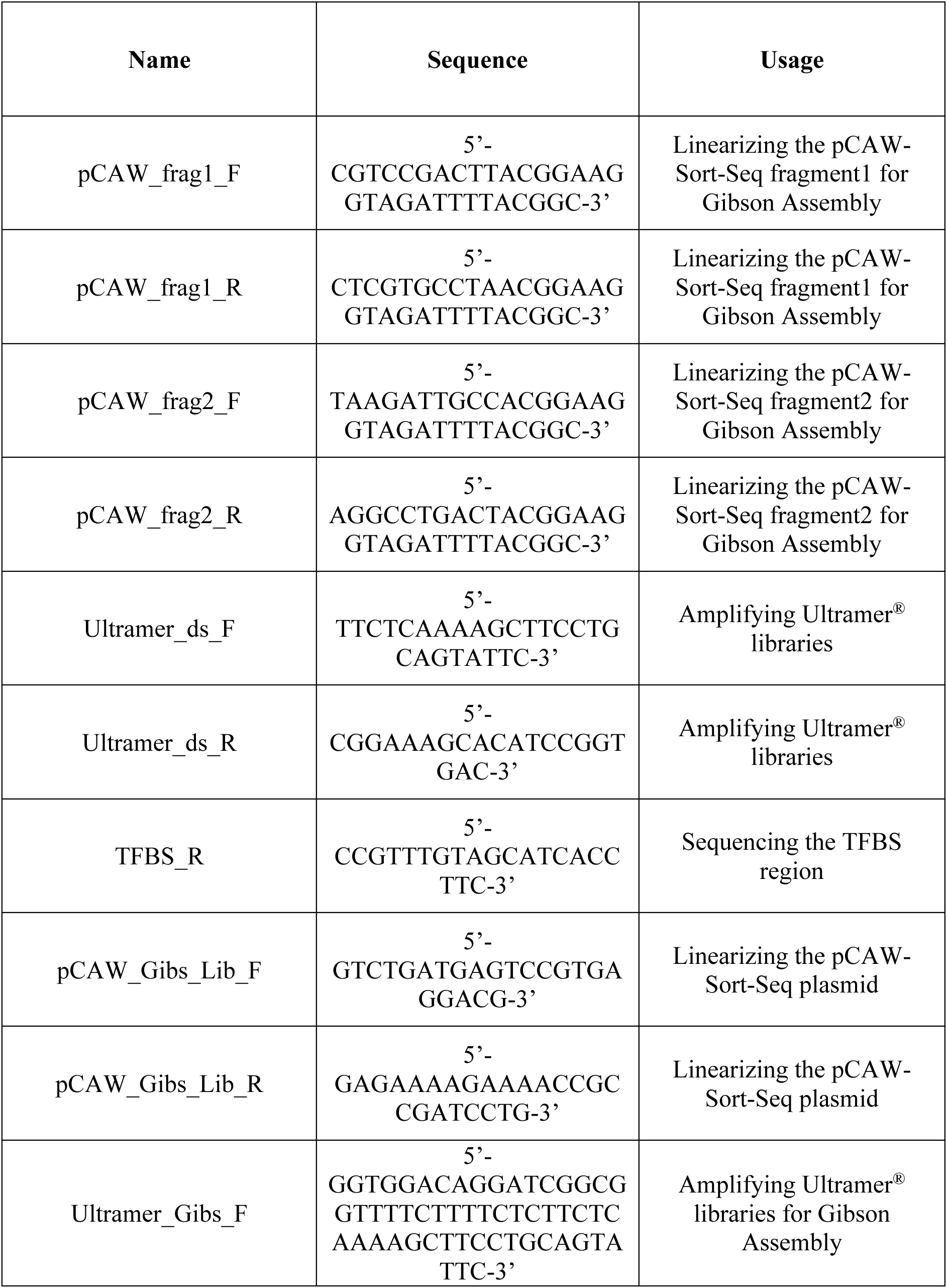
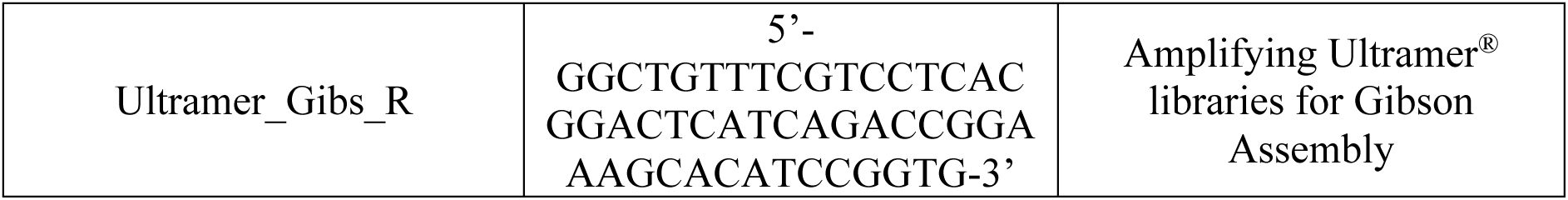
Primers for engineering the pCAW-Sort-Seq-V2 plasmid and for cloning libraries.

## SUPPLEMENTARY FIGURES

**Supplementary Figure S1.**
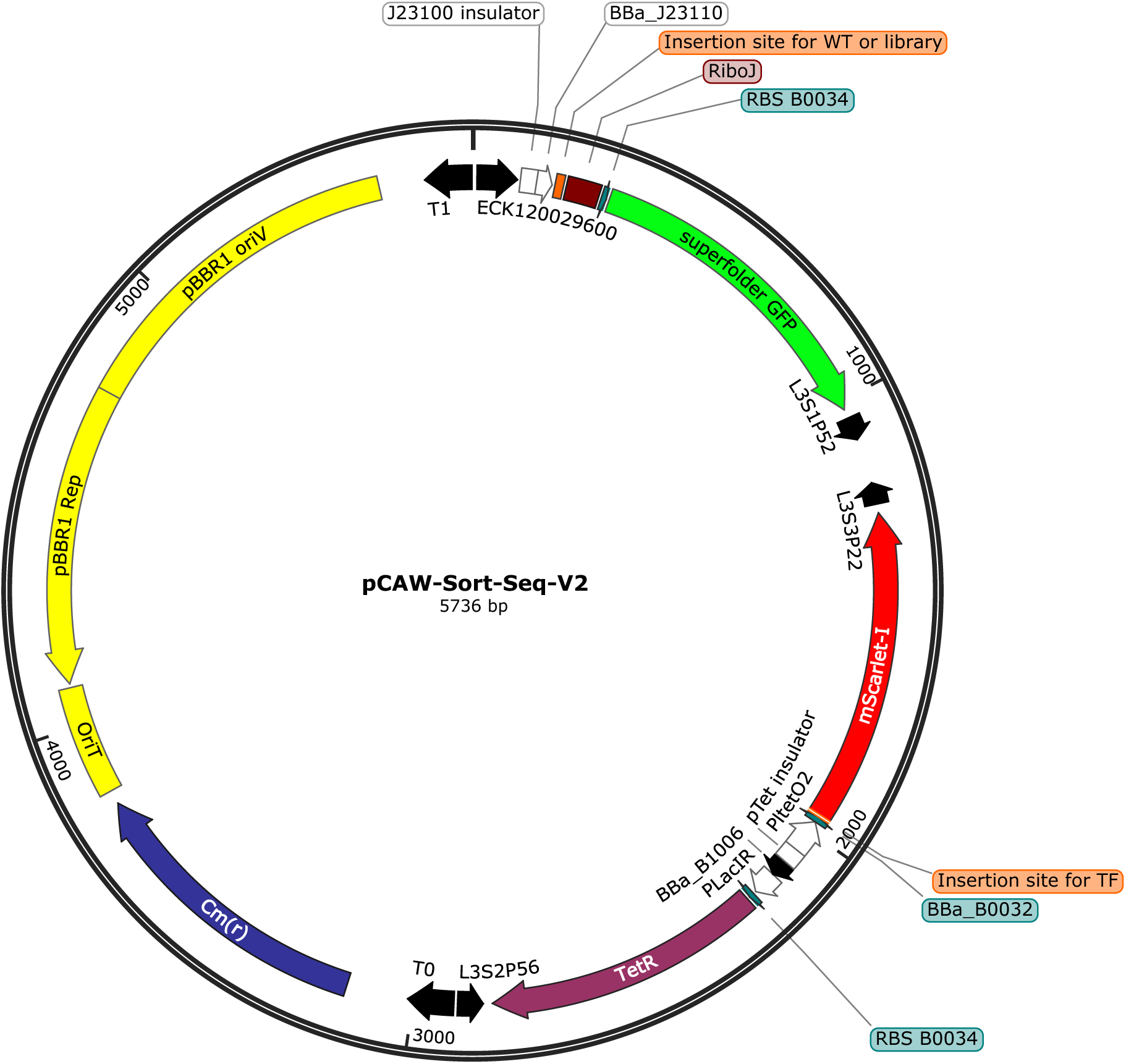
Components of plasmid pCAW-Sort-Seq-V2. The plasmid pCAW-Sort-Seq-V2 is a derivative of the plasmid pCAW-Sort-Seq^18^. It encodes the broad-host-range, low-copy-number replication origin pBBR1 ^185^ (in yellow, approximately 5–10 copies per cell^185^), together with an origin of transfer (OriT) for conjugation (yellow), and a chloramphenicol resistance gene (dark blue). The interchangeable regulatory region where the TF binding site is inserted (orange) is positioned between a constitutive promoter (BBa_J23110 ^208^) and a superfolder GFP (*sfgfp*, green) fluorescent reporter gene ^187^. The transcriptional insulator RiboJ^188^ (dark red) is located upstream of the *sfgfp* gene. The *tetR* gene (purple), derived from the original Tn10 transposon^186^, is expressed under the control of a low-strength constitutive promoter (a pLac promoter variant ^209^). TetR regulates the expression of a bicistronic operon encoding the TF of interest and the red fluorescent reporter *mScarlet-I* ^189^. The TF coding region is inserted at the TF insertion site (orange). Promoters and their associated insulators^210^ are represented as white arrows, transcriptional terminators as black arrows ^211^, and ribosome binding sites (RBSs) as small teal rectangles.

**Supplementary Figure S2.**
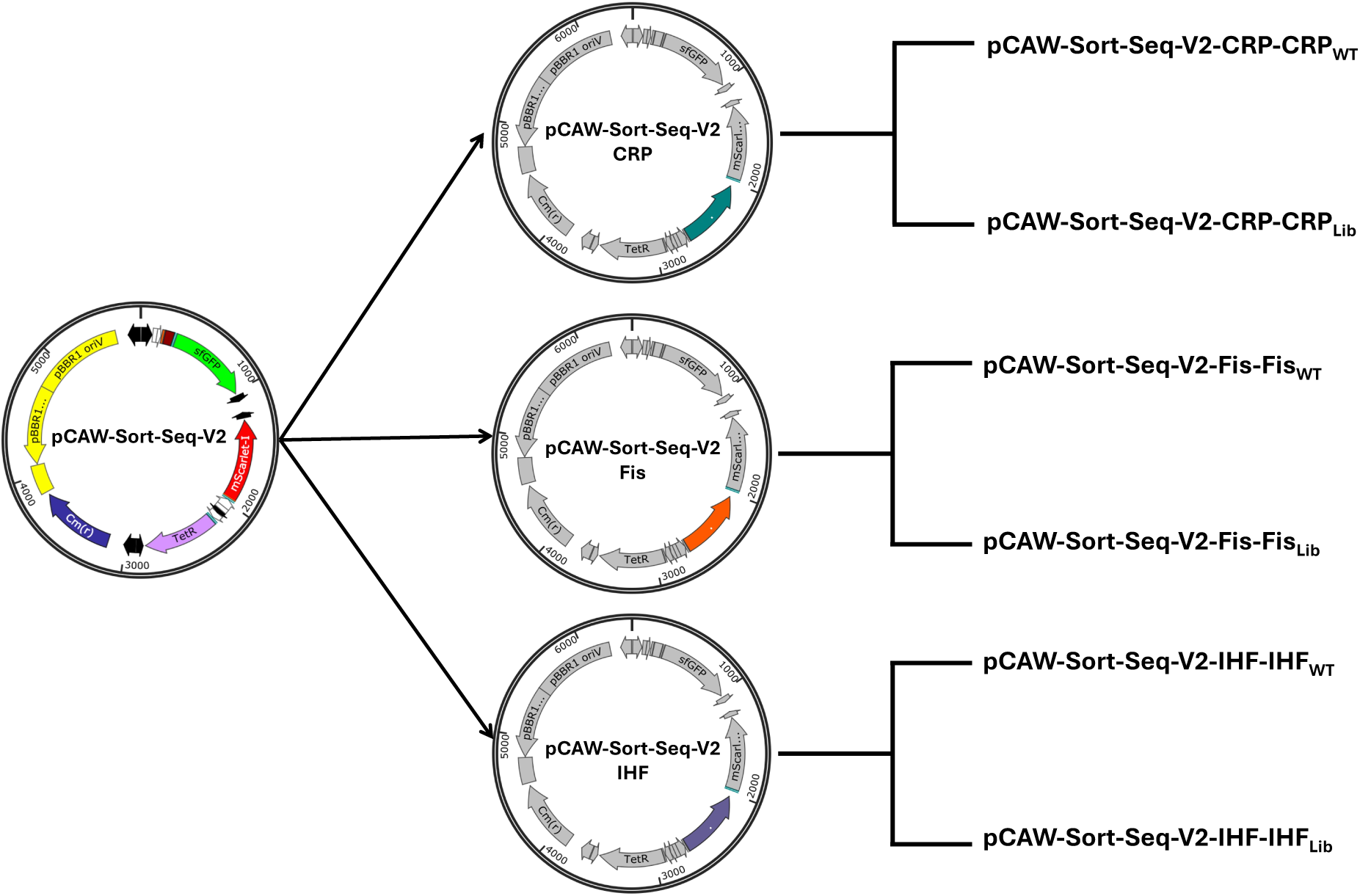
Construction of pCAW-Sort-Seq-V2 global TF derivatives. The plasmid pCAW-Sort-Seq-V2^18^ was derived from pCAW-Sort-Seq through a series of sequential cloning steps (see Supplementary Methods). First, a codon-optimized *mscarlet* expression cassette was synthesized and cloned into the parental pCAW-Sort-Seq backbone in an orientation antiparallel to the tetracycline resistance gene (*tetR*), generating the intermediate plasmid pCAW-Sort-Seq-*mscarlet*. Next, individual global transcription factor genes were inserted upstream of *mscarlet* to form a bicistronic operon, yielding three TF-specific intermediate plasmids expressing CRP, Fis, or IHF (CRP shown in green, Fis in orange, and IHF in purple; all other plasmid features are shown in grey for clarity). In the final step, depicted here, the appropriate wild-type TF binding sites and corresponding TFBS libraries were cloned upstream of the *gfp* reporter gene in each TF-specific backbone, producing the final library plasmids used in the sort-seq experiments. An additional negative control without a promoter region for the *sfgfp* gene was created for each TF (not shown). For full cloning details, see Supplementary Methods 1-4.

**Supplementary Figure S3.**
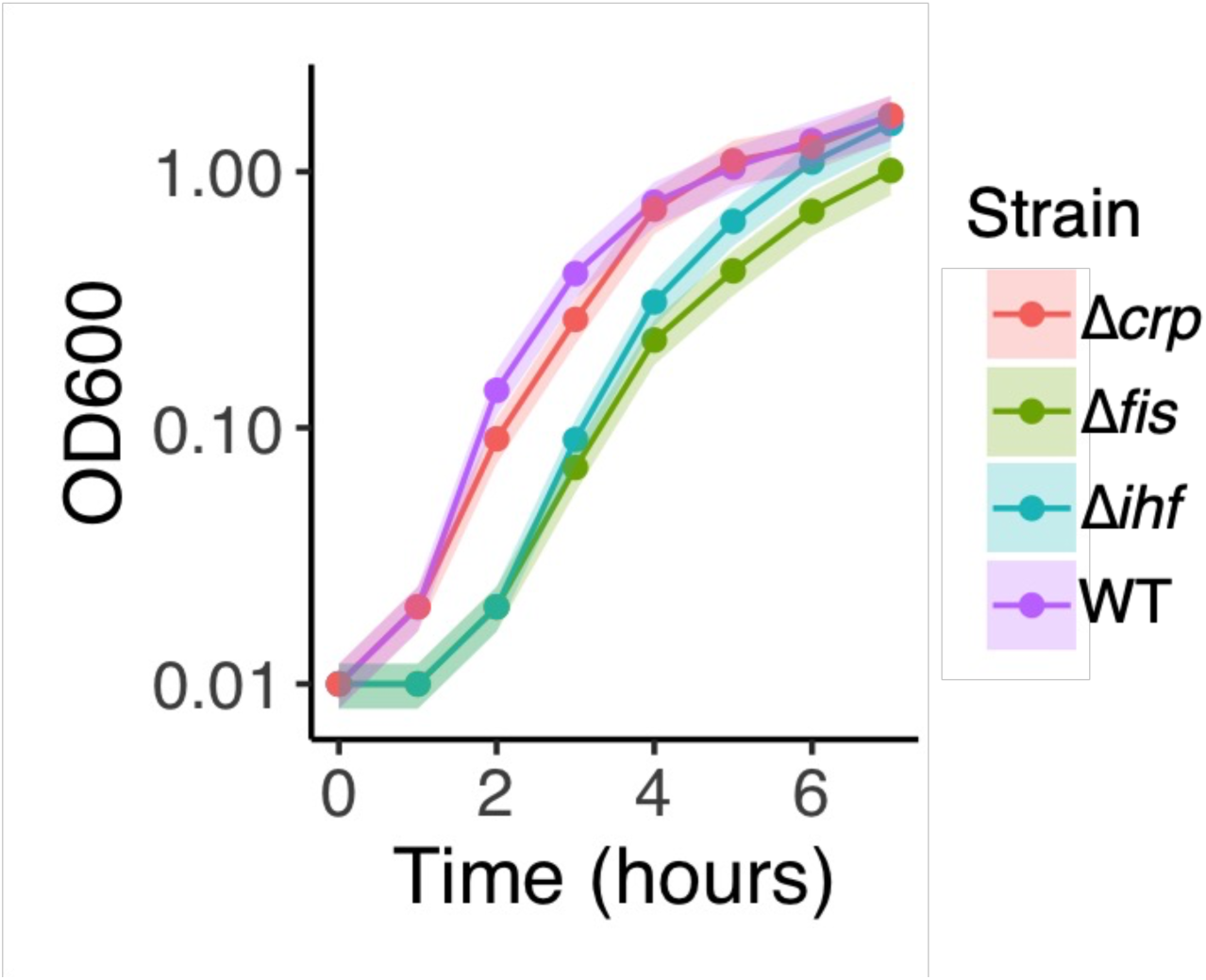
Growth profile for strains used in this study. Growth profiles for all strains used for landscape mapping are represented as optical densities (OD_600_, y-axis) measured over 7 hours of growth at 1-hour intervals (x-axis). We transformed all strains with the plasmid pCAW-Sort-Seq-V2 and grew them in LB medium without inducers. Colors (legend) represent the different strains. Wild-type (purple) refers to the BW25113 parent-strain from which the mutants (Δ*crp*, *Δfis*, Δ*ihf*) were derived. Each circle represents the mean of three biological replicates independently measured in a plate reader. Shaded areas correspond to one standard deviation calculated from the three replicates.

**Supplementary Figure S4.**
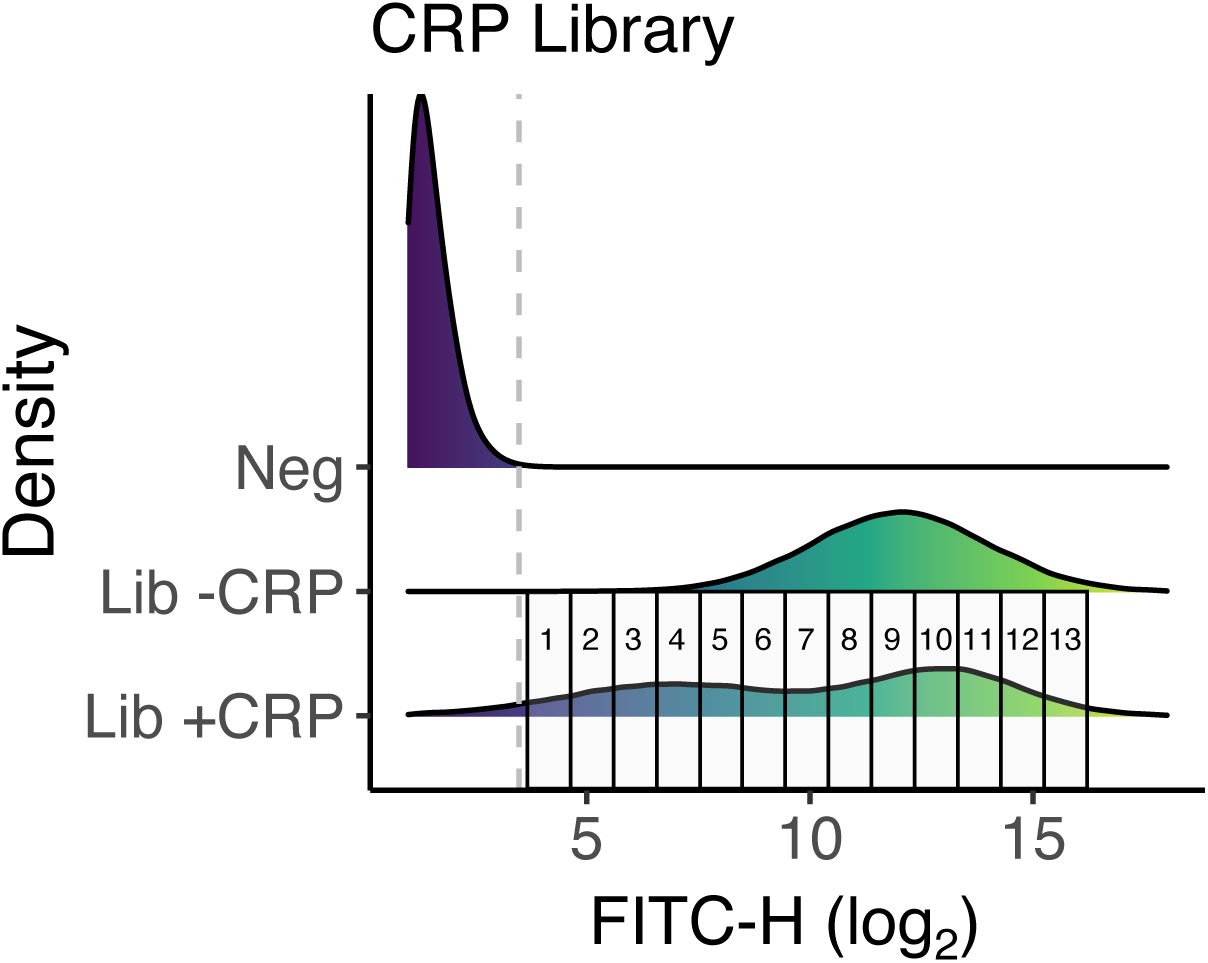
Sorting procedure for CRP library. Each density plot is based on a histogram representing the distribution of GFP fluorescence for 10^6^ cells. The histogram represents data from one replicate, and the same sorting strategy was applied to the other two replicates. Specifically, the horizontal axis shows GFP fluorescence (FITC-H values, arbitrary units, log_2_ scale), while the vertical axis indicates the relative frequency of cells with a given fluorescence value. Heatmap colors indicate fluorescence, from low (dark purple) to high (yellow). The vertical axis indicates the three experimental conditions under which we measured fluorescence: 1) negative control (“Neg”), without GFP expression (promoterless pCAW-v2); 2) CRP library in pCAW-v2 without TF expression (“Lib -CRP”) and 3) CRP library in pCAW-v2 with TF expression (“Lib +CRP”). To create each density plot from the histogram data, we performed density smoothing using a Gaussian kernel function with the aid of the *ggplot2* package^212^. The vertical dashed grey line marks the upper limit of cell autofluorescence (negative control). We did not select cells with fluorescence below this threshold for sorting. Rectangular boxes (1 to 13) represent fluorescence bins/gates into which we sorted *E.coli* cells, spaced equally on a binary logarithmic (log_2_) scale.

**Supplementary Figure S5.**
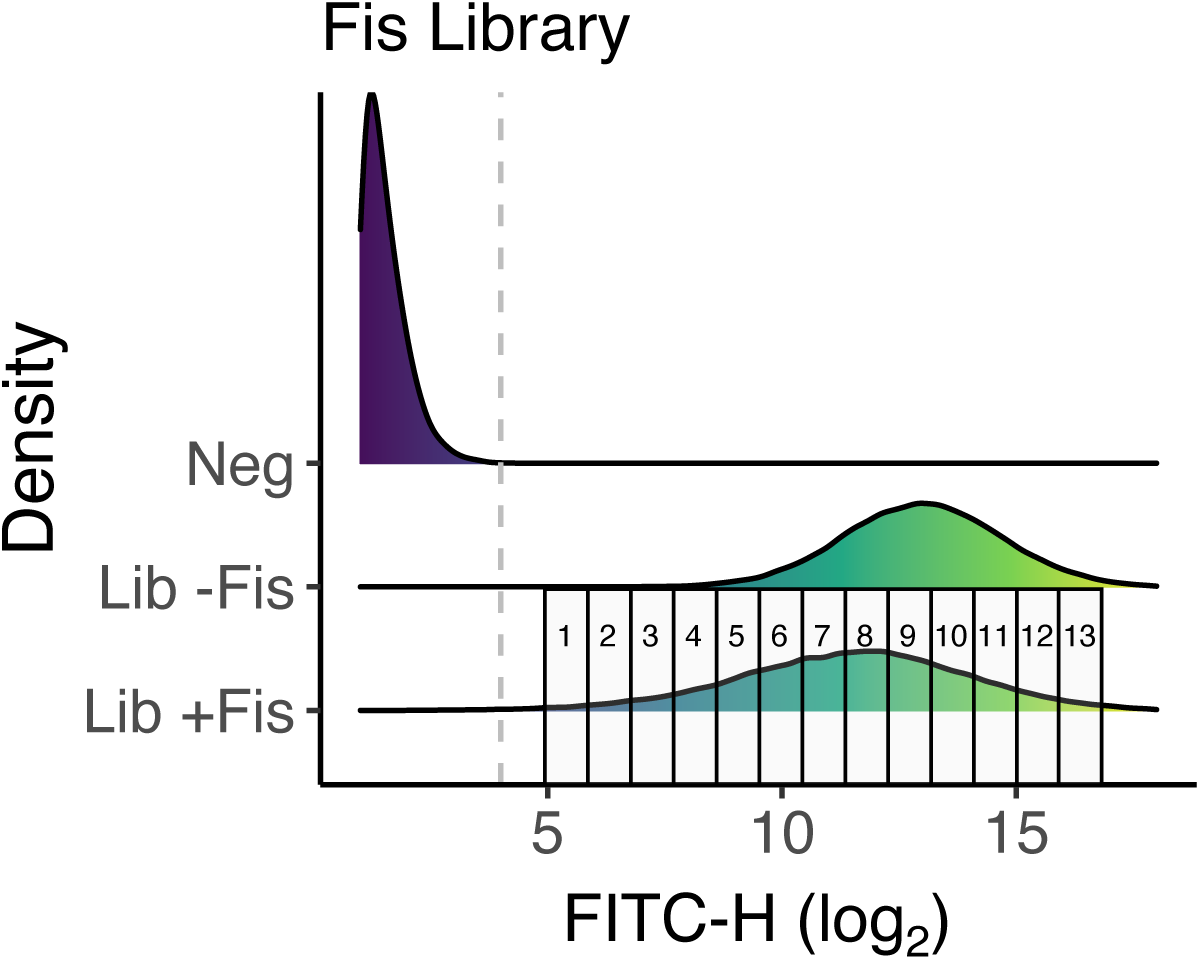
Sorting procedure for Fis library. Each density plot represents the distribution of GFP fluorescence for 10^6^ cells. The histogram represents data from one replicate, and the same sorting strategy was applied to the other two replicates. Specifically, the horizontal axis shows GFP fluorescence (FITC-H values, arbitrary units, log_2_ scale), while each histogram indicates the relative frequency of cells with a given fluorescence value. Heatmap colors indicate fluorescence, from low (dark purple) to high (yellow). The vertical axis indicates the three experimental conditions under which we measured fluorescence: 1) negative control (“Neg”), without GFP expression (promoterless pCAW-v2); 2) Fis library in pCAW-v2 without TF expression (“Lib -Fis”) and 3) Fis library in pCAW-v2 with TF expression (“Lib +Fis”). We performed density smoothing using a Gaussian kernel function to create a smooth density plot for each histogram with the aid of the *ggplot2* package ^212^. The vertical dashed grey line marks the upper limit of cell autofluorescence (negative control). We did not select cells with fluorescence below this threshold for sorting. Rectangular boxes (1 to 13) represent bins/gates for each sample, spaced equally on a binary logarithmic (log_2_) scale.

**Supplementary Figure S6.**
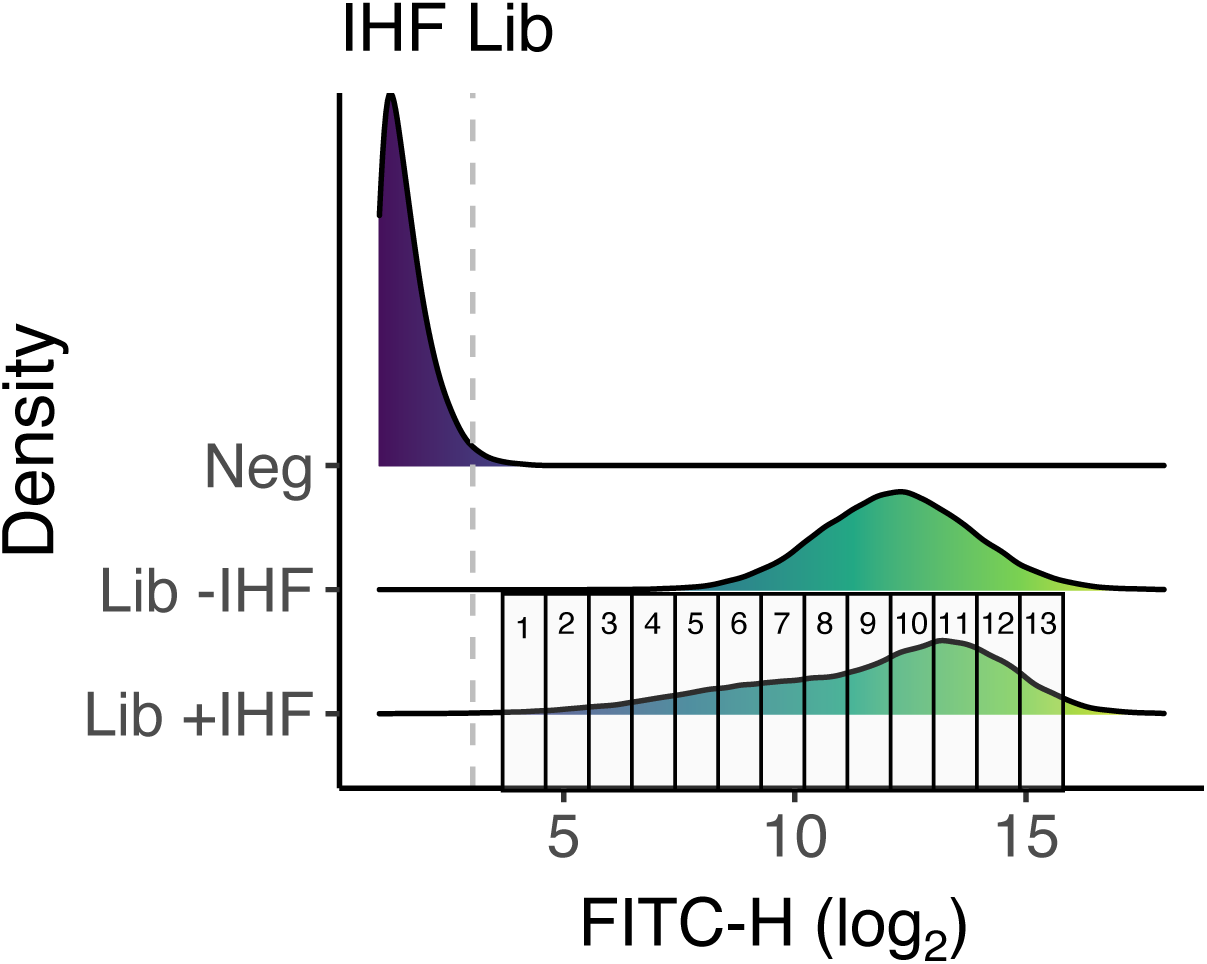
Sorting procedure for IHF library. Each density plot represents the distribution of GFP fluorescence for 10^6^ cells. The histogram represents data from one replicate, and the same sorting strategy was applied to the other two replicates. Specifically, the horizontal axis shows GFP fluorescence (FITC-H values, arbitrary units, log_2_ scale), while each histogram indicates the relative frequency of cells with a given fluorescence value. Heatmap colors indicate fluorescence, from low (dark purple) to high (yellow). The vertical axis indicates the three experimental conditions under which we measured fluorescence: 1) negative control (“Neg”), without GFP expression (promoterless pCAW-v2); 2) IHF library in pCAW-v2 without TF expression (“Lib -IHF”) and 3) IHF library in pCAW-v2 with TF expression (“Lib +IHF”). We performed density smoothing using a Gaussian kernel function to create a smooth density plot for each histogram with the aid of the *ggplot2* package^212^. The vertical dashed grey line marks the upper limit of cell autofluorescence (negative control). We did not select cells with fluorescence below this threshold for sorting. Rectangular boxes (1 to 13) represent bins/gates for each sample, spaced equally on a binary logarithmic (log_2_) scale.

**Supplementary Figure S7.**
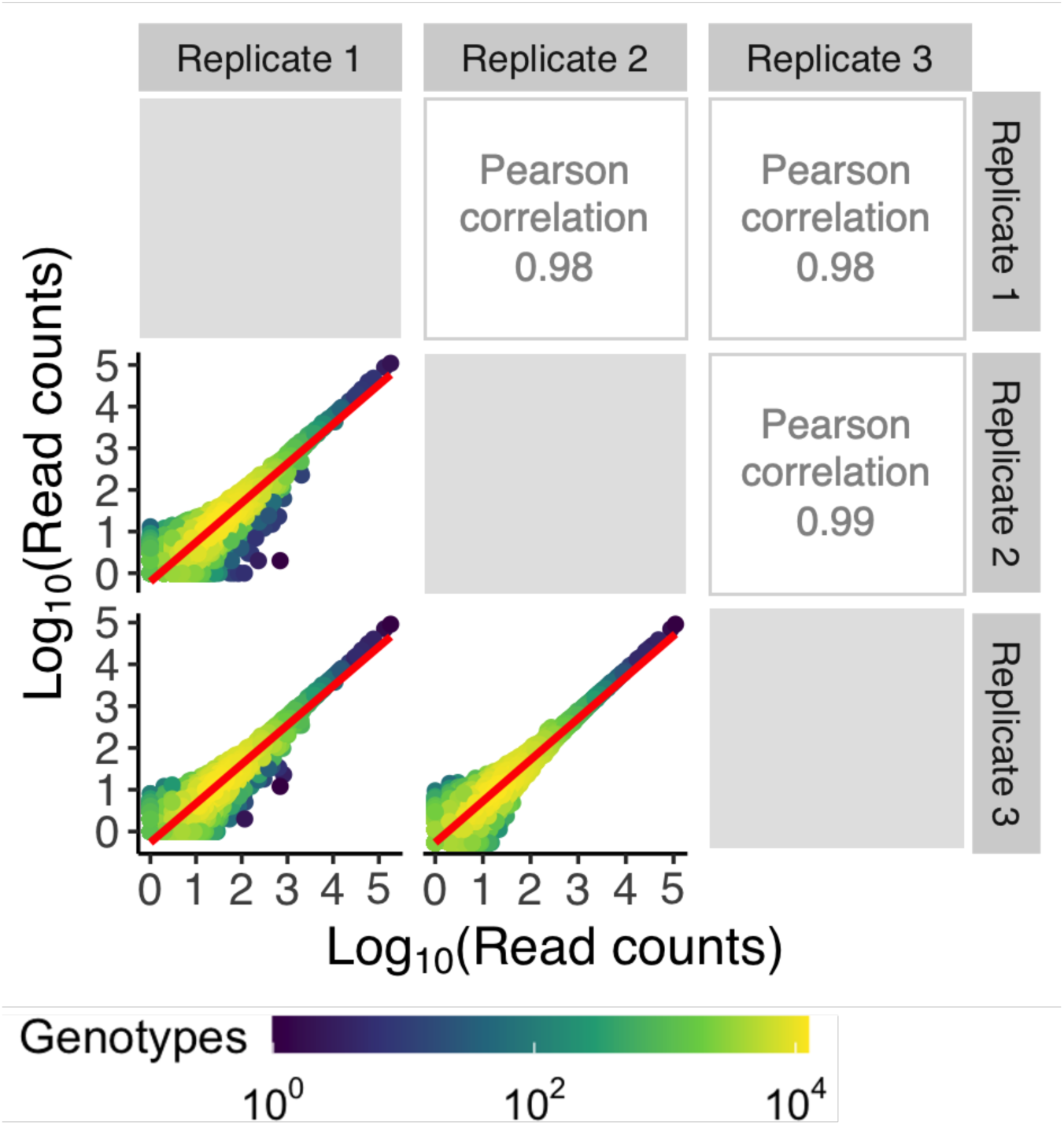
Correlation of read counts among replicates of the CRP library. The figure presents a pairwise comparison of log-transformed read counts across three experimental replicates for CRP. Each circle represents a distinct genotype, plotted according to its read counts in two sequencing replicates (x-axis for one replicate and y-axis for the other). The density of genotypes within the plot is indicated in the legend, with purple representing lower-density regions and yellow representing higher-density regions. The red line across each plot depicts a linear model’s fit to the data, illustrating the association between the read counts of genotypes across replicates. Pearson correlation coefficients (upper triangle) are indicated for each pair of replicates. Their values are close to 1, demonstrating high reproducibility.

**Supplementary Figure S8.**
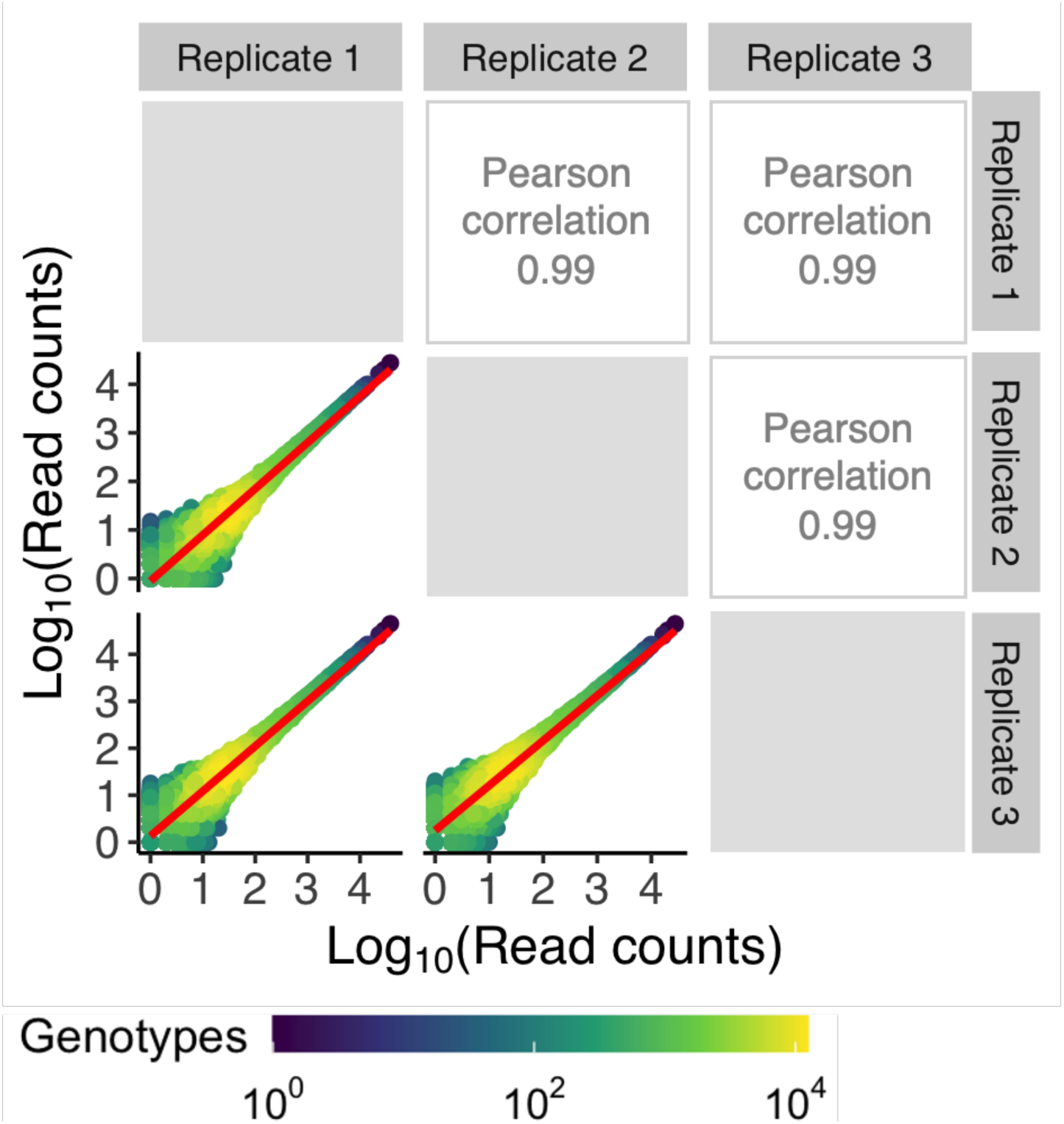
Correlation of read counts among replicates of the Fis library. The figure presents a pairwise comparison of log-transformed read counts across three experimental replicates for Fis. Each circle represents a distinct genotype, plotted according to its read counts in two sequencing replicates (x-axis for one replicate and y-axis for the other). The density of genotypes within the plot is indicated in the legend, with purple representing lower-density regions and yellow representing higher-density regions. The red line across each plot depicts a linear model’s fit to the data, illustrating the association between the read counts of genotypes across replicates. Pearson correlation coefficients (upper triangle) are indicated for each pair of replicates. Their values are close to 1, demonstrating high reproducibility.

**Supplementary Figure S9.**
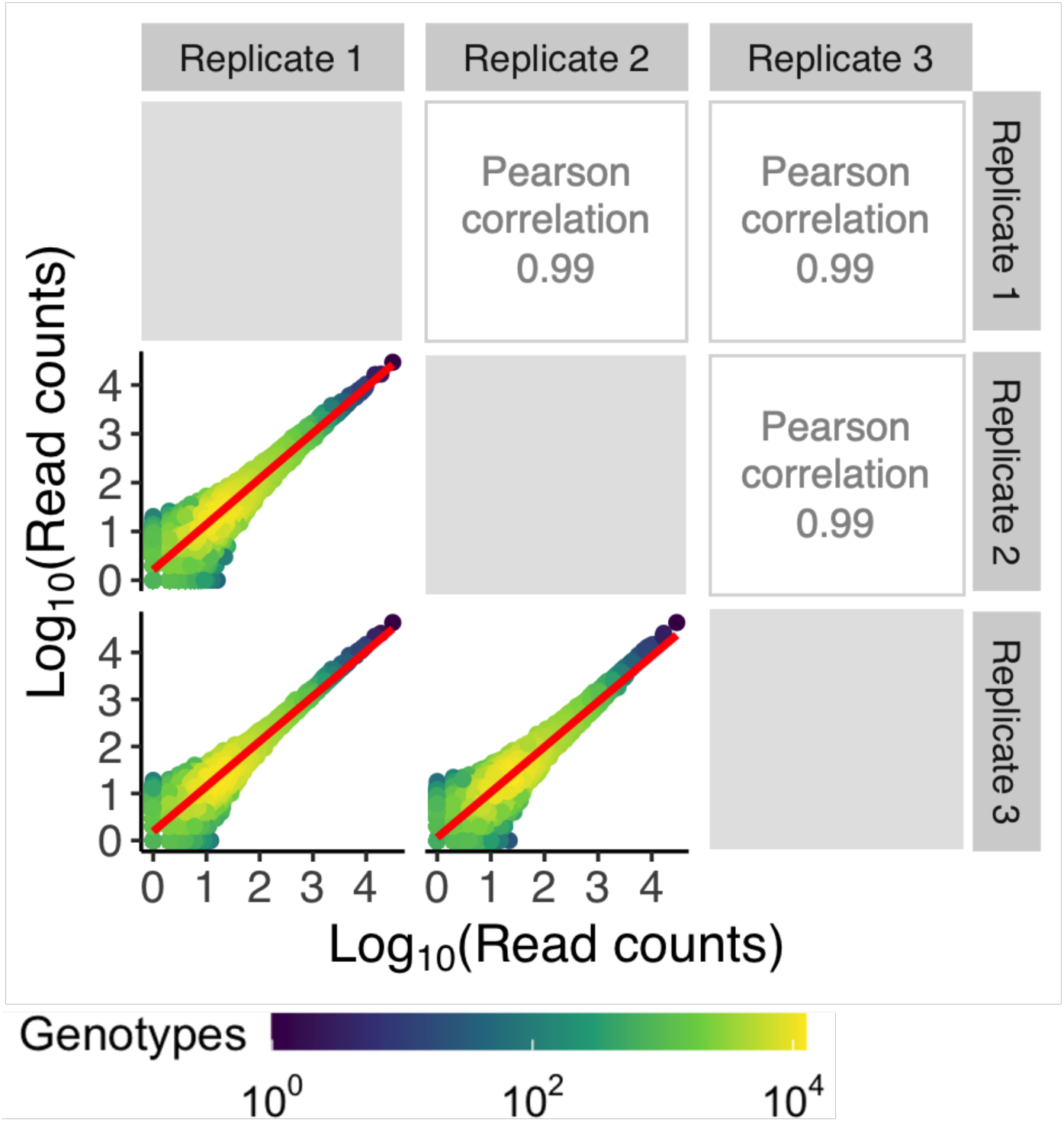
Correlation of read counts among replicates of the IHF library. The figure presents a pairwise comparison of log-transformed read counts across three experimental replicates for IHF. Each circle represents a distinct genotype, plotted according to its read counts in two sequencing replicates (x-axis for one replicate and y-axis for the other). The density of genotypes within the plot is indicated in the legend, with purple representing lower-density regions and yellow representing higher-density regions. The red line across each plot depicts a linear model’s fit to the data, illustrating the association between the read counts of genotypes across replicates. Pearson correlation coefficients (upper triangle) are indicated for each pair of replicates. Their values are close to 1, demonstrating high reproducibility.

**Supplementary Figure S10.**
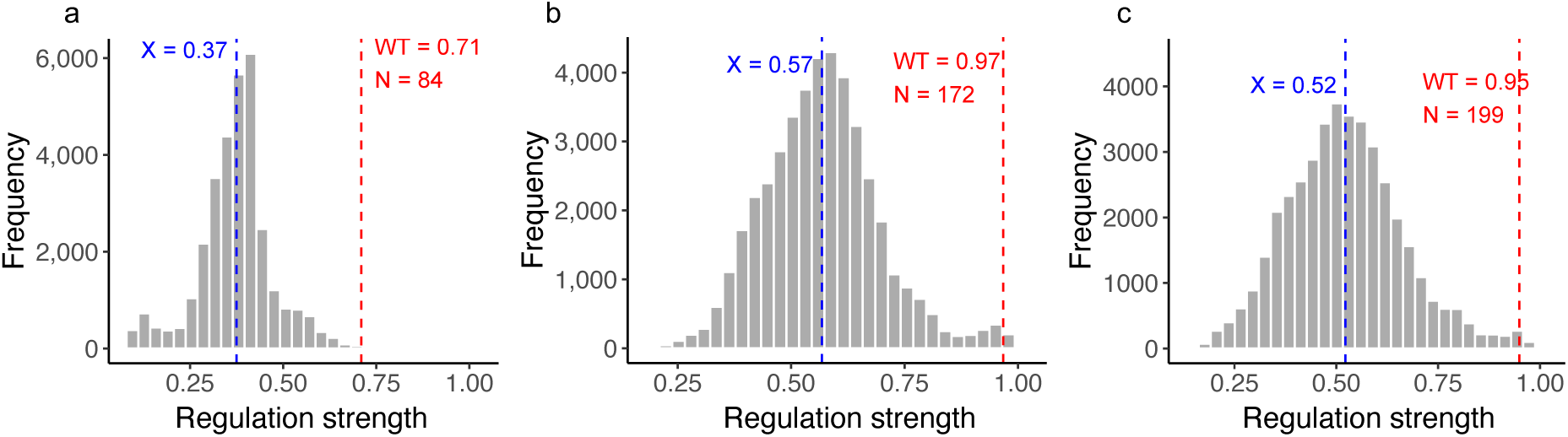
Regulation strength distribution across the libraries a. CRP library. The histogram illustrates the distribution of regulation strengths (normalized to the maximally observed value) among 31,975 library variants. The average regulation strength across all variants is highlighted by a dashed blue line (mean = 0.37), while the wild type’s (WT = 0.71) strength is denoted by a dashed red line. A total of N=84 variants surpass the WT in regulation strength. **B. Fis library.** The histogram illustrates the distribution of regulation strengths (normalized to the maximum observed value) among 43,222 library variants. The average regulation strength across all variants is highlighted by a dashed blue line (mean = 0.57), while the wild type’s (WT = 0.97) strength is denoted by a dashed red line. A total of N=172 variants surpass the WT in regulation strength. **c. IHF library.** The histogram illustrates the distribution of regulation strengths (normalized to the maximum observed value) among 41,325 library variants. The average regulation strength across all variants is highlighted by a dashed blue line (mean = 0.52), while the wild type’s (WT = 0.95) strength is denoted by a dashed red line. A total of N=199 variants surpass the WT in regulation strength.

**Supplementary Figure S11.**
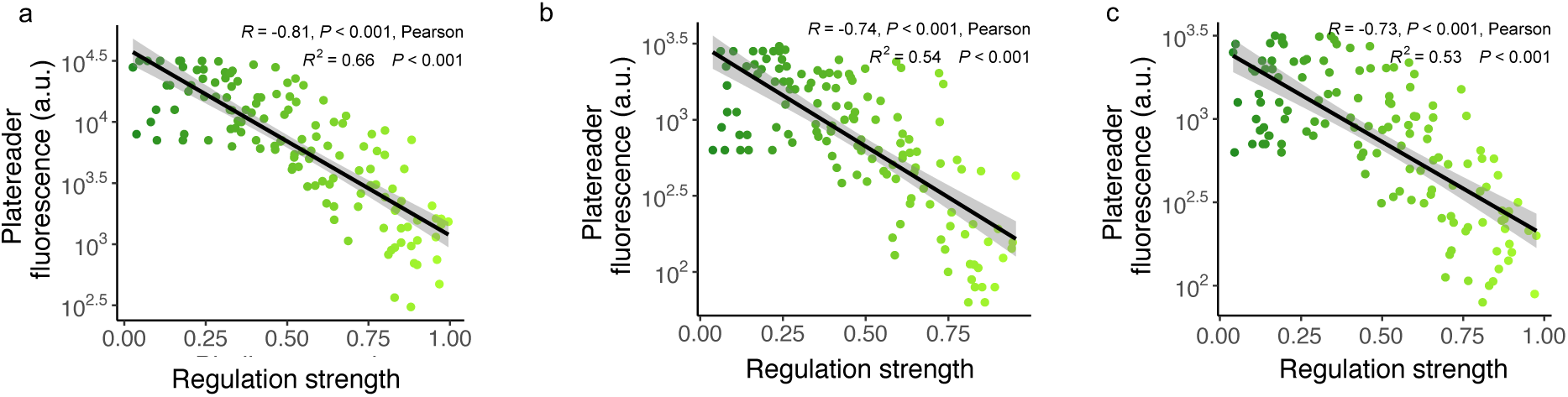
Validating regulation strengths. In this analysis, we validated the regulation strengths of 10 selected TFBSs for each bin and for each library (130 sequences per library). These TFBSs covered a wide range of regulation strengths determined from sort-seq data. The horizontal axis of each panel shows the regulation strengths of the 130 binding sites as quantified through sort-seq. The vertical axis shows the fluorescence levels of the same binding sites, as quantified by a plate reader (arbitrary units). The top of each panel shows the result of a test of the null hypothesis that the Pearson correlation coefficient R between the two quantities is equal to zero, together with a 95% confidence interval. The shaded grey area surrounding the regression line indicates the 95% confidence interval, i.e., the probability that the true linear regression line of the population lies within the confidence interval of the regression line calculated from the sample data at a confidence level of 95%. R^2^ indicates the coefficient of determination. Each circle in the plot represents the mean of three independent (biological) replicate fluorescence measurements in a plate reader after eight hours of growth. Circle colors indicate GFP fluorescence from low (dark green) to high (light green **a. Association between plate reader fluorescence and regulation strengths for the CRP library.** R = -0.81, P < 0.001; R² = 0.66, P < 0.001, N= 130. **b. Association between plate reader fluorescence and regulation strengths for the Fis library**. R = -0.74, P < 0.001; R² = 0.54, P < 0.001, N=130. **c. Association between plate reader fluorescence and regulation strengths for the IHF library.** R = -0.73, P < 0.001; R² = 0.53, P < 0.001 N = 130

**Supplementary Figure S12.**
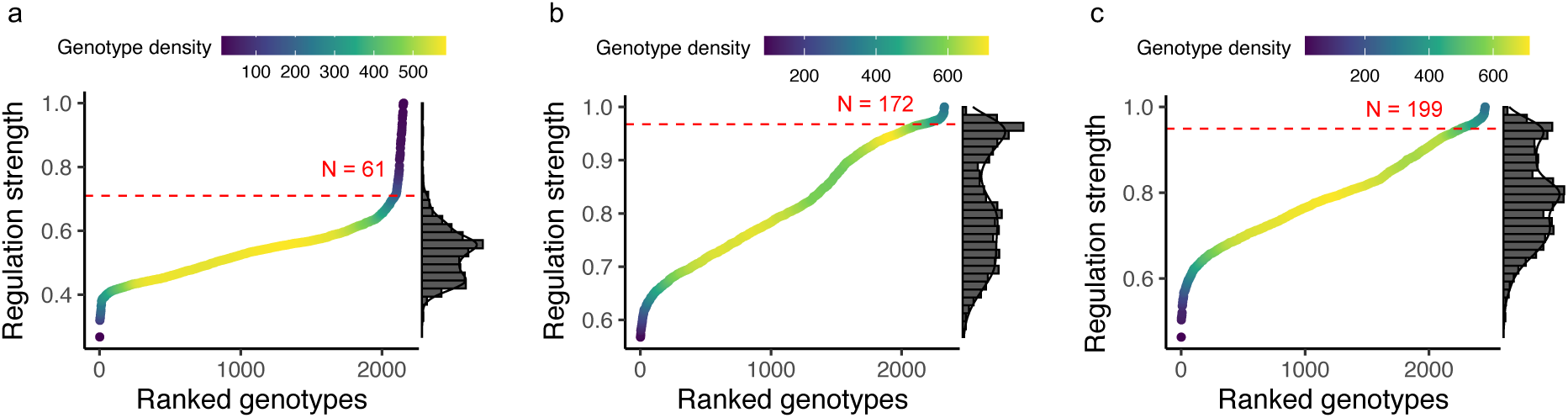
Adaptive peaks and their regulation strengths across TF landscapes. **a. Adaptive peaks in the CRP landscape.** The distribution of regulation strengths across the landscape’s 2,154 adaptive peaks is shown. The WT’s regulation strength (0.71) is marked by a horizontal dashed red line for reference. A total of N=61 peaks have higher regulation strength than the WT. The color gradient indicates genotype density along the regulation strength axis, with the histogram on the right detailing the regulation strength distribution. **b)** like a) but for the **Fis landscape,** based on its 2,312 adaptive peaks. (WT regulation strength: 0.97) N=172 peaks regulate expression more strongly than the WT**. c)** like b) but for the IHF landscape, based on its 2,434 adaptive peaks. (WT regulation strength: 0.95) N=199 peaks regulate expression more strongly than the WT.

**Supplementary Figure S13.**
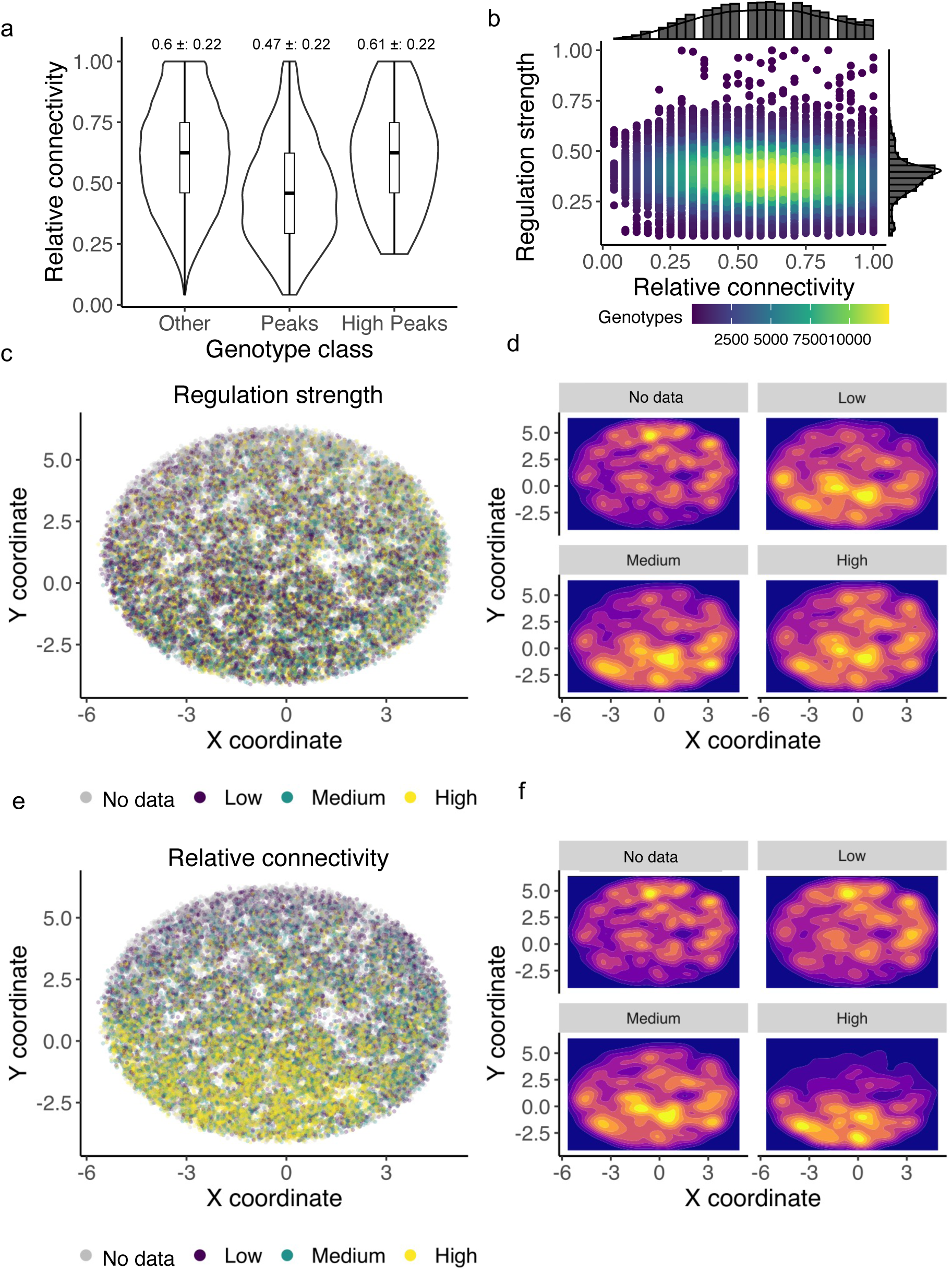
Quantitative analysis of CRP landscape sparsity. **a. Distribution of relative connectivity among genotypes.** Violin plots augmented with embedded boxplots illustrate the distribution of relative connectivity, i.e,. the ratio of the empirically observed number of adjacent genotypes to the theoretical maximum number of 24(=8×3) within a fully connected network. Genotypes are stratified into the three categories “other“ (non-peak genotypes), “peaks” (excluding high peaks), and “high peaks”. Mean relative connectivities and standard deviations, are shown above each categorical division, revealing lower relative connectivity in peaks relative to non-peak genotypes (Two-sample Welch’s t-test, p-value = 4.59 × 10^-77^, N1= 29,821; N2 = 2,093). High peaks exhibit comparable or superior connectivity relative to non-peak genotypes (Two-sample Welch’s t-test, p-value = 1, N1 = 29,821; N2 = 61). **b. Weak association between connectivity and regulation strength.** Scatterplot of relative connectivity against regulation strength. The density of genotypes (circles) in the scatterplot area is represented by a color gradient from purple (low density) to yellow (high density). The weak association (Pearson correlation, R = -0.13; t = -23.888; degrees of freedom (df) = 31,973; p-value < 2.2 × 10^-16^) indicates that strongly regulating genotypes dare not necessarily densely connected. Marginal histograms adjacent to the scatterplot quantify the distributions of regulation strength and relative connectivity. **c. Visualization of CRP landscape based on regulation strength.** Visual overview of the CRP landscape with genotypes color-coded according to regulation strength (low strength: purple; medium: green; high: yellow; grey: genotypes with missing data). To enhance clarity, edges between genotypes are not shown. The landscape is projected onto a 2D space through a force-directed algorithm^197^, with axes representing arbitrary units. **d. Spatial distribution of regulation strengths.** Each panel shows a density contour plot of the distribution of regulation strength scores of genotypes within one of four regulation strength categories (no data [missing data], low, medium, high). It indicates the density of genotypes within each of the four categories through a color gradient from blue (low density) to yellow (high density). **e. Visualization of CRP landscape based on relative connectivity.** Analogous to panel c, but for relative connectivity. Color-codes indicate relative connectivity (see color legend). **f. Spatial distribution of relative connectivity.** Like panel d, but for four categories of relative connectivity (no data, low, medium, high).

**Supplementary Figure S14.**
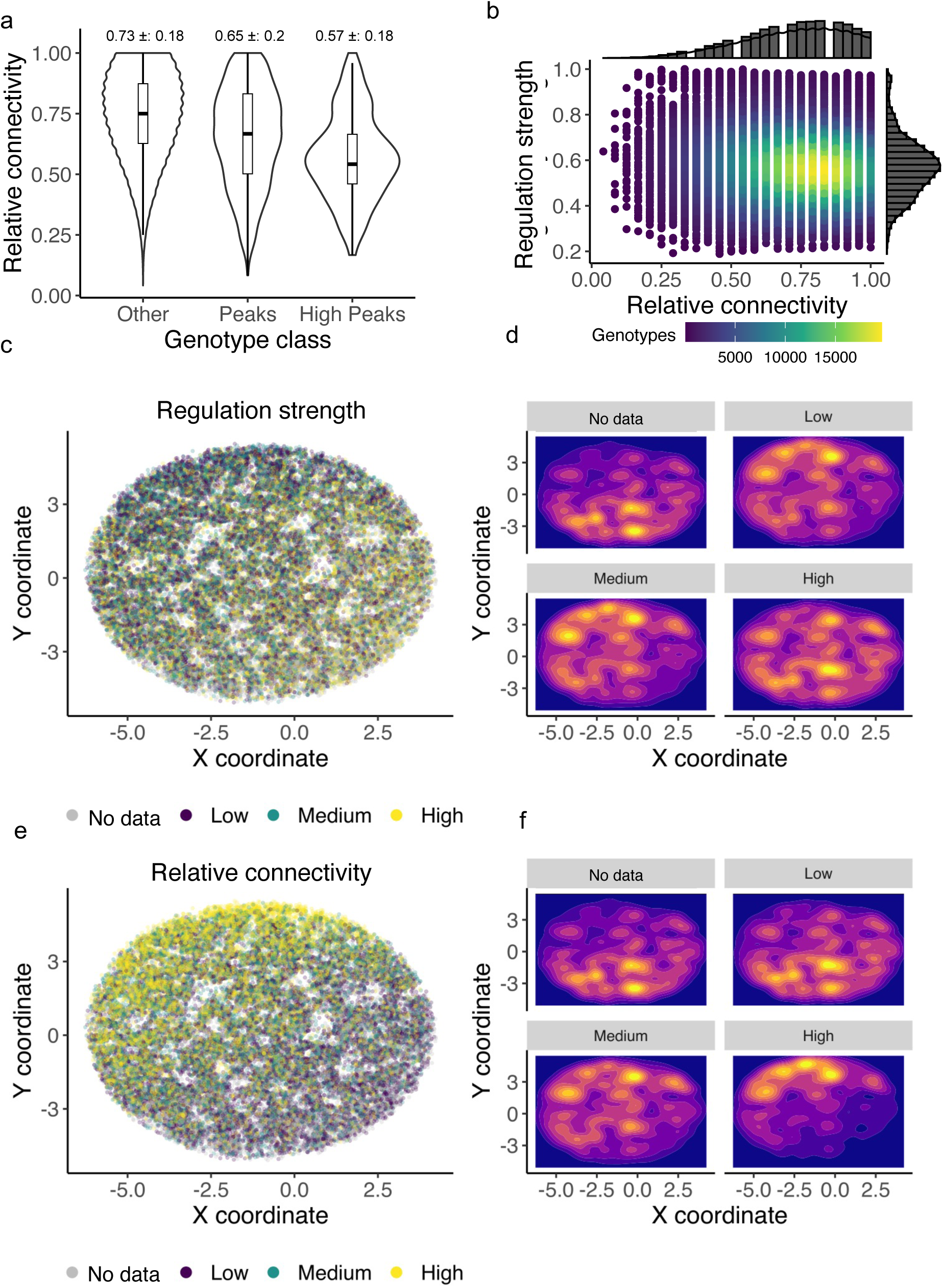
Quantitative analysis of the Fis landscape sparsity. **a. Distribution of relative connectivity among genotypes.** Violin plots augmented with embedded boxplots illustrate the distribution of relative connectivity, i.e., the ratio of the empirically observed number of adjacent genotypes to the theoretical maximum number of 24 (=8×3) within a fully connected network. Genotypes are stratified into the three categories “other” (non-peak genotypes), “peaks” (excluding high peaks), and “high peaks.” Mean relative connectivities and standard deviations are shown above each categorical division, revealing lower relative connectivity in peaks relative to non-peak genotypes (Two-sample Welch’s t-test, p-value = 4.59 × 10^-77^, N1 = 40,910; N2 = 2,312). High peaks exhibit comparable or superior connectivity relative to non-peak genotypes (Two-sample Welch’s t-test, p-value = 9.31 × 10^-13^, N1 = 40,910; N2 = 172). **b. Weak association between connectivity and regulation strength.** Scatterplot of relative connectivity against regulation strength. The density of genotypes (circles) in the scatterplot area is represented by a color gradient from purple (low density) to yellow (high density). The weak association (Pearson correlation, R = -0.13; t = -23.888; degrees of freedom (df) = 31,973; p-value < 2.2 × 10^-16^) indicates that strongly regulating genotypes are not necessarily densely connected. Marginal histograms adjacent to the scatterplot quantify the distributions of regulation strength and relative connectivity. **c. Visualization of Fis landscape based on regulation strength.** Visual overview of the Fis landscape with genotypes color-coded according to regulation strength (low strength: purple; medium: green; high: yellow; grey: genotypes with missing data). To enhance clarity, edges between genotypes are not shown. The landscape is projected onto a 2D space through a force-directed algorithm ^197^, with axes representing arbitrary units. **d. Spatial distribution of regulation strengths.** Each panel shows a density contour plot of the distribution of regulation strength scores of genotypes within one of four regulation strength categories (no data [missing data], low, medium, high). The density of genotypes within each category is indicated by a color gradient from blue (low density) to yellow (high density). **e. Visualization of Fis landscape based on relative connectivity.** Analogous to panel c, but for relative connectivity. Color codes indicate relative connectivity (see color legend). **f. Spatial distribution of relative connectivity.** Similar to panel d, but for four categories of relative connectivity (no data, low, medium, high).

**Supplementary Figure S15.**
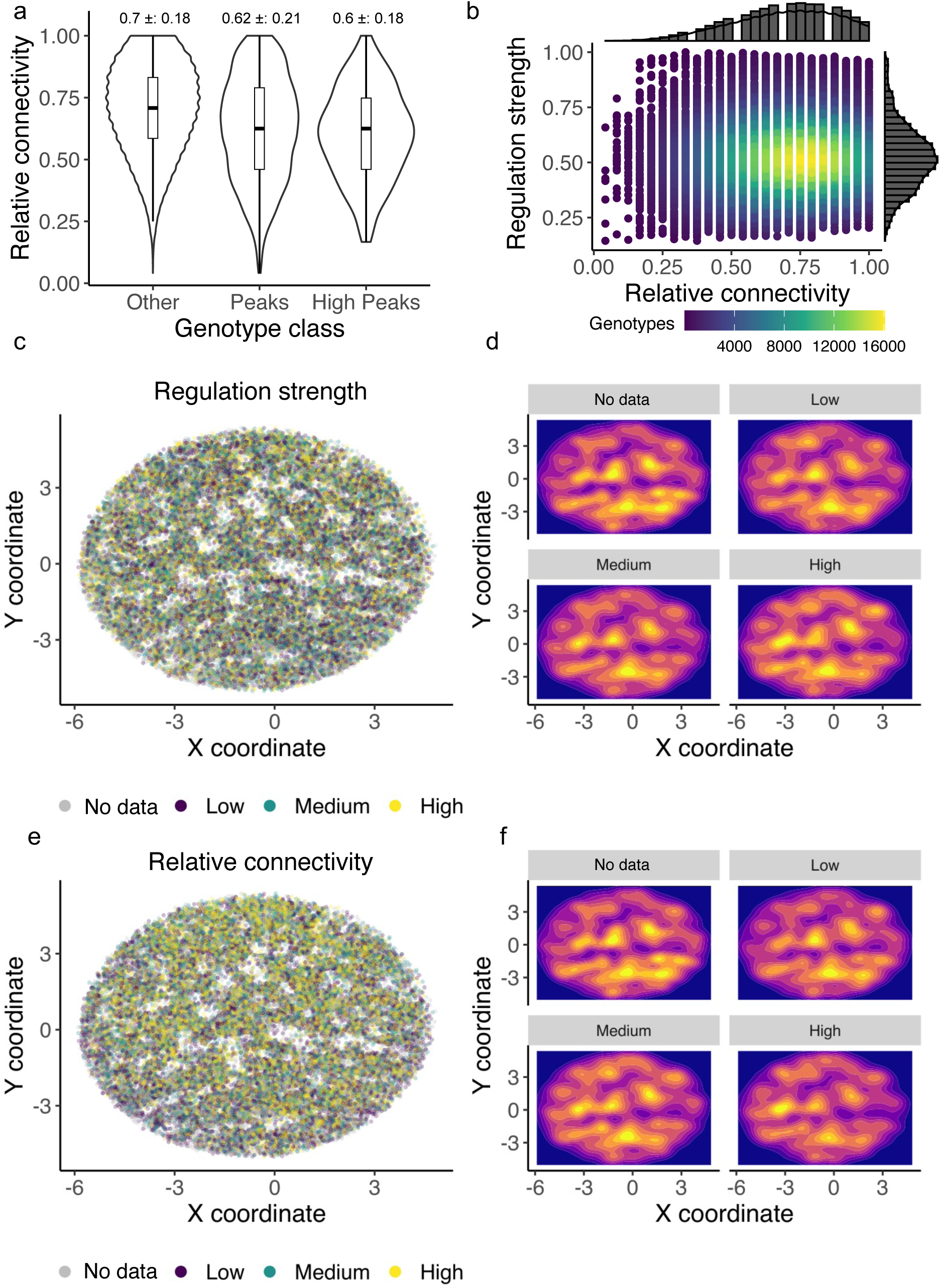
Quantitative analysis of the IHF landscape sparsity. **a. Distribution of relative connectivity among genotypes.** Violin plots augmented with embedded boxplots illustrate the distribution of relative connectivity, i.e., the ratio of the empirically observed number of adjacent genotypes to the theoretical maximum number of 24 (=8×3) within a fully connected network. Genotypes are stratified into the three categories “other” (non-peak genotypes), “peaks” (excluding high peaks), and “high peaks.” Mean relative connectivities and standard deviations are shown above each categorical division, revealing lower relative connectivity in peaks relative to non-peak genotypes (Two-sample Welch’s t-test, p-value = 1.14 × 10^-77^, N1 = 38,872; N2 = 2,254). High peaks exhibit comparable or superior connectivity relative to non-peak genotypes (Two-sample Welch’s t-test, p-value = 9.31e-13, N1 = 38,872; N2 = 199). **b. Weak association between connectivity and regulation strength.** Scatterplot of relative connectivity against regulation strength. The density of genotypes (circles) in the scatterplot area is represented by a color gradient from purple (low density) to yellow (high density). The slight positive correlation (Pearson correlation, R = 0.017; t = 3.51; degrees of freedom (df) = 41,323; p-value = 0.0004485) indicates that genotypes with elevated regulation strength are only marginally more likely to be densely connected. Marginal histograms adjacent to the scatterplot quantify the distributions of regulation strength and relative connectivity. **c. Visualization of IHF landscape based on regulation strength.** Visual overview of the IHF landscape with genotypes color-coded according to regulation strength (low strength: purple; medium: green; high: yellow; grey: genotypes with missing data). To enhance clarity, edges between genotypes are not shown. The landscape is projected onto a 2D space through a force-directed algorithm ^197^, with axes representing arbitrary units. **d. Spatial distribution of regulation strengths.** Each panel shows a density contour plot of the distribution of regulation strength scores of genotypes within one of four regulation strength categories (no data [missing data], low, medium, high). The density of genotypes within each category is indicated by a color gradient from blue (low density) to yellow (high density). **e. Visualization of IHF landscape based on relative connectivity.** Analogous to panel c, but for relative connectivity. Color codes indicate relative connectivity (see color legend). **f. Spatial distribution of relative connectivity.** Similar to panel d, but for four categories of relative connectivity (no data, low, medium, high).

**Supplementary Figure S16.**
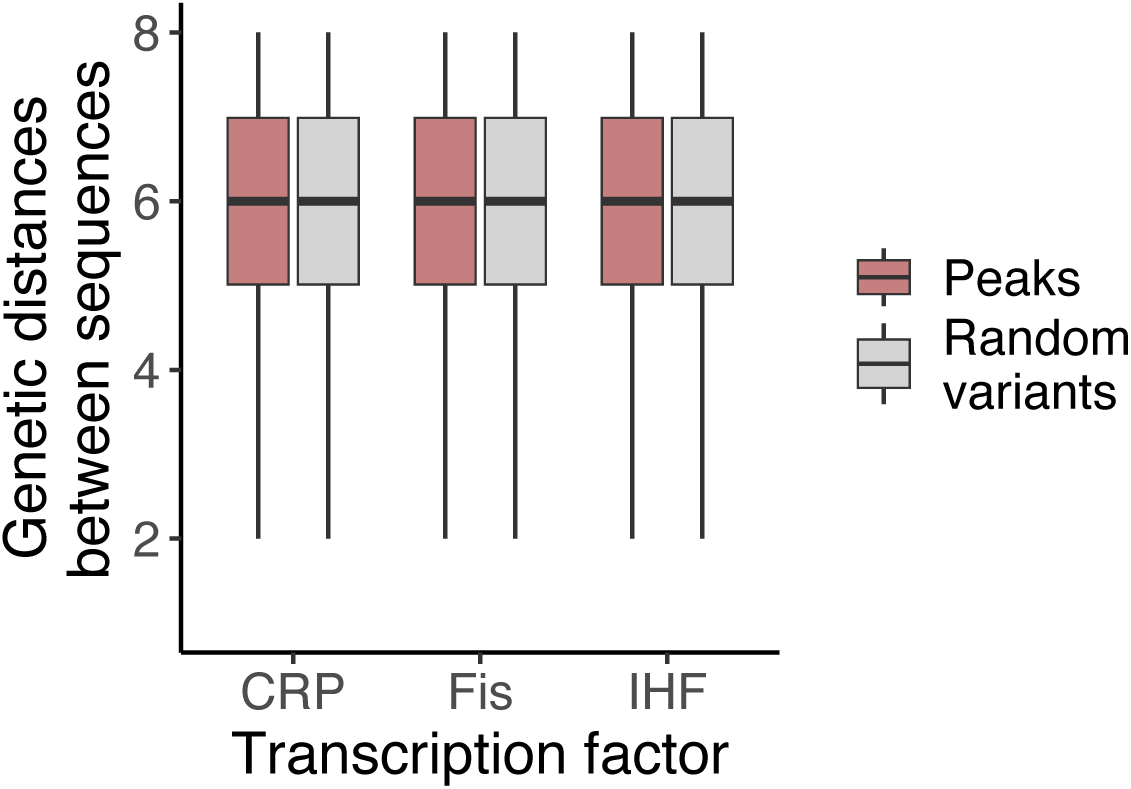
Peaks are spread across each landscape. The vertical axis shows the genetic distance *d* between peaks and an equal number of randomly chosen non-peak genotypes (color legend) for the CRP (N= 2,154), Fis (N= 2,312), and IHF (N= 2,434) landscape (horizontal axis). (CRP: d_peak_= 6, d_random_= 5.94, t-statistic= 52.49, p-value <0.01; Fis: d_peak_ = 6, d_random_ = 5.97, t-statistic= 30.96, p-value <0.01; IHF: d_peak_ = 6, d_random_ = 5.97, t-statistic= 34.07, p-value <0.01; null-hypothesis: means do not differ). Although the differences are statistically significant, peaks are almost as dispersed within genotype space as non-peaks. Each box covers the range between the first and third quartiles (IQR). The horizontal line within the box represents the median value, and whiskers span 1.5 times the IQR.

**Supplementary Figure S17.**
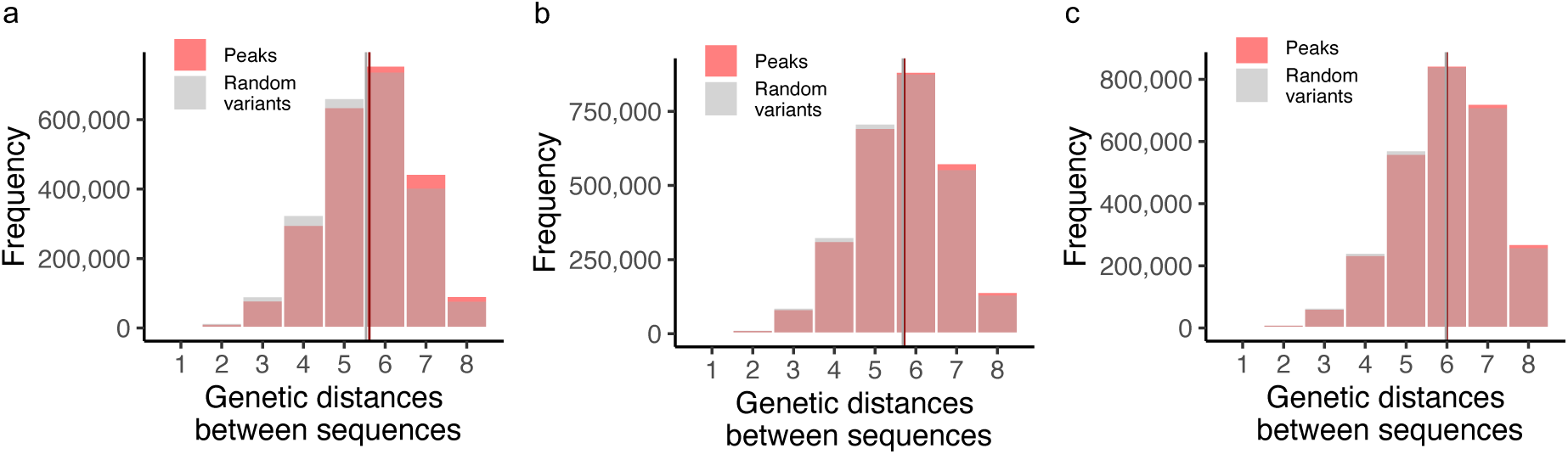
Genetic diversity among peaks across TF landscapes. **a. CRP landscape.** Comparative analysis of genetic distances among nucleotide sequences from 2,154 peaks (red) versus 2,154 non-peak, random variants (grey). Mean genetic distances are displayed with vertical lines, showing a slight difference between peaks (d=5.6, in red) and non-peaks (d=5.5, in grey), demonstrating large genetic diversity within the peaks (Welch Two Sample t-test, t = 78.299, df = 4637305, p-value < 2.2 × 10^-16^). **b. Fis landscape.** Comparative analysis of genetic distances among nucleotide sequences from 2,312 peaks (red) versus 2,312 non-peak, random variants (grey). Mean genetic distances are displayed with vertical lines, showing no significant difference between peaks (d=5.7, in red) and non-peaks (d=5.7, in grey), demonstrating large genetic diversity within the peaks (t = 27.234, df = 5412460, p-value < 2.2 × 10^-16^). **c. IHF landscape.** Comparative analysis of genetic distances among nucleotide sequences from 2,434 peaks (red) versus 2,434 non-peak, random variants (grey). Mean genetic distances are displayed with vertical lines, showing no significant difference between peaks (d=6, in red) and non-peaks (d=6, in grey), demonstrating large genetic diversity within the peaks (t = 54.689, df = 6014743, p-value < 2.2 × 10^-16^).

**Supplementary Figure S18.**
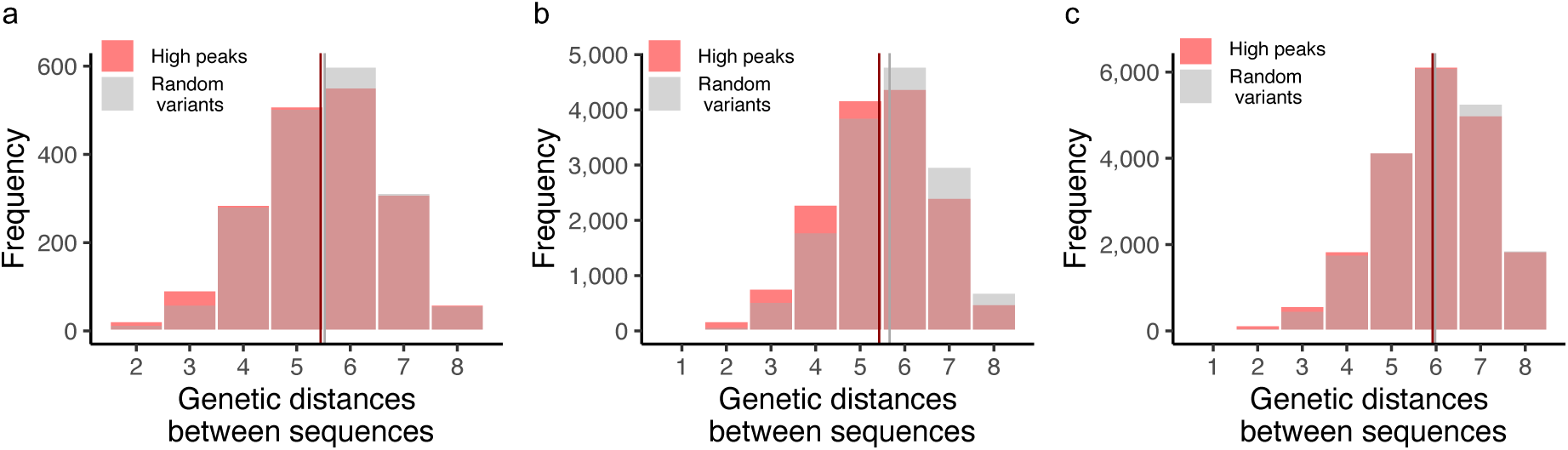
Genetic diversity among high peaks across TF landscapes. **a. CRP landscape.** The overlayed histograms compare the genetic diversity between 61 high regulation strength peaks (red) and 61 non-peak, randomly selected variants (grey). No significant difference exists (W) between high-regulation strength peaks (mean d=5.4, in red) compared to randomly selected genotypes (mean d=5.5, in grey; Welch Two Sample t-test, t = -0.78555, df = 3653.7, p-value = 0.4322). **c. Fis landscape.** The overlayed histograms compare the genetic diversity between 172 high-regulation strength peaks (red) and 172 non-peak, randomly selected genotypes (grey). No significant difference exists between high-regulation strength peaks (mean d=5.4, in red) and random genotypes (mean d=5.8, in grey; Welch Two Sample t-test, t = -16.634, df = 29357, p-value < 2.2 × 10^-16^). **c. IHF landscape.** The overlayed histograms compare the genetic diversity between 199 high regulation strength peaks (red) and 199 non-peak, randomly selected genotypes (grey). The analysis reveals no significant difference between high regulation strength peaks (mean d=5.8, in red) compared to the random variants (mean d=5.8, in grey; Welch Two Sample t-test, t = 0.73365, df = 39366, p-value = 0.4632).

**Supplementary Figure S19.**
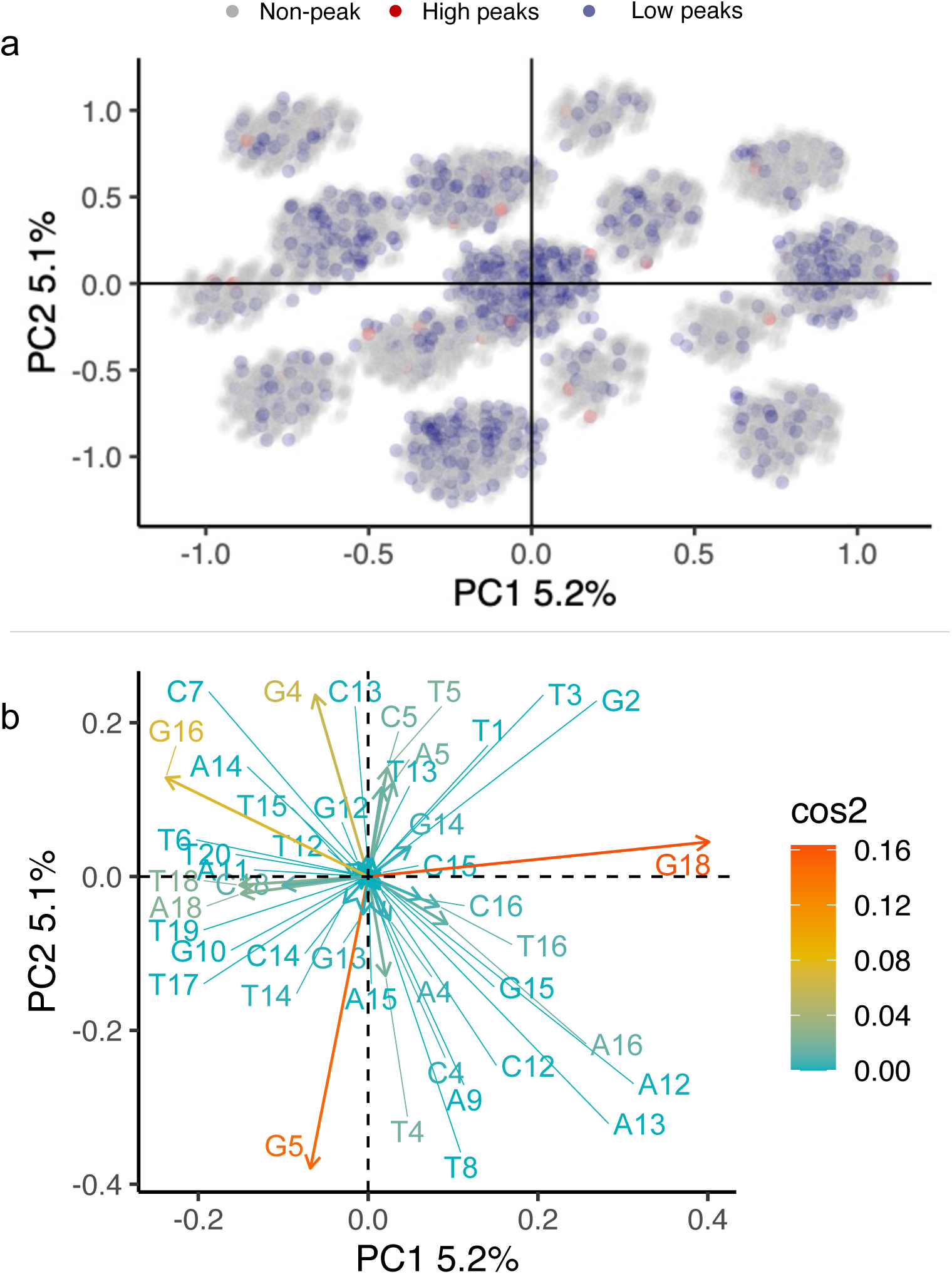
Principal component analysis of the CRP landscape. **a. PCA plot.** A principal component analysis reveals that adaptive peaks are distributed widely within the genotype space (Supplementary Methods). The panels show principal components (PC) 1 and 2. Axes labels also indicate the percentage of variation explained by each component. Every circle corresponds to one among the 31,975 genotypes within the landscape. Light blue and light red circles represent low and high peaks, respectively, while grey circles represent non-peak variants. The clustered structure in the genotype space is due to the different influence of individual nucleotides on the variation observed in the data (see panel **b**). **b. The contribution of individual nucleotides to principal component analysis.** A PCA contribution plot is a way to visualize the relative importance of different variables to the variation observed in the data. In this plot, the contribution of each variable (nucleotide identity at each position of a TFBS) is expressed as the square of the cosine of the angle (cos^2^) between the variable’s vector (column representing the presence/absence of a base letter at each position in the sequence) and each principal component axis. This quantity is represented as an arrow that indicates the correlation of the variable with PC1 and PC2, the two principal components that capture the largest amount of variation in the data. Both length and color of the arrow represent the contribution of the variable to the variation observed in the data. A high cos^2^ value (red) indicates that the variable is strongly correlated with the principal component, and therefore makes a large contribution to the variation observed in the data. A low cos^2^ value (blue) indicates that the variable is weakly correlated with the principal component, and therefore makes a small contribution to the variation observed in the data. Each arrow’s length represents the importance of the variable’s contribution relative to other variables in the plot. Each nucleotide is represented with a base letter (A, T, C, G) followed by a number that indicates the position of that base in the binding site sequence (e.g. G18 stands for a guanine at position 18 of the binding site).

**Supplementary Figure S20.**
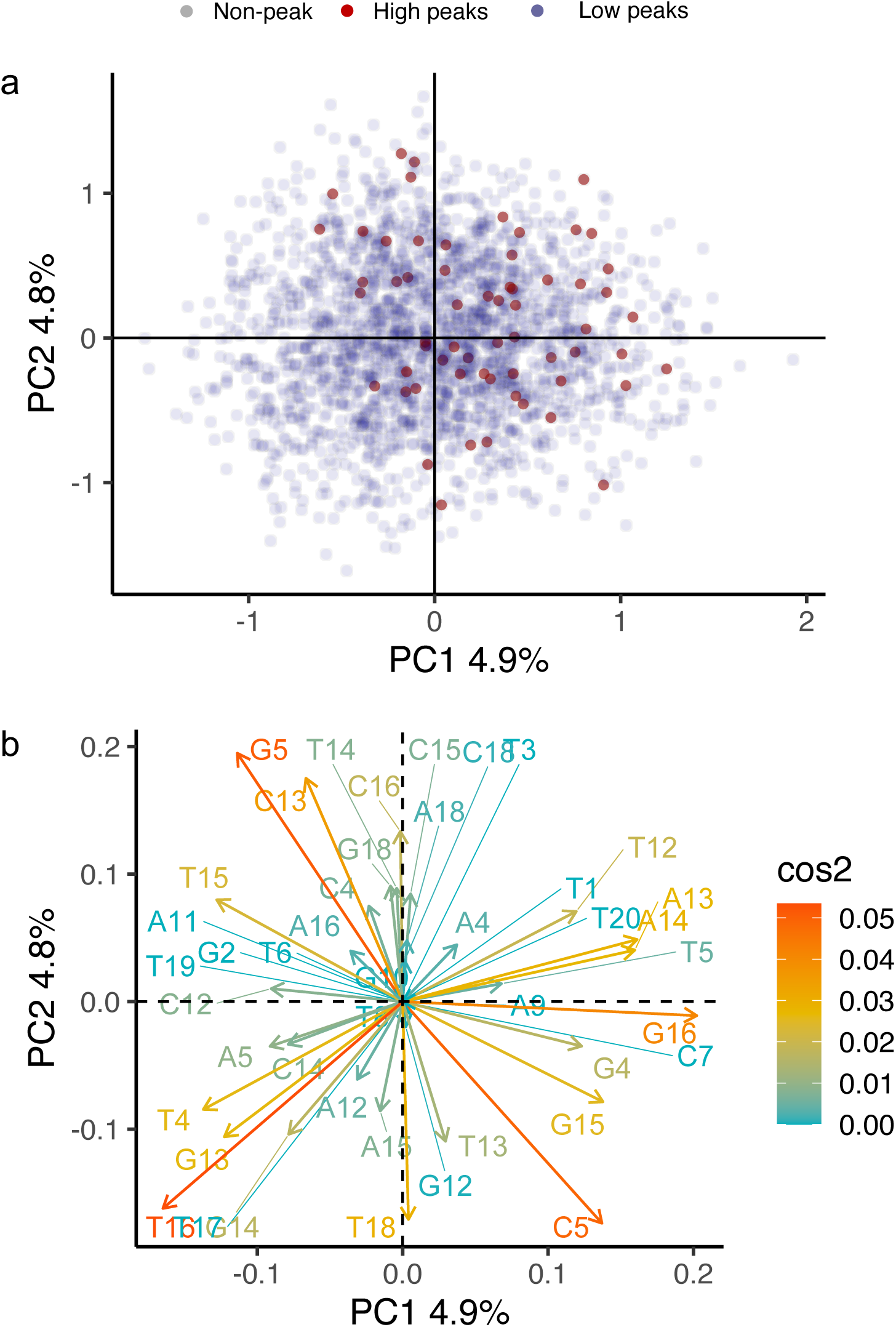
**Principal component analysis of the Fis landscape a. PCA plot**. A principal component analysis reveals that adaptive peaks are distributed widely within the genotype space (Supplementary Methods). The panels show principal components (PC) 1 and 2. Axes labels also indicate the percentage of variation explained by each component. Every circle corresponds to one among the 43,222 genotypes within the landscape. Light blue and light red circles represent low and high peaks, respectively, while grey circles represent non-peak variants. The clustered structure of the data in genotype space is due to the different influence of individual nucleotides on the variation observed in the data (see panel b). **b. The contribution of individual nucleotides to principal component analysis**. A PCA contribution plot visualizes the relative importance of different variables to the variation observed in the data. In this plot, the contribution of each variable (nucleotide identity at each position of a TFBS) is expressed as the square of the cosine of the angle (cos^2^) between the variable’s vector (column representing the presence/absence of a base letter at each position in the sequence) and each principal component axis. This quantity is represented as an arrow that indicates the correlation of the variable with PC1 and PC2, the two principal components that capture the largest amount of variation in the data. Both the length and color of the arrow represent the contribution of the variable to the variation observed in the data. A high cos^2^ value (red) indicates that the variable is strongly correlated with the principal component and therefore makes a large contribution to the variation observed in the data. A low cos^2^ value (blue) indicates that the variable is weakly correlated with the principal component and therefore makes a small contribution to the variation observed in the data. Each arrow’s length represents the importance of the variable’s contribution relative to other variables in the plot. Each nucleotide is represented with a base letter (A, T, C, G) followed by a number that indicates the position of that base in the binding site sequence (e.g., G18 stands for a guanine at position 18 of the binding site).

**Supplementary Figure S21.**
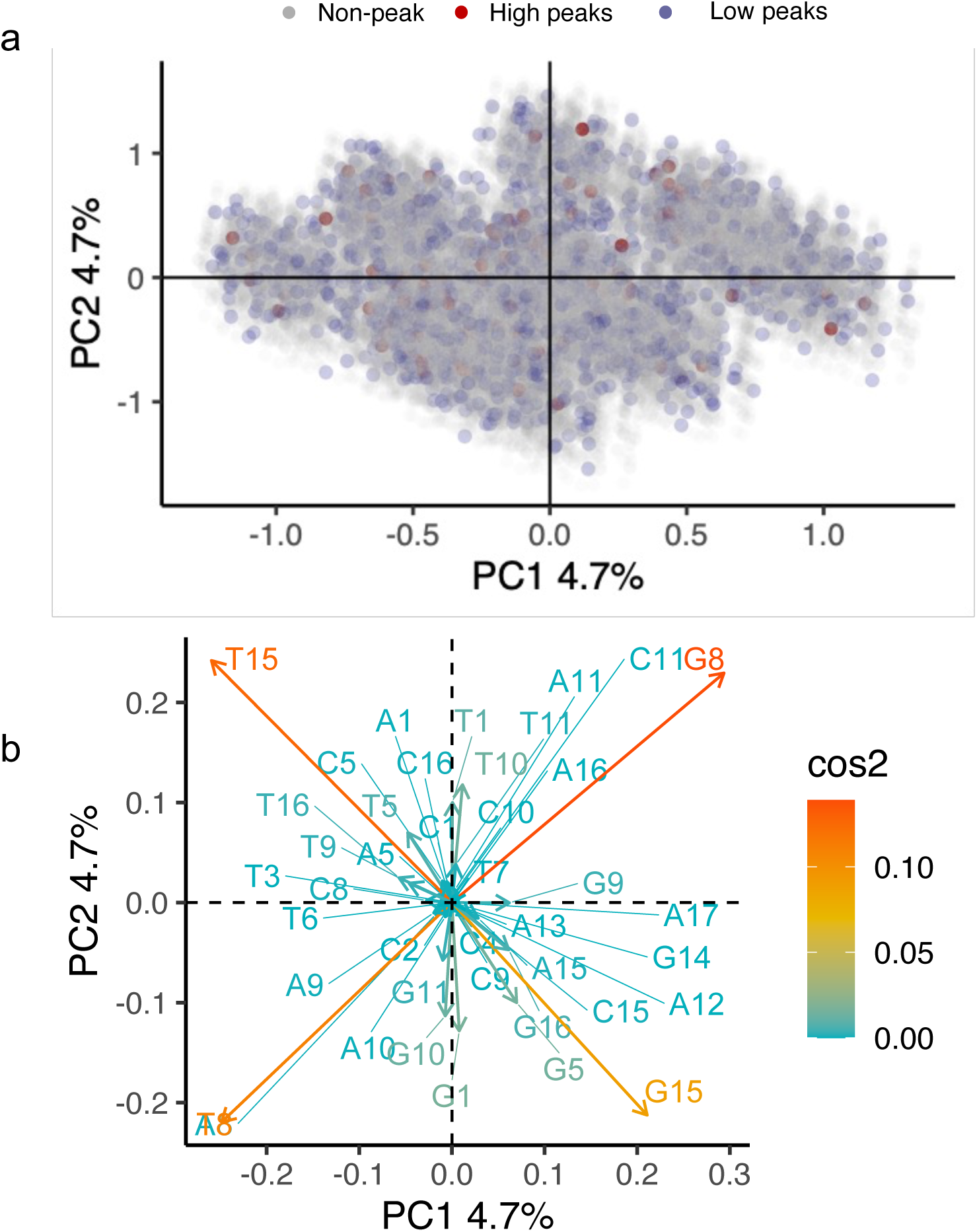
**Principal Component Analysis of the IHF Landscape a. PCA Plot**. A principal component analysis reveals that adaptive peaks are distributed widely within the genotype space (Supplementary Methods). The panels show principal components (PC) 1 and 2. Axes labels also indicate the percentage of variation explained by each component. Every circle corresponds to one among the 41,325 genotypes within the landscape. Light blue and light red circles represent low and high peaks, respectively, while grey circles represent non-peak variants. The clustered structure in the genotype space is due to the different influence of individual nucleotides on the variation observed in the data (see panel b). **b. The contribution of individual nucleotides to principal component analysis**. A PCA contribution plot is a way to visualize the relative importance of different variables to the variation observed in the data. In this plot, the contribution of each variable (nucleotide identity at each position of a TFBS) is expressed as the square of the cosine of the angle (cos^2^) between the variable’s vector (column representing the presence/absence of a base letter at each position in the sequence) and each principal component axis. This quantity is represented as an arrow that indicates the correlation of the variable with PC1 and PC2, the two principal components that capture the largest amount of variation in the data. Both the length and color of the arrow represent the contribution of the variable to the variation observed in the data. A high cos^2^ value (red) indicates that the variable is strongly correlated with the principal component and therefore makes a large contribution to the variation observed in the data. A low cos^2^ value (blue) indicates that the variable is weakly correlated with the principal component and therefore makes a small contribution to the variation observed in the data. Each arrow’s length represents the importance of the variable’s contribution relative to other variables in the plot. Each nucleotide is represented with a base letter (A, T, C, G) followed by a number that indicates the position of that base in the binding site sequence (e.g., G18 stands for a guanine at position 18 of the binding site).

**Supplementary Figure S22.**
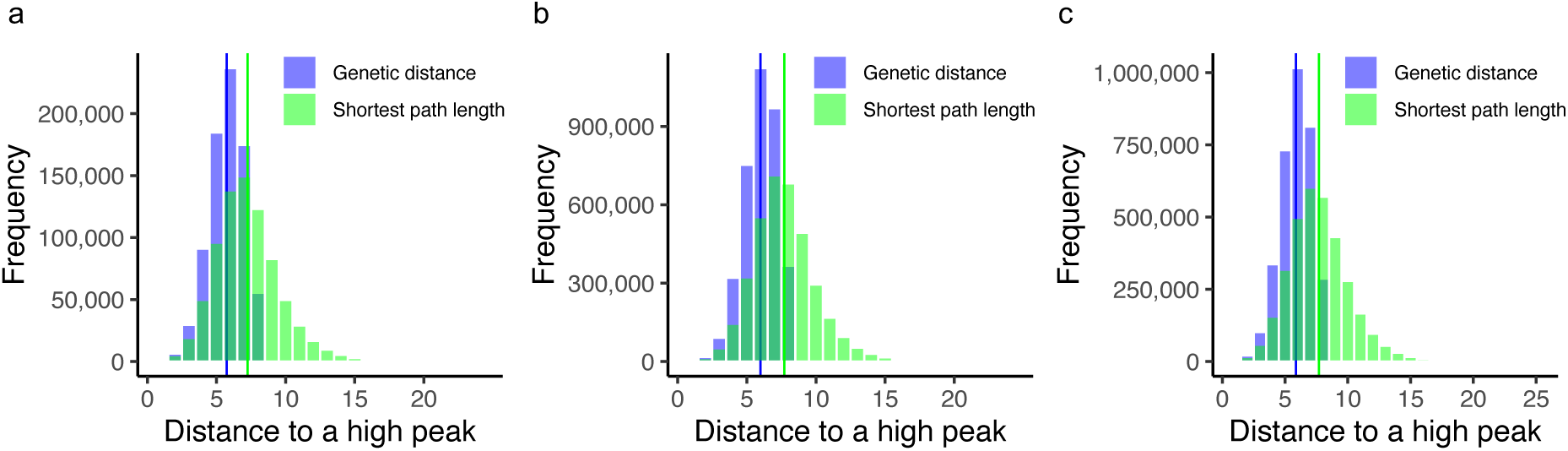
Accessible paths to high peaks are less than two mutational steps longer than the shortest genetic distances to the peak. Each composite histogram shows the distribution of genetic distances (blue) and shortest accessible path length (green) for all pairs of high peaks and non-peak genotypes for one of the three landscapes. We used a t-test of the null hypothesis that the means of these distance distributions are statistically indistinguishable. This null hypothesis is rejected for all three landscapes. **a. CRP Landscape.** Mean genetic distance: 5.74; mean number of mutational steps in shortest accessible paths: 7.2; t = 500.17, df = 1,212,712, p < 2.2 × 10^-16^. **b. Fis Landscape.** Mean genetic distance: 5.98; mean number of mutational steps in shortest accessible paths: 7.7; t = -1271, df = 5,639,001, p < 2.2 × 10^-16^. **c. IHF Landscape.** Mean genetic distance: 5.78; mean number of mutational steps in shortest accessible paths: 7.6; t = -1365, df = 6,636,001, p < 2.2 × 10^-16^.

**Supplementary Figure S23.**
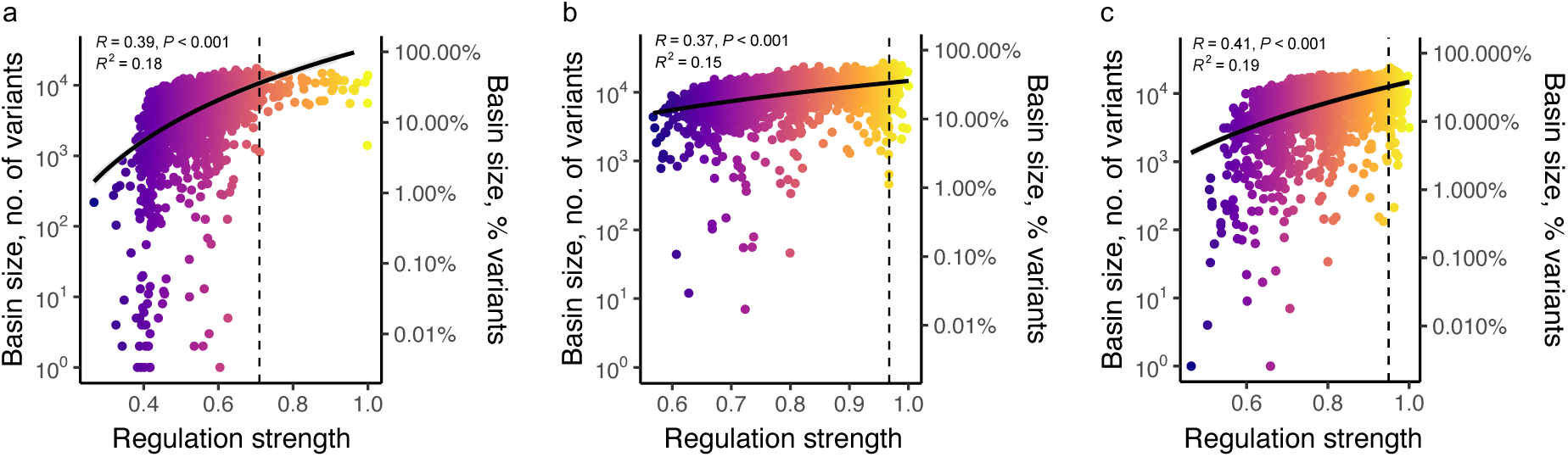
Moderate positive correlation between peak regulation strength and basin size for the three regulatory landscapes. **a. CRP landscape.** The scatter plot shows the association between regulation strength conveyed by a peak variant (horizontal axis, colors from dark blue [low] to yellow [high]) and the size of its basin of attraction (absolute numbers on the left vertical axis, percentages on the right vertical axis, note the logarithmic scales). The dashed vertical line represents the regulation strength of the wild type. The black curve represents a linear regression line (in linear space). *R* is the Pearson correlation coefficient (R=0.39) and *R*^2^ is the coefficient of determination for the linear regression model (*R*^2^ =0.18, N = 2,154) **b. Fis landscape.** Like panel *a* (R=0.37, *R*^2^ =0.15, N = 2,312), **c. IHF landscape.** Like panel *a* (R=0.41, *R*^2^ =0.19, N = 2,434).

**Supplementary Figure S24.**
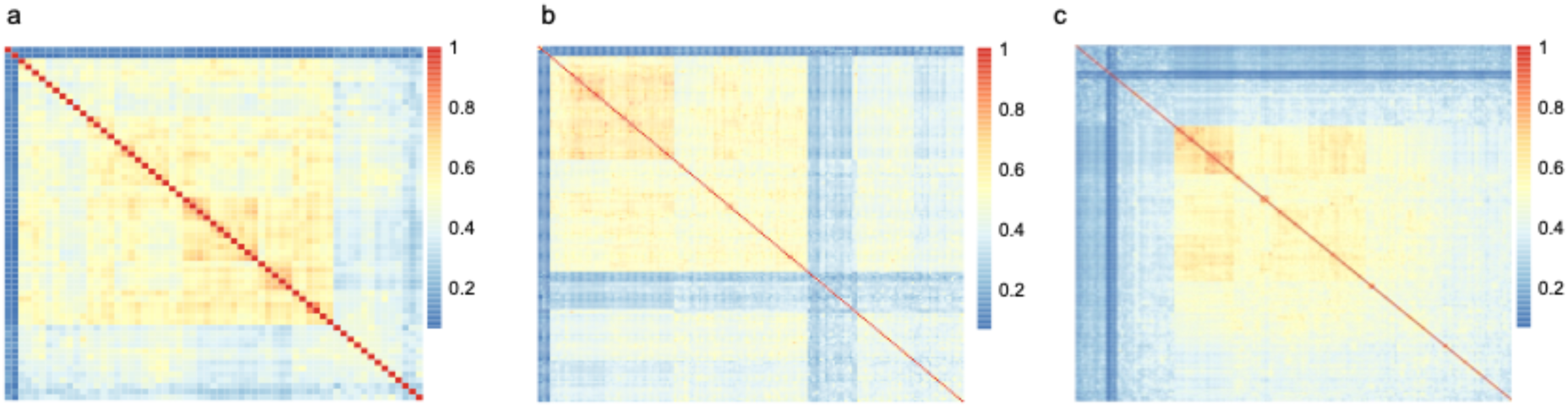
The basins of attractions of high peaks overlap. Each heatmap matrix quantifies the proportion of shared TFBS variants in the basins of attractions of all pairs of high peaks (Methods). Each matrix entry quantifies the basin overlap for one pair of high peaks using the Jaccard index (**Supplementary Methods 7.5**), where row and column indices correspond to specific pairs of peaks. The maximum overlap of one (red) signifies that two basins share all variants, while the minimum value of zero (blue) indicates that they share no variants. **a. Basin overlap in the CRP landscape.** The average Jaccard index is 0.461 with a standard deviation of ±0.144, based on 3,721 pairwise comparisons. **b. Basin overlap in the Fis landscape**. The average Jaccard index is 0.426 with a standard deviation of ±0.136, derived from 29,584 pairwise comparisons. **c. Basin overlap in the IHF landscape.** The average Jaccard index is 0.364, with a standard deviation of ±0.145, based on 39,601 pairwise comparisons.

**Supplementary Figure S25.**
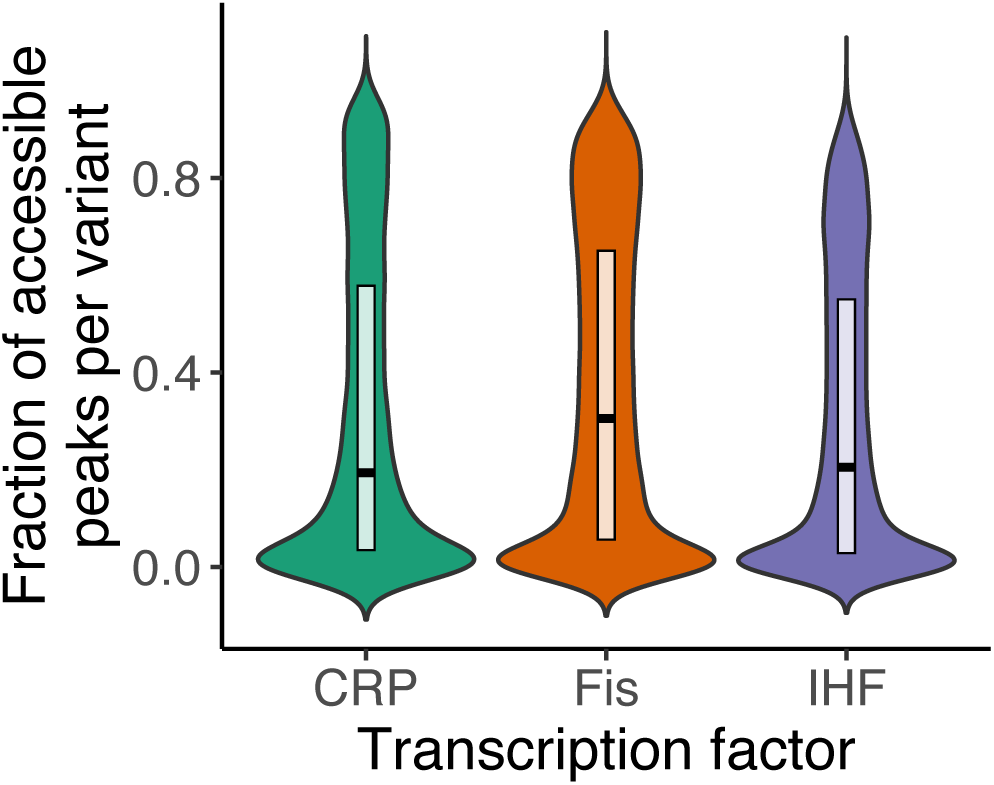
Distribution of the fraction of accessible high peaks per TFBS variant across TF landscapes. The figure shows violin plots with embedded boxplots, for the distribution of high peak accessibility of each variant within the CRP (green), Fis (orange), and IHF (purple) landscape. The y-axis quantifies peak accessibility as the fraction of the total number of high peaks that can be accessed from each (non-peak) genotype within each TF landscape, revealing the extent to which individual variants can potentially access different high peaks. In the CRP landscape, each genotype can access, on average, a proportion of 0.314±0.315 (mean±s.d.) high peaks from the total number of high peaks, with a median value of 0.194. Notably, 5,152 out of 29,821 genotypes (17%) can access no high peak. Fis: mean high peak accessibility: 0.363±0.311 (median: 0.305), with 4,304 out of 40,895 (10%) genotypes lacking access to any high peaks. IHF: mean peak accessibility: 0.301±0.289 (median: 0.205); 4,629 out of 38,872 genotypes (12%) show lack access to high peaks.

**Supplementary Figure S26.**
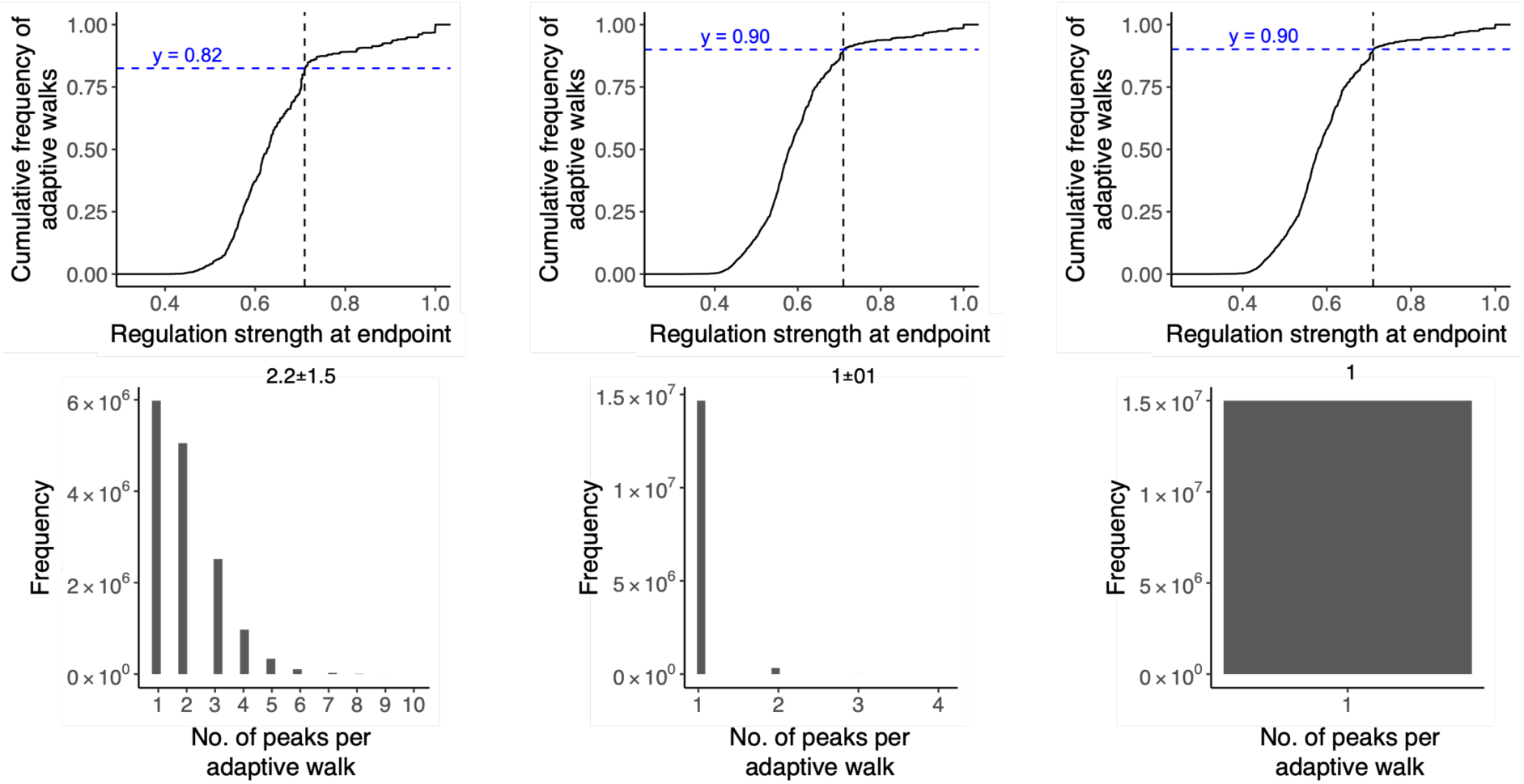
Kimura adaptive walks on the CRP landscape a-c. Fraction of adaptive walks attaining TFBS variants with strong regulation for different population sizes. The vertical axis indicates the cumulative fraction of adaptive walks on the CRP landscape that end at a genotype with a regulation strength shown on the horizontal axes. Data is based on 15 million simulated adaptive walks (15,000 random starting genotypes × 1,000 adaptive walks each). The vertical dashed line indicates the regulation strength of the wild-type TFBS for CRP, and can be used to infer the proportion of walks that terminate at lower regulatory peaks (Horizontal line, blue lettering). For example, a value of y=0.82 implies that 82% of walks terminate at peaks lower than the wild-type. The remaining 18% reach higher peaks. **a. Population size 10**^2^. 18% of walks terminate at peaks higher than the wild-type. **b. Population size 10**^5^. 10% of the walks terminate at higher peaks. **c. Population size 10**^8^. 10% of walks terminate at higher peaks. **d-f. Number of high peaks reached by a single adaptive walks.** The histograms show the distribution of the number of high peaks visited by a single adaptive walk, based on the same 15 × 10^6^ adaptive walks analyzed in a-c. Note that a walk can visit multiple peaks when genetic drift is strong, because genetic drift can displace a population from an adaptive peak. The mean ± s.d. of the distribution is shown at the top of each plot. Population sizes: **d** 10^2^**, e** 10^5^, and **f** 10^8^.

**Supplementary Figure S27.**
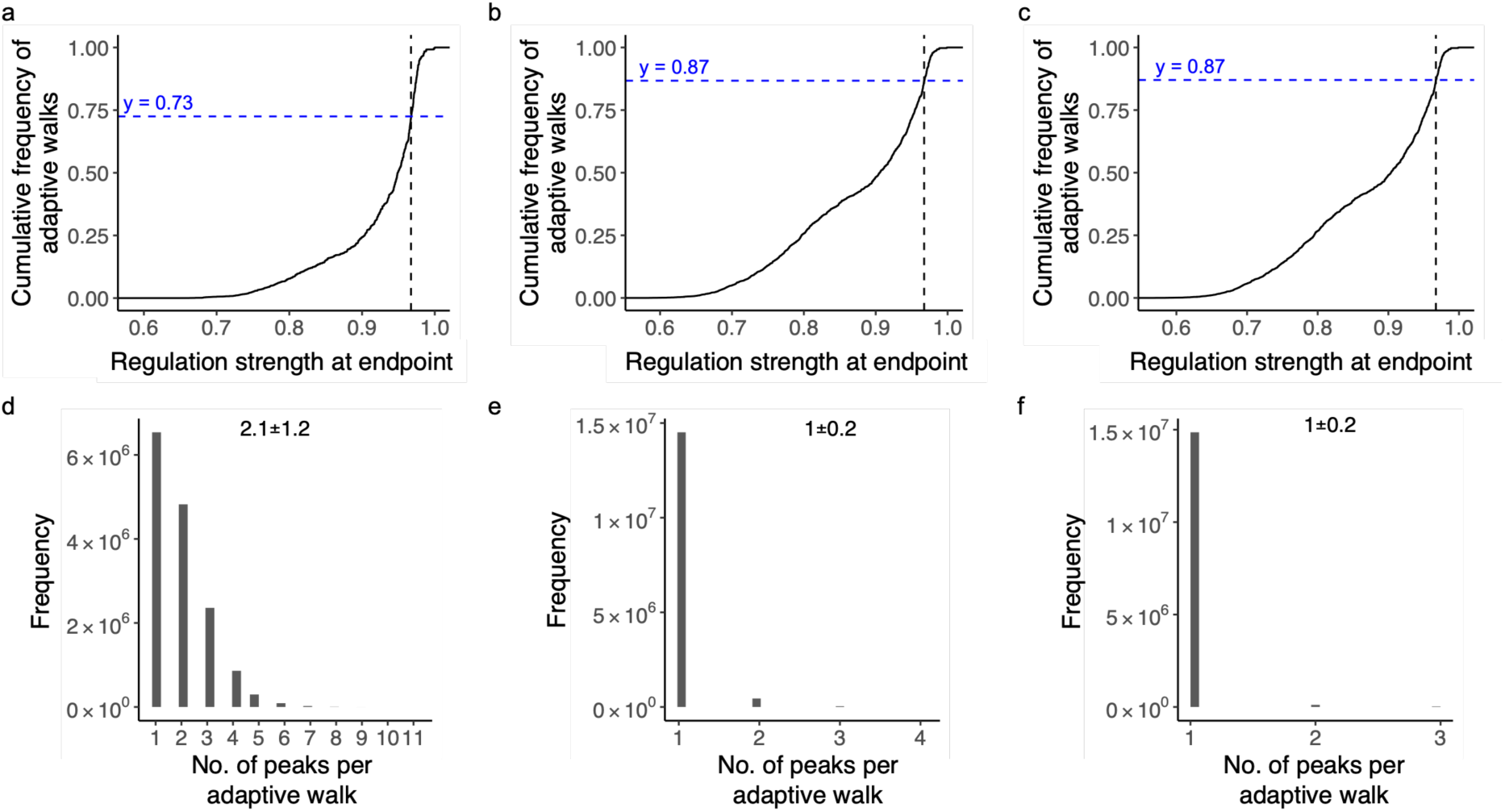
Kimura adaptive walks on the Fis landscape. **a-c. Fraction of adaptive walks attaining high peaks for different population sizes.** The vertical axis indicates the cumulative percentage of adaptive walks on the Fis landscape that end at a genotype with a regulation strength shown on the horizontal axis. Data is based on 15 million simulated adaptive walks (15,000 starting genotypes × 1,000 adaptive walks each). The vertical dashed line indicates the regulation strength of the wild-type TFBS for Fis, and can be used to infer the proportion of walks that terminate at lower regulatory peaks (horizontal line, blue lettering). For example, a y-value of 0.9 implies that 90% of walks terminate at peaks lower than the wild-type, with the remaining 10% reaching higher peaks. **a. Population size 10².** 27% of walks terminate at peaks higher than the wild-type. **b. Population size 10⁵.** 13% of walks terminate at higher peaks. **c. Population size 10⁸.** 13% of walks terminate at higher peaks. **d-f. Number of high peaks reached by individual adaptive walks.** The histograms show the distribution of the number of high peaks visited by a single adaptive walk, based on the same 150 million adaptive walks analyzed in a-c. Note that a walk can visit multiple peaks when genetic drift is strong because genetic drift can displace a population from an adaptive peak. The mean ± s.d. of the distribution is shown at the top of each plot. Population sizes: **d** 10², **e** 10⁵, and **f** 10⁸.

**Supplementary Figure S28.**
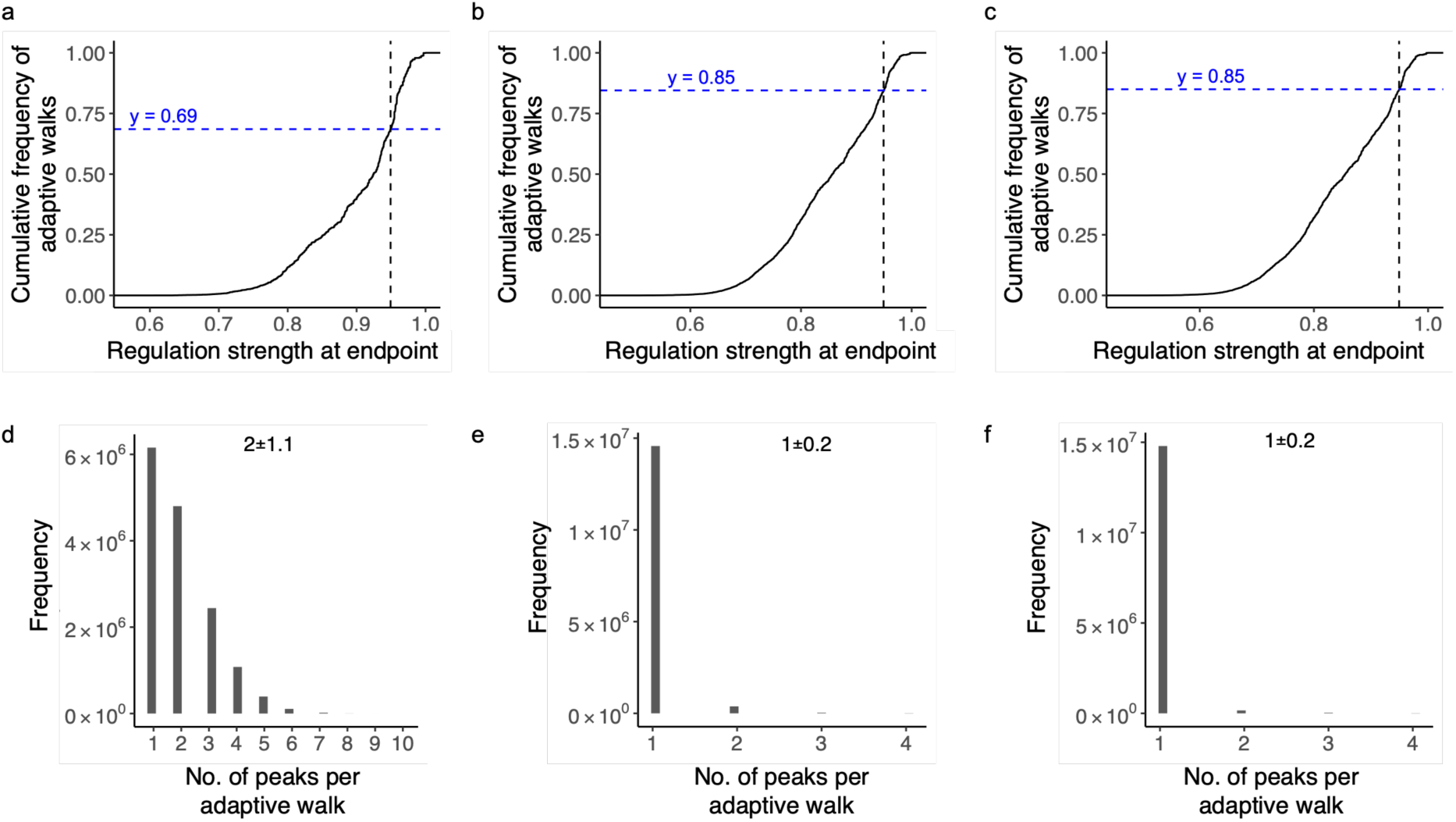
Kimura adaptive walks on the IHF landscape. **a-c. Fraction of adaptive walks attaining high peaks for different population sizes.** The vertical axis indicates the cumulative percentage of adaptive walks on the IHF landscape that end at a genotype with a regulation strength shown on the horizontal axis. Data is based on 15 million simulated adaptive walks (15,000 starting points × 1,000 adaptive walks each). The vertical dashed line indicates the regulation strength of the wild-type TFBS for IHF, and can be used to infer the proportion of walks that terminate at lower regulatory peaks (horizontal line, blue lettering). For example, a y-value of 0.9 implies that 90% of walks terminate at peaks lower than the wild-type, with the remaining 10% reaching higher peaks. **a. Population size 10².** 31% of walks terminate at peaks higher than the wild-type. **b. Population size 10⁵.** 15% of walks terminate at higher peaks. **c. Population size 10⁸.** 15% of walks terminate at higher peaks. **d-f. Number of high peaks reached by individual adaptive walks.** The histograms show the distribution of the number of high peaks visited by a single adaptive walk, based on the same 150 million adaptive walks analyzed in a-c. Note that a walk can visit multiple peaks when genetic drift is strong because genetic drift can displace a population from an adaptive peak. The mean ± s.d. of the distribution is shown at the top of each plot. Population sizes: **d** 10², **e** 10⁵, and **f** 10⁸.

**Supplementary Figure S29.**
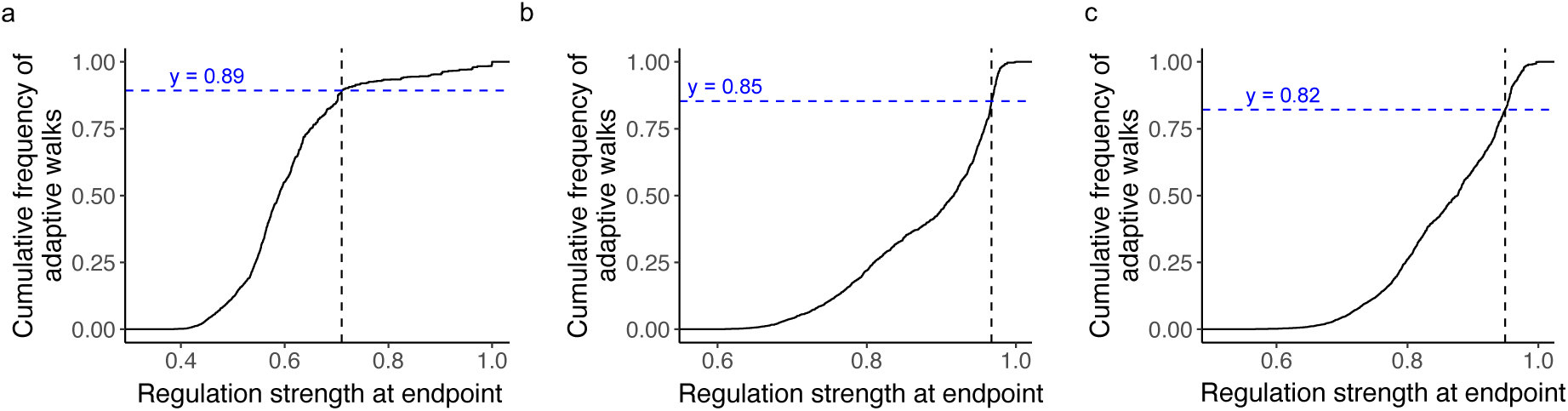
Fraction of greedy adaptive walks attaining high peaks for different TFs. **a-c. Fraction of greedy adaptive walks attaining high peaks for different TFs.** The vertical axis indicates the cumulative distribution function (CDF) of adaptive walks on each TF landscape that end at a genotype with a regulation strength shown on the horizontal axis. Data is based on 15,000 adaptive walks (15,000 starting points × 1 adaptive greedy walk for each) for each TF landscape. The vertical dashed line indicates the regulation strength of the wild-type TFBS for their respective TFs and can be used to infer the proportion of walks that terminate at lower regulatory peaks (horizontal line, blue lettering). For example, a value of y=0.9 implies that 90% of walks terminate at peaks lower than the wild-type, with the remaining 10% terminating at higher peaks. **a. CDF for adaptive walks in the CRP landscape.** Note the intersection at y= 0.89, meaning that 11% of the walks terminated on high peaks. **b. CDF for adaptive walks in the Fis landscape.** Note the intersection at y= 0.85, meaning that 15% of the walks terminated on high peaks. **c. CDF for adaptive walks in the IHF landscape.** Note the intersection at y= 0.82, meaning that 18% of the walks terminated on high peaks.

**Supplementary Figure S30.**
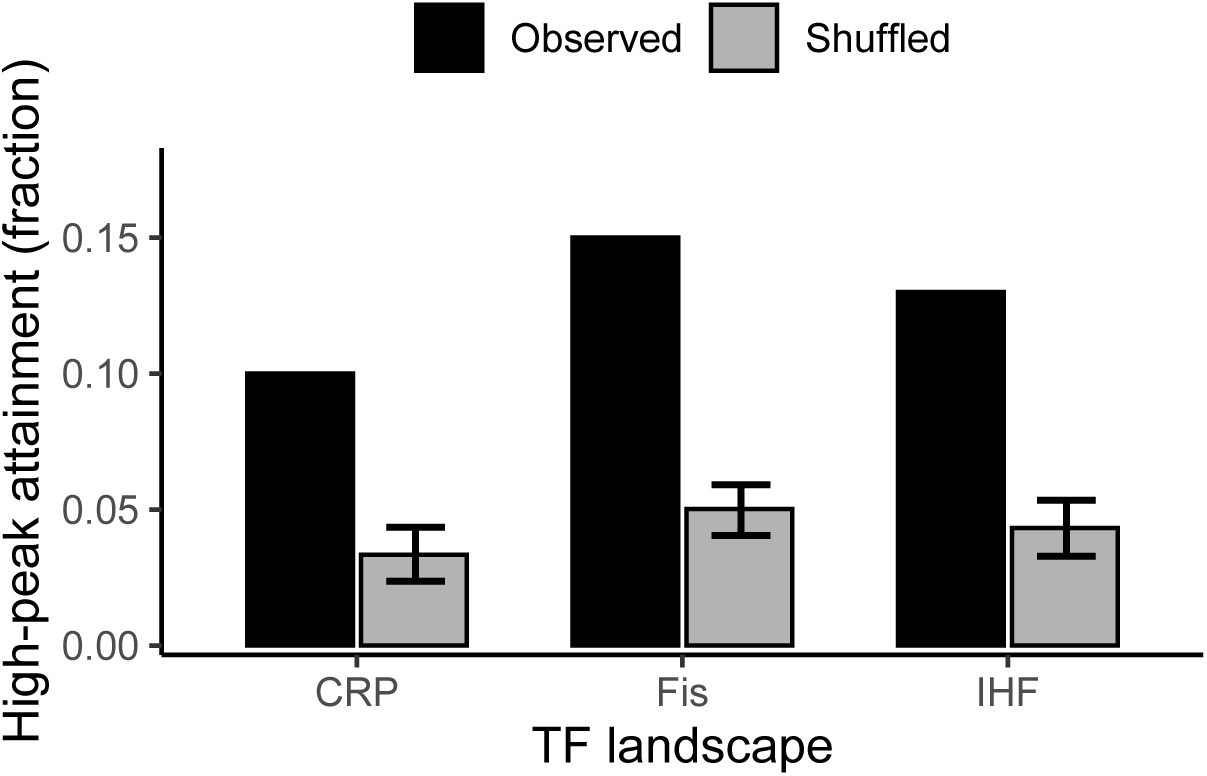
Empirical regulatory landscapes enable more frequent access to strong regulatory peaks than randomized landscapes. Bars show the fraction of adaptive walks that reach a high regulatory peak in empirical landscapes (black) and in randomized null landscapes (gray) for the transcription factors CRP, Fis, and IHF. We define high peaks as local peaks whose regulation strength exceeds that of the wild-type TFBS for the corresponding TF. We modeled adaptive evolution using Kimura walks for a population size of 10^8^. For each landscape, we simulated 10^3^ adaptive walks starting from each of 15,000 randomly and uniformly selected non-peak TFBS genotypes, allowing up to 8 mutational fixation steps per walk until a peak is reached. We quantified the null expectations from 10^3^ shuffled landscapes per TF by permuting regulation-strength values across genotypes while preserving the sampled genotype network (see **Supplementary Methods 9**). Gray bars show the mean fraction of walks reaching high peaks across shuffled landscapes; error bars indicate ± s.d. For all three TFs, empirical landscapes show significantly higher accessibility of high regulatory peaks than expected under the null model, with empirical landscapes exhibiting on average a 3-fold higher accessibility of high regulatory peaks (One-sided Monte Carlo permutation test; 𝑁_2_ = 1 empirical landscape, 𝑁_9_ = 1,000 shuffled landscapes; CRP: observed = 0.10, null = 0.035 ± 0.010, 𝑧 = 13.4, 𝑝 < 0.001; Fis: observed = 0.15, null = 0.050 ± 0.010, 𝑧 = 20.9, 𝑝 < 0.001; IHF: observed = 0.13, null = 0.045 ± 0.010, 𝑧 = 16.8, 𝑝 < 0.001; 𝑁_walks_ = 1.5 × 10:per landscape).

**Supplementary Figure S31.**
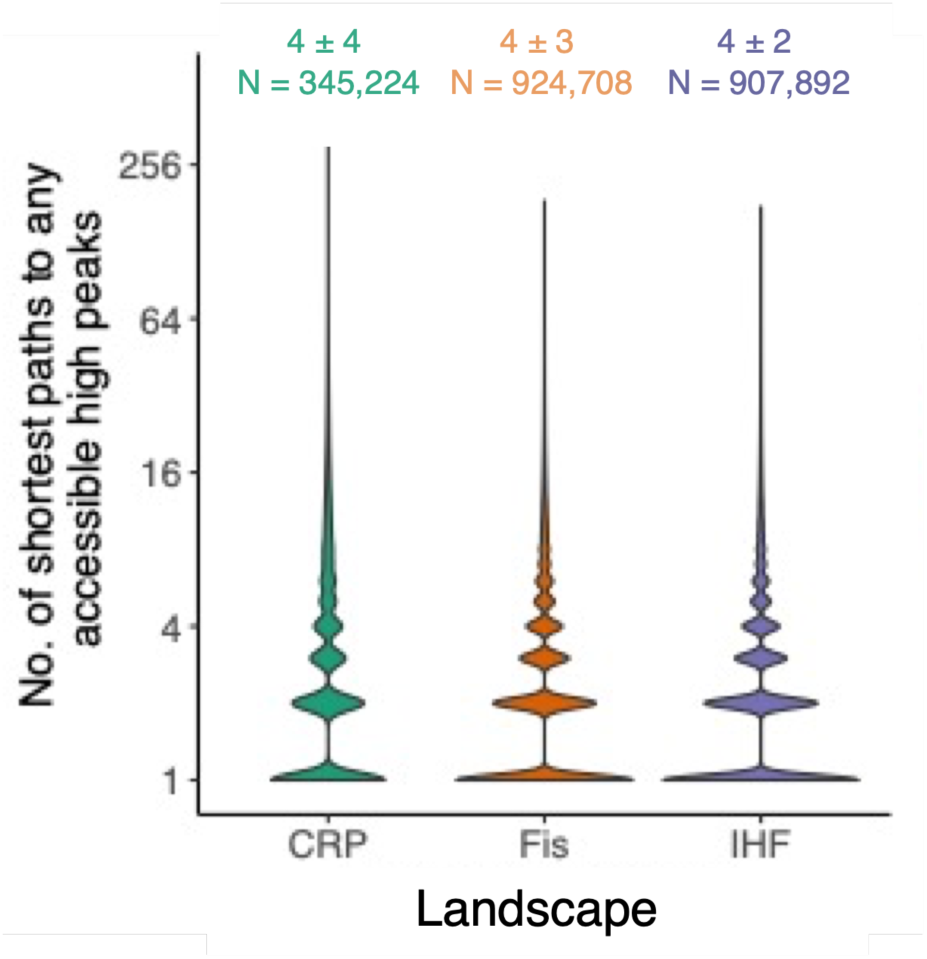
Diversity of alternative shortest paths for pairs of starting genotypes and their respective attained peaks. Each violin plot is based on data from 15,000 randomly (and uniformly) sampled starting (non-peak) genotypes from each TF landscape – CRP, Fis and IHF (see color legend) – and on 1’000 Kimura adaptive walks per starting genotypes. For each pair of a starting genotype and a high peak that is attainable from this starting genotype, we calculated the number of distinct shortest paths reaching the peak (vertical axis, note logarithmic scale). The width of the violin plot at any given y-axis value represents the density of data points at that value. Wider sections indicate a higher density of data points. The text at the top of each violin indicates the median number of shortest paths and the interquartile range (IQR) for each TF. For example, in the CRP landscape peaks are reached via a median number (± IQR) of 4 (± 4) shortest paths, with a sample size (number of starting genotypes × attainable peak pairs) of N = 345,224. The violin regions appear discretized because the number of shortest paths is a discrete variable, not a continuous one. This means that the data can only take on specific integer values, resulting in discrete regions within the violin plots with a non-zero density of data points.

**Supplementary Figure S32.**
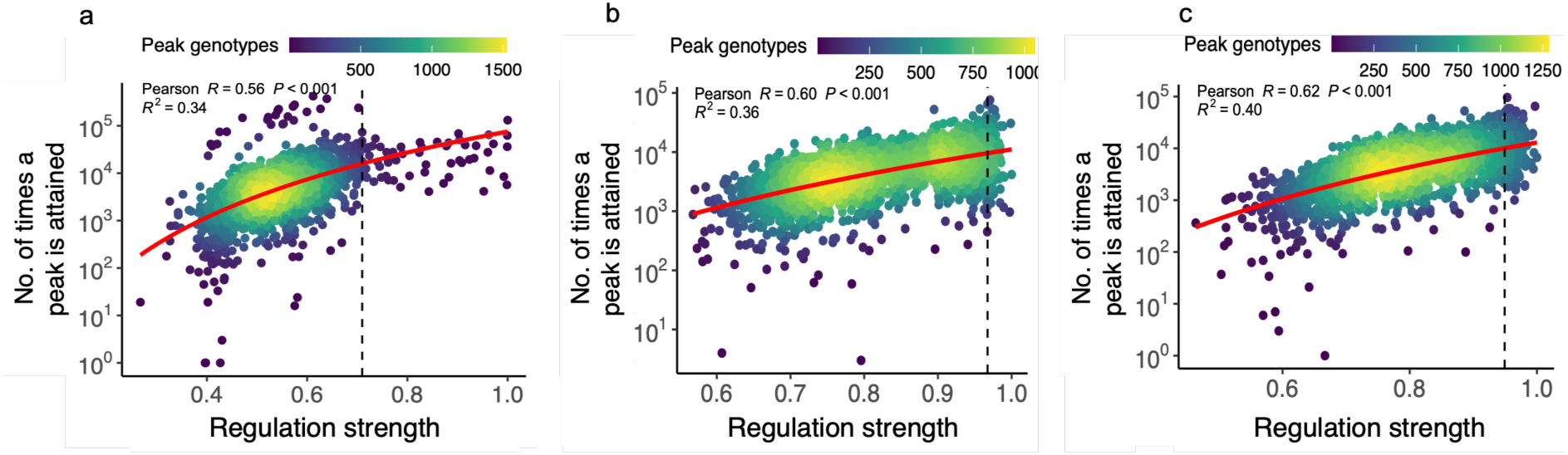
Peak genotypes conveying stronger regulation are also reached more often during adaptive evolution. Each scatter plot represents the association between the strength of regulation conveyed by a peak genotype (horizontal axis) and the number of times the peak is attained (vertical axis, note the logarithmic scale). The data for each plot is based on 15,000 randomly (and uniformly) sampled starting (non-peak) genotypes from each TF landscape – CRP, Fis and IHF (see color legend) – and on 1’000 Kimura adaptive walks per starting genotypes (15×10^6^ adaptive walks per landscape). The heatmap colors represent the density of peak genotypes from low (dark blue) to high (yellow). The dashed vertical line represents the regulation strength of the wild type TFBS. The red curve is derived from linear regression (in linear space). P-values in each panel result from a test of the null hypothesis that the two variables are not associated (Pearson correlation coefficient R=0; Coefficient of determination *R*^2^=0). **a**. **CRP landscape** (*R*= 0.56; *R*^2^ =0.34, N = 2,154). **b. Fis landscape.** (R=0.6, *R*^2^ =0.36, N = 2,312), **c. IHF landscape.** (R=0.62, *R*^2^ =0.4, N = 2,434).

**Supplementary Figures S33.**
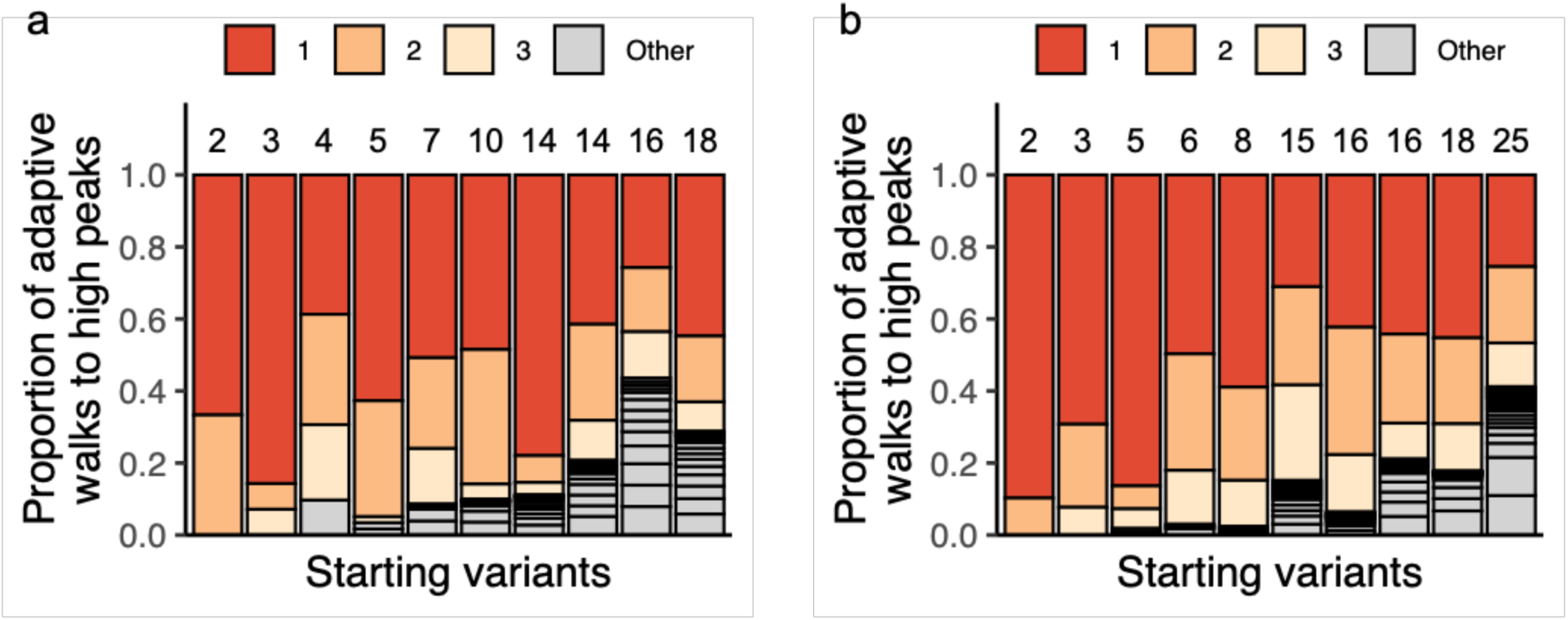
The frequency distribution of attained high peaks is biased towards some frequently attained peaks. We randomly and uniformly sampled 10 starting genotypes, started 10^3^ adaptive walks from each, and recorded the number and frequency of distinct high peaks attained in these random walks. Each starting variant is symbolized by a vertical bar. The number of stacks within each bar (delineated by horizontal lines, also indicated by an integer above each bar) shows the number of high peaks reached by the 10^3^ adaptive walks. Starting variants are ordered in ascending order based on this number of attained peaks. Stack height indicates the fraction of walks that reached the same peak, and is indicated in red, orange, and yellow for the three most frequently attained peaks. **a. Fis landscape. b. IHF landscape.**

**Supplementary Figure S34.**
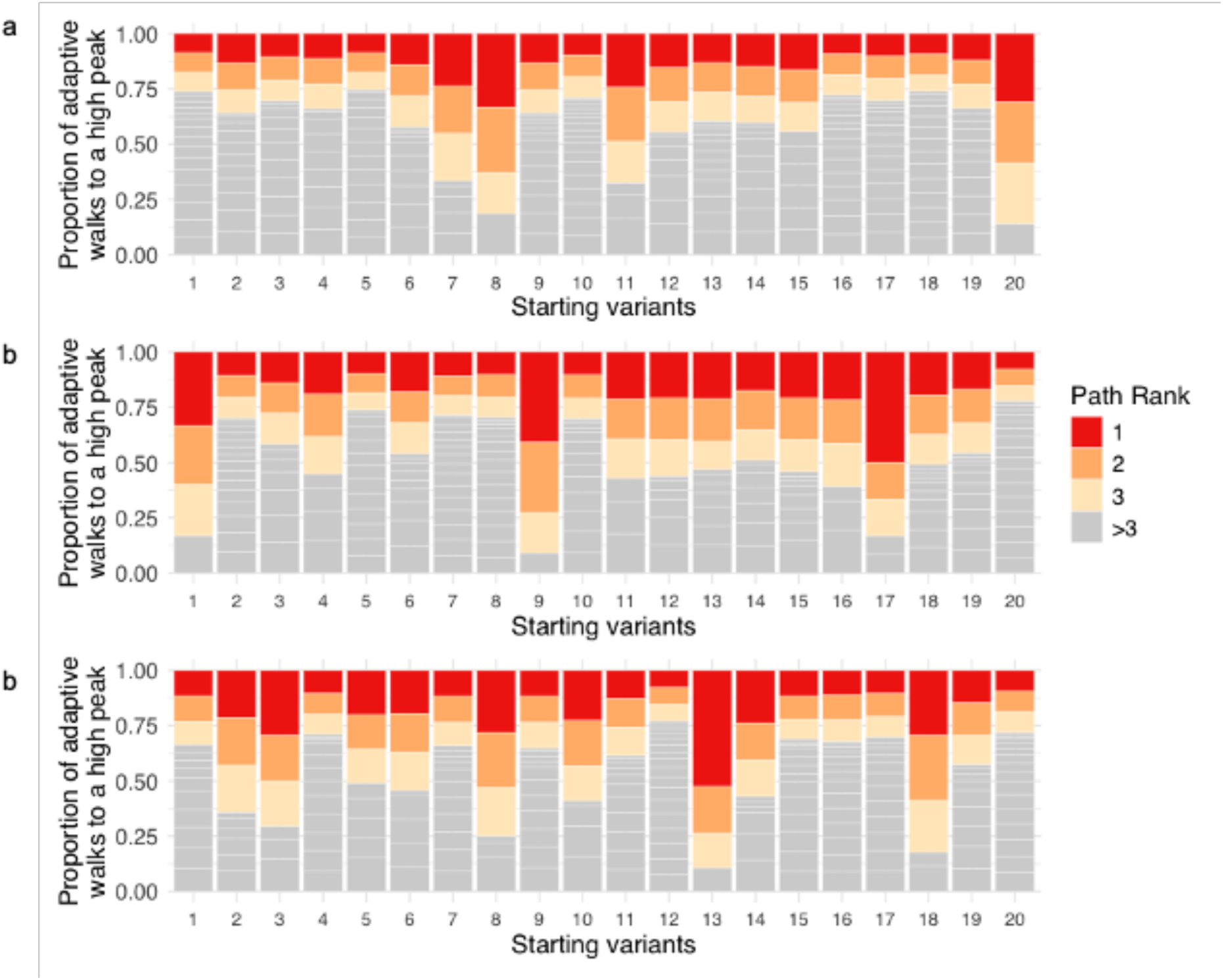
The frequency distribution of traversed paths to high peaks is biased towards some paths that are frequently traversed. We randomly and uniformly sampled 20 starting genotypes, initiated 10^3^ adaptive walks from each, and recorded the number and frequency of distinct paths traversed by these random walks for each attained peak. We then randomly and uniformly sampled a high peak attained by each starting variant for our analysis. Each starting variant is represented by a vertical bar. The number of stacks within each bar indicates the number of unique paths traversed to reach the sampled high peak. Stack height represents the fraction of walks that reached the high peak, with the three most frequently transversed paths in red, orange, and yellow. **a. CRP landscape. b. Fis landscape. c. IHF landscape.**

**Supplementary Figure S35.**
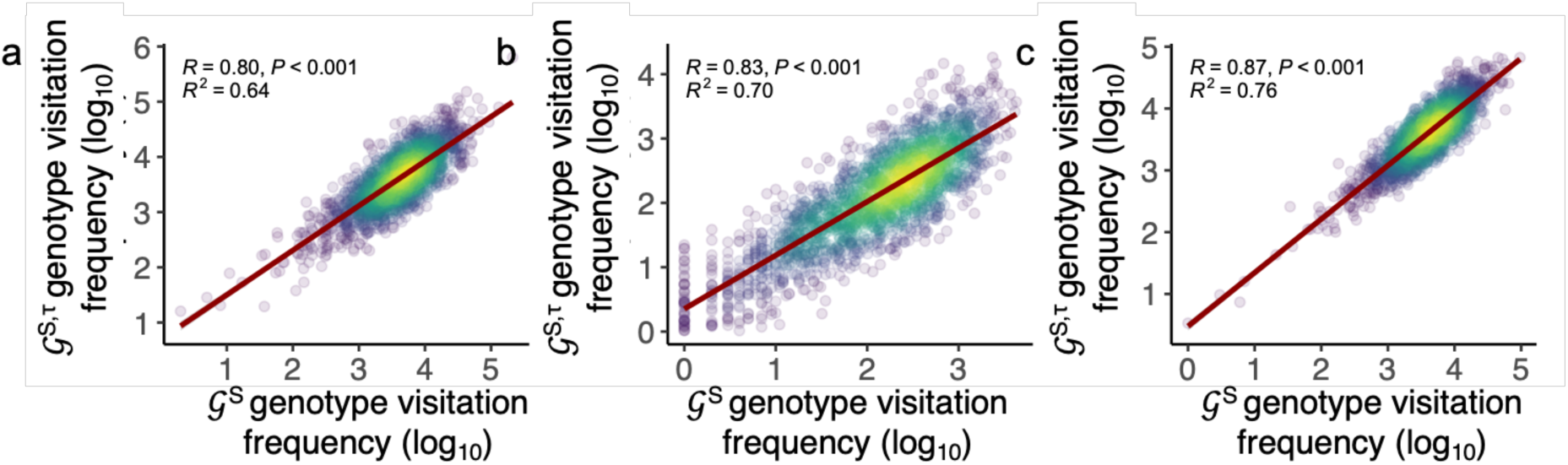
Evolutionary accessibility is robust to experimental uncertainty in regulation strength. Each panel shows a pairwise comparison of the log-transformed genotype visitation frequency during adaptive (Kimura) walks in the absence of noise (𝒢^*S*^, x-axis) and after allowing for empirically estimated measurement noise in regulation strengths (𝒢^*S,τ*^, y-axis), for the CRP, Fis, and IHF landscapes. Each circle represents visitation data from a different genotype, i.e., the total number of times the genotype was visited across all adaptive walks. We computed regulation strength as a continuous weighted average of fluorescence across sorting bins based on sequencing read distributions (Methods), and incorporated experimental uncertainty as described in **Supplementary Methods 10**. Point density is indicated by color, with warmer colors denoting higher densities of genotypes. The red line shows the best-fit linear model. Spearman correlation coefficients (R) together with p-values for a test of the null hypothesis that genotype visitation rankings are uncorrelated between noisy and noise-free landscapes. The strong correlations observed for all three landscapes indicate that genotype visitation frequencies—and thus landscape-level evolutionary accessibility patterns—are robust to realistic levels of experimental uncertainty. R^2^: coefficient of determination. The panels show correlation plots for **a.** CRP, **b.** Fis, **c.** IHF.

## REFERENCES

1. Browning, D. F. & Busby, S. J. W. Local and global regulation of transcription initiation in bacteria. Nat Rev Microbiol 14, 638–650 (2016).

2. Cases, I., De Lorenzo, V. & Ouzounis, C. A. Transcription regulation and environmental adaptation in bacteria. Trends in Microbiology vol. 11 248–253 Preprint at 10.1016/S0966-842X(03)00103-3 (2003).

3. Seshasayee, A. S., Bertone, P., Fraser, G. M. & Luscombe, N. M. Transcriptional regulatory networks in bacteria: from input signals to output responses. Curr Opin Microbiol 9, 511–519 (2006).

4. Barnard, A., Wolfe, A. & Busby, S. Regulation at complex bacterial promoters: How bacteria use different promoter organizations to produce different regulatory outcomes. Curr Opin Microbiol 7, 102–108 (2004).

5. Browning, D. F. D. F. & Busby, S. J. W. S. The regulation of bacterial transcription initiation. Nat Rev Microbiol 2, 57–65 (2004).

6. Aguilar-Rodríguez, J., Payne, J. L. & Wagner, A. A thousand empirical adaptive landscapes and their navigability. Nat Ecol Evol 1, 0045 (2017).

7. Sharon, E. et al. Inferring gene regulatory logic from high-throughput measurements of thousands of systematically designed promoters. Nat Biotechnol 30, 521–530 (2012).

8. Shultzaberger, R. K., Malashock, D. S., Kirsch, J. F. & Eisen, M. B. The Fitness Landscapes of cis-Acting Binding Sites in Different Promoter and Environmental Contexts. PLoS Genet 6, e1001042 (2010).

9. Haldane, A., Manhart, M. & Morozov, A. V. Biophysical Fitness Landscapes for Transcription Factor Binding Sites. PLoS Comput Biol 10, 36–38 (2014).

10. Shen-Orr, S. S., Milo, R., Mangan, S. & Alon, U. Network motifs in the transcriptional regulation network of Escherichia coli. Nat Genet 31, 64–68 (2002).

11. Martínez-Antonio, A. et al. Identifying global regulators in transcriptional regulatory networks in bacteria. Curr Opin Microbiol 6, 482–489 (2003).

12. Lozada-Chávez, I., Angarica, V. E., Collado-Vides, J. & Contreras-Moreira, B. The Role of DNA-binding Specificity in the Evolution of Bacterial Regulatory Networks. J Mol Biol 379, 627–643 (2008).

13. Kurafeiski, J. D., Pinto, P. & Bornberg-Bauer, E. Evolutionary potential of cis-regulatory mutations to cause rapid changes in transcription factor binding. Genome Biol Evol 11, 406–414 (2019).

14. Browning, D. F., Butala, M. & Busby, S. J. W. Bacterial Transcription Factors: Regulation by Pick “N” Mix. J Mol Biol 431, 4067–4077 (2019).

15. Visweswariah, S. S. & Busby, S. J. W. Evolution of bacterial transcription factors: How proteins take on new tasks, but do not always stop doing the old ones. Trends Microbiol 23, 463–467 (2015).

16. Vaishnav, E. D. et al. The evolution, evolvability and engineering of gene regulatory DNA. Nature 2022 603:7901 603, 455–463 (2022).

17. Schweizer, G. & Wagner, A. Both Binding Strength and Evolutionary Accessibility Affect the Population Frequency of Transcription Factor Binding Sequences in Arabidopsis thaliana. Genome Biol Evol 13, (2021).

18. Westmann, C. A., Goldbach, L. & Wagner, A. The highly rugged yet navigable regulatory landscape of the bacterial transcription factor TetR. Nat Commun 15, 10745 (2024).

19. Westmann, C. A., Goldbach, L. & Wagner, A. Entangled adaptive landscapes facilitate the evolution of gene regulation by exaptation. bioRxiv 2024.11.10.620926 (2024) doi:10.1101/2024.11.10.620926.

20. Wright, S. The roles of mutation, inbreeding, crossbreeding and selection in evolution. Proc of the 6th International Congress of Genetics Preprint at (1932).

21. Wright, S. Evolution in Mendelian Populations. Genetics 16, 97 (1931).

22. Blanco, C., Janzen, E., Pressman, A., Saha, R. & Chen, I. A. Molecular Fitness Landscapes from High-Coverage Sequence Profiling. Annu Rev Biophys 48, 1–18 (2019).

23. Yi, X. & Dean, A. M. Adaptive Landscapes in the Age of Synthetic Biology. Mol Biol Evol 36, 890–907 (2019).

24. Bank, C. Epistasis and Adaptation on Fitness Landscapes. 10.1146/annurev-ecolsys-102320-112153 53, 457–479 (2022).

25. Fragata, I., Blanckaert, A., Dias Louro, M. A., Liberles, D. A. & Bank, C. Evolution in the light of fitness landscape theory. Trends Ecol Evol 34, 69–82 (2019).

26. Otwinowski, J. & Nemenman, I. Genotype to Phenotype Mapping and the Fitness Landscape of the E. coli lac Promoter. PLoS One 8, e61570 (2013).

27. Chattopadhyay, G., Papkou, A. & Wagner, A. The fitness landscape of the E.coli lac operator is highly rugged in two different environments. bioRxiv 2025.07.23.666252 (2025) doi:10.1101/2025.07.23.666252.

28. Herrera-Álvarez, S., Patton, J. E. J. & Thornton, J. W. The structure of an ancient genotype–phenotype map shaped the functional evolution of a protein family. Nat Ecol Evol 9, 1656–1669 (2025).

29. Weeks, R. & Ostermeier, M. Fitness and Functional Landscapes of the E. coli RNase III Gene rnc. Mol Biol Evol 40, (2023).

30. Steinberg, B. & Ostermeier, M. Environmental changes bridge evolutionary valleys. Sci Adv 2, e1500921 (2016).

31. Papkou, A., Garcia-Pastor, L., Escudero, J. A. & Wagner, A. A rugged yet easily navigable fitness landscape. Science (1979) 382, eadh3860 (2023).

32. Sarkisyan, K. S. et al. Local fitness landscape of the green fluorescent protein. Nature 2015 533:7603 533, 397–401 (2016).

33. Schuster, P. A testable genotype-phenotype map: modeling evolution of RNA molecules. Biological Evolution and Statistical Physics. 55–81 (2002) doi:10.1007/3-540-45692-9_4.

34. García-Galindo, P., Ahnert, S. E. & Martin, N. S. The non-deterministic genotype–phenotype map of RNA secondary structure. J R Soc Interface 20, (2023).

35. Bendixsen, D. P., Collet, J., Østman, B. & Hayden, E. J. Genotype network intersections promote evolutionary innovation. PLoS Biol 17, e3000300 (2019).

36. de Visser, J. A. G. M., Cooper, T. F. & Elena, S. F. The causes of epistasis. Proceedings of the Royal Society B: Biological Sciences 278, 3617–3624 (2011).

37. Wray, G. A. The evolutionary significance of cis-regulatory mutations. Nat Rev Genet 8, 206–216 (2007).

38. Signor, S. A. & Nuzhdin, S. V. The Evolution of Gene Expression in cis and trans. Trends in Genetics 34, 532–544 (2018).

39. Wittkopp, P. J., Haerum, B. K. & Clark, A. G. Evolutionary changes in cis and trans gene regulation. Nature 430, 85–88 (2004).

40. Yona, A. H., Alm, E. J. & Gore, J. Random sequences rapidly evolve into de novo promoters. Nat Commun 9, 111880 (2018).

41. Fuqua, T. & Wagner, A. Mobile DNA is replete with hotspots for the de novo emergence of gene regulation. bioRxiv 2023.10.22.563463 (2023) doi:10.1101/2023.10.22.563463.

42. McAdams, H. H., Srinivasan, B. & Arkin, A. P. The evolution of genetic regulatory systems in bacteria. Nature Reviews Genetics vol. 5 169–178 Preprint at 10.1038/nrg1292 (2004).

43. Dorman, C. J., Bhriain, N. N. & Dorman, M. J. The Evolution of Gene Regulatory Mechanisms in Bacteria. in 125–152 (2018). doi:10.1007/978-3-319-69078-0_6.

44. Kinney, J. B. & McCandlish, D. M. Massively Parallel Assays and Quantitative Sequence–Function Relationships. Annu Rev Genomics Hum Genet 20, annurev-genom-083118-014845 (2019).

45. de Visser, J. A. G. M., Elena, S. F., Fragata, I. & Matuszewski, S. The utility of fitness landscapes and big data for predicting evolution. Heredity 2018 121:5 121, 401–405 (2018).

46. Draghi, J. A. & Ogbunugafor, C. B. Exploring the expanse between theoretical questions and experimental approaches in the modern study of evolvability. J Exp Zool B Mol Dev Evol 340, 8–17 (2023).

47. Louis, A. A. Contingency, convergence and hyper-astronomical numbers in biological evolution. Studies in History and Philosophy of Science Part C: Studies in History and Philosophy of Biological and Biomedical Sciences 58, 107–116 (2016).

48. D’Haeseleer, P. What are DNA sequence motifs? Nature Biotechnology 2006 24:4 24, 423–425 (2006).

49. Stormo, G. D. DNA binding sites: Representation and discovery. Bioinformatics vol. 16 16–23 Preprint at 10.1093/bioinformatics/16.1.16 (2000).

50. Stormo, G. D. & Zhao, Y. Determining the specificity of protein–DNA interactions. Nat Rev Genet 11, 751–760 (2010).

51. Zhou, T. et al. Quantitative modeling of transcription factor binding specificities using DNA shape. Proc Natl Acad Sci U S A 112, 4654–4659 (2015).

52. Mathelier, A. et al. DNA Shape Features Improve Transcription Factor Binding Site Predictions In Vivo. Cell Syst 10.1016/j.cels.2016.07.001(2016) doi:10.1016/j.cels.2016.07.001.

53. Ibarra, I. L. et al. Mechanistic insights into transcription factor cooperativity and its impact on protein-phenotype interactions. Nature Communications 2020 11:1 11, 1–16 (2020).

54. Bintu, L. et al. Transcriptional regulation by the numbers: models. Curr Opin Genet Dev 15, 116–124 (2005).

55. Jones, D. L., Brewster, R. C. & Phillips, R. Promoter architecture dictates cell-to-cell variability in gene expression. Science (1979) 346, 1533–1536 (2014).

56. Scholz, S. A. et al. High-Resolution Mapping of the Escherichia coli Chromosome Reveals Positions of High and Low Transcription. Cell Syst 10.1016/j.cels.2019.02.004(2019) doi:10.1016/j.cels.2019.02.004.

57. Kim, D. et al. Systems assessment of transcriptional regulation on central carbon metabolism by Cra and CRP. Nucleic Acids Res 10.1093/nar/gky069(2018) doi:10.1093/nar/gky069.

58. Borirak, O. et al. Time-series analysis of the transcriptome and proteome of Escherichia coli upon glucose repression. Biochimica et Biophysica Acta (BBA) - Proteins and Proteomics 1854, 1269–1279 (2015).

59. Pal, A., Iyer, M. S., Srinivasan, S., Seshasayee, A. S. N. & Venkatesh, K. V. Global pleiotropic effects in adaptively evolved Escherichia coli lacking CRP reveal molecular mechanisms that define the growth physiology. Open Biol 12, (2022).

60. Khankal, R., Chin, J. W., Ghosh, D. & Cirino, P. C. Transcriptional effects of CRP* expression in Escherichia coli. J Biol Eng 3, 1–14 (2009).

61. Nowak-Lovato, K. et al. Binding of Nucleoid-Associated Protein Fis to DNA Is Regulated by DNA Breathing Dynamics. PLoS Comput Biol 9, (2013).

62. Kahramanoglou, C. et al. Direct and indirect effects of H-NS and Fis on global gene expression control in Escherichia coli. Nucleic Acids Res 39, 2073–2091 (2011).

63. Gawade, P., Gunjal, G., Sharma, A. & Ghosh, P. Reconstruction of transcriptional regulatory networks of Fis and H-NS in Escherichia coli from genome-wide data analysis. Genomics 112, 1264–1272 (2020).

64. Prieto, A. I. et al. Genomic analysis of DNA binding and gene regulation by homologous nucleoid-associated proteins IHF and HU in Escherichia coli K12. Nucleic Acids Res 40, 3524–3537 (2012).

65. Arfin, S. M. et al. Global gene expression profiling in Escherichia coli K12. The effects of integration host factor. J Biol Chem 275, 29672–29684 (2000).

66. Taylor, L. R. & Provine, W. B. Sewall Wright and Evolutionary Biology. J Anim Ecol 10.2307/5082(2006) doi:10.2307/5082.

67. Lynch, M. et al. Genetic drift, selection and the evolution of the mutation rate. Nature Reviews Genetics 2016 17:11 17, 704–714 (2016).

68. Iwasa, Y., Michor, F. & Nowak, M. A. Stochastic tunnels in evolutionary dynamics. Genetics 166, 1571–1579 (2004).

69. Taylor, C. F. & Higgs, P. G. A population genetics model for multiple quantitative traits exhibiting pleiotropy and epistasis. J Theor Biol 203, 419–437 (2000).

70. Salgado, H. et al. RegulonDB v12.0: a comprehensive resource of transcriptional regulation in E. coli K-12. Nucleic Acids Res 52, D255–D264 (2024).

71. Peterman, N. & Levine, E. Sort-seq under the hood: Implications of design choices on large-scale characterization of sequence-function relations. BMC Genomics 17, 1–17 (2016).

72. de Boer, C. G. et al. Deciphering eukaryotic gene-regulatory logic with 100 million random promoters. Nature Biotechnology 2019 38:1 38, 56–65 (2019).

73. Feng, H. et al. Deep-learning-assisted Sort-Seq enables high-throughput profiling of gene expression characteristics with high precision. Sci Adv 9, eadg5296 (2023).

74. Garcia, H. G., Lee, H. J., Boedicker, J. Q. & Phillips, R. Comparison and Calibration of Different Reporters for Quantitative Analysis of Gene Expression. Biophys J 101, 535 (2011).

75. Friedlander, T., Prizak, R., Barton, N. H. & Tkačik, G. Evolution of new regulatory functions on biophysically realistic fitness landscapes. Nat Commun 8, (2017).

76. Le, D. D. et al. Comprehensive, high-resolution binding energy landscapes reveal context dependencies of transcription factor binding. Proceedings of the National Academy of Sciences 115, 201715888 (2018).

77. Maerkl, S. J. S. J. & Quake, S. R. A systems approach to measuring the binding energy landscapes of transcription factors. Science (1979) 315, (2007).

78. Poelwijk, F. J., Kiviet, D. J., Weinreich, D. M. & Tans, S. J. Empirical fitness landscapes reveal accessible evolutionary paths. Nature 10.1038/nature05451(2007) doi:10.1038/nature05451.

79. Nghe, P. et al. Predicting Evolution Using Regulatory Architecture. Annu Rev Biophys 49, 181–197 (2020).

80. Struhl, K. Fundamentally different logic of gene regulation in eukaryotes and prokaryotes. Cell 98, 1–4 (1999).

81. Barnes, S. L., Belliveau, N. M., Ireland, W. T., Kinney, J. B. & Phillips, R. Mapping DNA sequence to transcription factor binding energy in vivo. PLoS Comput Biol 15, e1006226 (2019).

82. Kolb, A., Spassky, A., Chapon, C., Blazy, B. & Buc, H. On the different binding affinities of CRP at the lac, gal and malt promoter regions. Nucleic Acids Res 10.1093/nar/11.22.7833(1983) doi:10.1093/nar/11.22.7833.

83. Shao, Y., Feldman-Cohen, L. S. & Osuna, R. Functional Characterization of the Escherichia coli Fis-DNA Binding Sequence. J Mol Biol 376, 771–785 (2008).

84. Huo, Y. X. et al. IHF-binding sites inhibit DNA loop formation and transcription initiation. Nucleic Acids Res 37, 3878–3886 (2009).

85. Gunasekera, A., Ebright, Y. W. & Ebright, R. H. DNA sequence determinants for binding of the Escherichia coli catabolite gene activator protein. Journal of Biological Chemistry 267, 14713–14720 (1992).

86. Aiyar, S. E. et al. Architecture of fis-activated transcription complexes at the Escherichia coli rrnB P1 and rrnE P1 promoters. J Mol Biol 316, 501–516 (2002).

87. Ho, S. Characteristics of IHF Binding to Holliday Junction DNA. (2013).

88. Song, S. & Zhang, J. Unbiased inference of the fitness landscape ruggedness from imprecise fitness estimates. Evolution (N Y) 75, 2658–2671 (2021).

89. Obolski, U., Ram, Y. & Hadany, L. Key issues review: evolution on rugged adaptive landscapes. Reports on Progress in Physics 81, 012602 (2018).

90. Kauffman, S. & Levin, S. Towards a general theory of adaptive walks on rugged landscapes. J Theor Biol 128, 11–45 (1987).

91. Weinberger, E. Correlated and uncorrelated fitness landscapes and how to tell the difference. Biol Cybern 63, 325–336 (1990).

92. Orr, H. A. A Minimum on the Mean Number of Steps Taken in Adaptive Walks. J Theor Biol 220, 241–247 (2003).

93. Saona, R., Kondrashov, F. A. & Khudiakova, K. A. Relation Between the Number of Peaks and the Number of Reciprocal Sign Epistatic Interactions. Bull Math Biol 84, (2022).

94. Poelwijk, F. J., Tǎnase-Nicola, S., Kiviet, D. J. & Tans, S. J. Reciprocal sign epistasis is a necessary condition for multi-peaked fitness landscapes. J Theor Biol 272, 141–144 (2011).

95. Kvitek, D. J. & Sherlock, G. Reciprocal sign epistasis between frequently experimentally evolved adaptive mutations causes a rugged fitness landscape. PLoS Genet 10.1371/journal.pgen.1002056(2011) doi:10.1371/journal.pgen.1002056.

96. Conrad, M. The geometry of evolution. Biosystems 24, 61–81 (1990).

97. Krug, J. & Oros, D. Evolutionary accessibility of random and structured fitness landscapes. https://arxiv.org/abs/2311.17432v2 (2023).

98. Kimura, M. On the Probability of Fixation of Mutant Genes in a Population. Genetics 47, 713–719 (1962).

99. Bank, C., Matuszewski, S., Hietpas, R. T. & Jensen, J. D. On the (un)predictability of a large intragenic fitness landscape. Proc Natl Acad Sci U S A 113, 14085–14090 (2016).

100. Orr, H. A. The population genetics of adaptation: the adaptation of DNA sequences. Evolution 56, 1317–1330 (2002).

101. Gillespie, J. H. MOLECULAR EVOLUTION OVER THE MUTATIONAL LANDSCAPE. Evolution (N Y) 38, 1116–1129 (1984).

102. Lind, P. A. & Andersson, D. I. Whole-genome mutational biases in bacteria. Proceedings of the National Academy of Sciences 105, 17878–17883 (2008).

103. Horton, J. S. & Taylor, T. B. Mutation bias and adaptation in bacteria. Microbiology (N Y) 169, (2023).

104. Lee, H., Popodi, E., Tang, H. & Foster, P. L. Rate and molecular spectrum of spontaneous mutations in the bacterium *Escherichia coli* as determined by whole-genome sequencing. Proceedings of the National Academy of Sciences 109, (2012).

105. Melissa, M. J., Good, B. H., Fisher, D. S. & Desai, M. M. Population genetics of polymorphism and divergence in rapidly evolving populations. Genetics 221, (2022).

106. Li, J., Amado, A. & Bank, C. Rapid adaptation of recombining populations on tunable fitness landscapes. Mol Ecol 10.1111/MEC.16900(2023) doi:10.1111/MEC.16900.

107. Stolyarova, A. V. et al. Complex fitness landscape shapes variation in a hyperpolymorphic species. Elife 11, (2022).

108. Park, S.-C., Neidhart, J. & Krug, J. Greedy adaptive walks on a correlated fitness landscape. J Theor Biol 397, 89–102 (2016).

109. Kauffman, S. & Levin, S. Towards a general theory of adaptive walks on rugged landscapes. J Theor Biol 128, 11–45 (1987).

110. Orr, H. A. A Minimum on the Mean Number of Steps Taken in Adaptive Walks. J Theor Biol 220, 241–247 (2003).

111. Blount, Z. D., Lenski, R. E. & Losos, J. B. Contingency and determinism in evolution: Replaying life’s tape. Science (1979) 362, (2018).

112. Nonoyama, T. & Chiba, S. Phenotypic determinism and contingency in the evolution of hypothetical tree-like organisms. PLoS One 14, (2019).

113. Xie, V. C., Pu, J., Metzger, B. P. H., Thornton, J. W. & Dickinson, B. C. Contingency and chance erase necessity in the experimental evolution of ancestral proteins. Elife 10, 1–87 (2021).

114. Spor, A. et al. Phenotypic and genotypic convergences are influenced by historical contingency and environment in yeast. Evolution 68, 772 (2014).

115. Lenski, R. E. Convergence and Divergence in a Long-Term Experiment with Bacteria*. 10.1086/691209 190, S57–S68 (2017).

116. Lind, P. A., Farr, A. D. & Rainey, P. B. Experimental evolution reveals hidden diversityin evolutionary pathways. Elife 2015, (2015).

117. Trippe, B. L. et al. Randomized gates eliminate bias in sort-seq assays. Protein Science 31, e4401 (2022).

118. Gilliot, P. A. & Gorochowski, T. E. Effective design and inference for cell sorting and sequencing based massively parallel reporter assays. Bioinformatics 39, (2023).

119. Evfratov, S. A. et al. Application of sorting and next generation sequencing to study 5΄-UTR influence on translation efficiency in Escherichia coli. Nucleic Acids Res 45, 3487–3502 (2017).

120. Cuperus, J. T. et al. Deep learning of the regulatory grammar of yeast 5 ′ untranslated regions from 500, 000 random sequences. Genome Res 163, 1–10 (2017).

121. Belliveau, N. M. et al. A Systematic and Scalable Approach for Dissecting the Molecular Mechanisms of Transcriptional Regulation in Bacteria. Biophys J 114, 151a (2018).

122. Kinney, J. B., Murugan, A., Callan, C. G. & Cox, E. C. Using deep sequencing to characterize the biophysical mechanism of a transcriptional regulatory sequence. Proceedings of the National Academy of Sciences 107, 9158–9163 (2010).

123. Lagator, M. et al. Predicting bacterial promoter function and evolution from random sequences. Elife 11, (2022).

124. Urtecho, G., Tripp, A. D., Insigne, K. D., Kim, H. & Kosuri, S. Systematic Dissection of Sequence Elements Controlling σ70 Promoters Using a Genomically Encoded Multiplexed Reporter Assay in Escherichia coli. Biochemistry 58, 1539–1551 (2019).

125. Aguilar-Rodríguez, J. & Payne, J. L. Robustness and Evolvability in Transcriptional Regulation. Evolutionary Systems Biology 197–219 (2021) doi:10.1007/978-3-030-71737-7_9.

126. Lagator, M., Igler, C., Moreno, A. B., Guet, C. C. & Bollback, J. P. Epistatic interactions in the arabinose cis-regulatory element. Mol Biol Evol 33, 761–769 (2016).

127. Lagator, M., Paixão, T., Barton, N. H., Bollback, J. P. & Guet, C. C. On the mechanistic nature of epistasis in a canonical cis-regulatory element. Elife 6, 1–16 (2017).

128. Stewart, A. J., Hannenhalli, S., Plotkin, J. B., Hannenhalli, S. & Plotkin, J. B. Why Transcription Factor Binding Sites Are Ten Nucleotides Long. Genetics 192, 973–985 (2012).

129. Babu, M. M. & Teichmann, S. A. Functional determinants of transcription factors in Escherichia coli: Protein families and binding sites. Trends in Genetics 19, 75–79 (2003).

130. Huffman, J. L. & Brennan, R. G. Prokaryotic transcription regulators: more than just the helix-turn-helix motif. Curr Opin Struct Biol 12, 98–106 (2002).

131. Perez-Rueda, E. et al. Abundance, diversity and domain architecture variability in prokaryotic DNA-binding transcription factors. PLoS One 13, (2018).

132. Oren, Y. et al. Transfer of noncoding DNA drives regulatory rewiring in bacteria. Proceedings of the National Academy of Sciences 111, 16112–16117 (2014).

133. Price, M. N., Dehal, P. S. & Arkin, A. P. Horizontal gene transfer and the evolution of transcriptional regulation in Escherichia coli. Genome Biol 9, (2008).

134. Weinreich, D. M., Watson, R. A. & Chao, L. Perspective: Sign epistasis and genetic constraint on evolutionary trajectories. Evolution (2005).

135. Gerland, U., Moroz, J. D. & Hwa, T. Physical constraints and functional characteristics of transcription factor-DNA interaction. Proceedings of the National Academy of Sciences 99, 12015–12020 (2002).

136. Zhao, Y., Ruan, S., Pandey, M. & Stormo, G. D. Improved models for transcription factor binding site identification using nonindependent interactions. Genetics 10.1534/genetics.112.138685(2012) doi:10.1534/genetics.112.138685.

137. O’Flanagan, R. A., Paillard, G., Lavery, R. & Sengupta, A. M. Non-additivity in protein–DNA binding. Bioinformatics 21, 2254–2263 (2005).

138. Payne, J. L. & Wagner, A. The Robustness and Evolvability of Transcription Factor Binding Sites. Science (1979) 343, (2014).

139. Blount, Z. D., Borland, C. Z. & Lenski, R. E. Historical contingency and the evolution of a key innovation in an experimental population of Escherichia coli. Proc Natl Acad Sci U S A 105, 7899–7906 (2008).

140. Palmer, A. C. et al. Delayed commitment to evolutionary fate in antibiotic resistance fitness landscapes. Nature Communications 2015 6:1 6, 1–8 (2015).

141. Vermeij, G. J. Historical contingency and the purported uniqueness of evolutionary innovations. Proceedings of the National Academy of Sciences 103, 1804–1809 (2006).

142. Lässig, M., Mustonen, V. & Walczak, A. M. Predicting evolution. Nat Ecol Evol 1, 1–9 (2017).

143. De Visser, J. A. G. M. & Krug, J. Empirical fitness landscapes and the predictability of evolution. Nat Rev Genet 15, 480–490 (2014).

144. Heyde, S. A. H., Frendorf, P. O., Lauritsen, I. & Nørholm, M. H. H. Restoring Global Gene Regulation through Experimental Evolution Uncovers a NAP (Nucleoid-Associated Protein)-Like Behavior of Crp/Cap. mBio 12, (2021).

145. Monteiro, L. M. O., Sanches-Medeiros, A., Westmann, C. A. & Silva-Rocha, R. Unraveling the Complex Interplay of Fis and IHF Through Synthetic Promoter Engineering. Front Bioeng Biotechnol 8, 510 (2020).

146. Dillon, S. C. & Dorman, C. J. Bacterial nucleoid-associated proteins, nucleoid structure and gene expression. Nat Rev Microbiol 8, 185–95 (2010).

147. Dorman, C. J. Genome architecture and global gene regulation in bacteria: Making progress towards a unified model? Nat Rev Microbiol 11, 349–355 (2013).

148. Dorman, C. J., Schumacher, M. A., Bush, M. J., Brennan, R. G. & Buttner, M. J. When is a transcription factor a NAP? Curr Opin Microbiol 55, 26 (2020).

149. Chen, Y. J. et al. Quantifying molecular bias in DNA data storage. Nature Communications 2020 11:1 11, 1–9 (2020).

150. Kebschull, J. M. & Zador, A. M. Sources of PCR-induced distortions in high-throughput sequencing data sets. Nucleic Acids Res 43, e143–e143 (2015).

151. Wong, T. S., Zhurina, D. & Schwaneberg, U. The Diversity Challenge in Directed Protein Evolution. Comb Chem High Throughput Screen 9, 271–288 (2006).

152. Aird, D. et al. Analyzing and minimizing PCR amplification bias in Illumina sequencing libraries. Genome Biol 12, 1–14 (2011).

153. Gilliot, P.-A. & Gorochowski, T. E. Effective design and inference for cell sorting and sequencing based massively parallel reporter assays. bioRxiv 2022.11.07.515414 (2022) doi:10.1101/2022.11.07.515414.

154. Wagner, A. AI predicts the effectiveness and evolution of gene promoter sequences. Nature 2022 603:7901 603, 399–400 (2022).

155. Boer, C. de, Sadeh, R., Friedman, N. & Regev, A. Deciphering cis-regulatory logic with 100 million random promoters. bioRxiv 224907 (2018) doi:10.1101/224907.

156. De Boer, C. G., Ray, J. P., Hacohen, N. & Regev, A. MAUDE: Inferring expression changes in sorting-based CRISPR screens. Genome Biol 21, 1–16 (2020).

157. Wu, N. C., Dai, L., Olson, C. A., Lloyd-Smith, J. O. & Sun, R. Adaptation in protein fitness landscapes is facilitated by indirect paths. Elife 5, (2016).

158. Diaz-Colunga, J., et al. Global epistasis on fitness landscapes. Philosophical Transactions of the Royal Society B 378, (2023).

159. MacLean, R. C., Perron, G. G. & Gardner, A. Diminishing Returns From Beneficial Mutations and Pervasive Epistasis Shape the Fitness Landscape for Rifampicin Resistance in Pseudomonas aeruginosa. Genetics 186, 1345–1354 (2010).

160. Rydenfelt, M., Garcia, H. G., Cox, R. S. & Phillips, R. The influence of promoter architectures and regulatory motifs on gene expression in Escherichia coli. PLoS One 9, 1–31 (2014).

161. Ishihama, A. Prokaryotic genome regulation: multifactor promoters, multitarget regulators and hierarchic networks. FEMS Microbiol Rev 34, 628–645 (2010).

162. Cases, I. & de Lorenzo, V. Promoters in the environment: Transcriptional regulation in its natural context. Nature Reviews Microbiology vol. 3 105–118 Preprint at 10.1038/nrmicro1084 (2005).

163. Shepherd, M. J., Reynolds, M., Pierce, A. P., Rice, A. M. & Taylor, T. B. Transcription factor expression levels and environmental signals constrain transcription factor innovation. Microbiology (N Y) 169, (2023).

164. González Pérez, A., Espinosa Angarica, V., Collado-Vides, J. & Vasconcelos, A. T. R. From sequence to dynamics: The effects of transcription factor and polymerase concentration changes on activated and repressed promoters. BMC Mol Biol 10, 92 (2009).

165. Azam, T. A., Iwata, A., Nishimura, A., Ueda, S. & Ishihama, A. Growth phase-dependent variation in protein composition of the Escherichia coli nucleoid. J Bacteriol (1999) doi:.

166. Weinert, F. M. et al. The Transcription Factor Titration Effect Dictates Level of Gene Expression. Biophys J 10.1016/j.bpj.2013.11.4483(2014) doi:10.1016/j.bpj.2013.11.4483.

167. Lagomarsino, M. C., Espéli, O. & Junier, I. From structure to function of bacterial chromosomes: Evolutionary perspectives and ideas for new experiments. FEBS Letters Preprint at 10.1016/j.febslet.2015.07.002 (2015).

168. Dorman, C. J. & Dorman, M. J. DNA supercoiling is a fundamental regulatory principle in the control of bacterial gene expression. Biophys Rev 8, 209–220 (2016).

169. Houdaigui, B. El et al. Bacterial genome architecture shapes global transcriptional regulation by DNA supercoiling. Nucleic Acids Res 47, 5648–5657 (2019).

170. Sobetzko, P., Travers, A. & Muskhelishvili, G. Gene order and chromosome dynamics coordinate spatiotemporal gene expression during the bacterial growth cycle. Proceedings of the National Academy of Sciences 109, E42–E50 (2011).

171. Meyer, S., Reverchon, S., Nasser, W. & Muskhelishvili, G. Chromosomal organization of transcription: in a nutshell. Curr Genet 64, 555–565 (2018).

172. Sánchez-Romero, M. A., Cota, I. & Casadesús, J. DNA methylation in bacteria: from the methyl group to the methylome. Curr Opin Microbiol 25, 9–16 (2015).

173. Crocker, J., Preger-Ben Noon, E. & Stern, D. L. The Soft Touch: Low-Affinity Transcription Factor Binding Sites in Development and Evolution. Current Topics in Developmental Biology vol. 117 (Elsevier Inc., 2016).

174. Srivastava, M. & Payne, J. L. On the incongruence of genotype-phenotype and fitness landscapes. PLoS Comput Biol 18, e1010524 (2022).

175. Rest, J. S. et al. Nonlinear fitness consequences of variation in expression level of a eukaryotic gene. Mol Biol Evol 30, 448–456 (2013).

176. Duveau, F., Toubiana, W. & Wittkopp, P. J. Fitness effects of cis-regulatory variants in the saccharomyces cerevisiae TDH3 promoter. Mol Biol Evol 34, (2017).

177. Bergen, A. C., Olsen, G. M. & Fay, J. C. Divergent MLS1 Promoters Lie on a Fitness Plateau for Gene Expression. Mol Biol Evol 33, (2016).

178. Ornelas, M. Y., Cournoyer, J. E., Bram, S. & Mehta, A. P. Evolution and synthetic biology. Curr Opin Microbiol 76, 102394 (2023).

179. Baba, T. et al. Construction of Escherichia coli K-12 in-frame, single-gene knockout mutants: The Keio collection. Mol Syst Biol 2, 2006.0008 (2006).

180. Elowitz, M. B., Levine, A. J., Siggia, E. D. & Swain, P. S. Stochastic Gene Expression in a Single Cell. Science (1979) 297, 1183–6 (2007).

181. Sharon, E. et al. Probing the effect of promoters on noise in gene expression using thousands of designed sequences. Genome Res 24, 1698–1706 (2014).

182. Rudge, T. J. et al. Characterization of Intrinsic Properties of Promoters. ACS Synth Biol 5, 89–98 (2016).

183. Gibson, D. G. et al. Enzymatic assembly of DNA molecules up to several hundred kilobases. Nat Methods 6, 343–5 (2009).

184. Warren, D. J. Preparation of highly efficient electrocompetent Escherichia coli using glycerol/mannitol density step centrifugation. Anal Biochem 413, 206–207 (2011).

185. Jahn, M., Vorpahl, C., Hübschmann, T., Harms, H. & Müller, S. Copy number variability of expression plasmids determined by cell sorting and droplet digital PCR. Microb Cell Fact 15, 211 (2016).

186. Lutz, R. & Bujard, H. Independent and tight regulation of transcriptional units in Escherichia coli via the LacR/O, the TetR/O and AraC/I1-I2 regulatory elements. Nucleic Acids Res 25, 1203–10 (1997).

187. Pédelacq, J. D., Cabantous, S., Tran, T., Terwilliger, T. C. & Waldo, G. S. Engineering and characterization of a superfolder green fluorescent protein. Nat Biotechnol 24, 79–88 (2006).

188. Lou, C., Stanton, B., Chen, Y. J., Munsky, B. & Voigt, C. A. Ribozyme-based insulator parts buffer synthetic circuits from genetic context. Nat Biotechnol 10.1038/nbt.2401(2012) doi:10.1038/nbt.2401.

189. Bindels, D. S. et al. mScarlet: a bright monomeric red fluorescent protein for cellular imaging. Nature Methods 2016 14:1 14, 53–56 (2016).

190. Beal, J. Biochemical complexity drives log-normal variation in genetic expression. Engineering Biology 1, 55–60 (2017).

191. Beal, J. et al. Reproducibility of fluorescent expression from engineered biological constructs in E. coli. PLoS One 11, (2016).

192. Martin, M. Cutadapt removes adapter sequences from high-throughput sequencing reads. EMBnet J 17, 10–12 (2011).

193. Magoč, T. & Salzberg, S. L. FLASH: fast length adjustment of short reads to improve genome assemblies. Bioinformatics 27, 2957–2963 (2011).

194. Hannon, G. J. FASTX-Toolkit.

195. Bunn, A. & Korpela, M. R: A language and environment for statistical computing. 10.1016/j.dendro.2008.01.002(2013) doi:10.1016/j.dendro.2008.01.002.

196. Schneider, T. D. & Stephens, R. M. Sequence logos: a new way to display consensus sequences. Nucleic Acids Res 18, 6097 (1990).

197. Csardi, G. The Igraph Software Package for Complex Network Research. https://www.researchgate.net/publication/221995787 (2014).

198. Khalid, F. et al. Genonets server-a web server for the construction, analysis and visualization of genotype networks. Nucleic Acids Res 44, W70–W76 (2016).

199. Jaccard, P. THE DISTRIBUTION OF THE FLORA IN THE ALPINE ZONE.1. New Phytologist 11, 37–50 (1912).

200. Papkou, A., Garcia-Pastor, L., Escudero, J. A. & Wagner, A. A rugged yet easily navigable fitness landscape of antibiotic resistance. bioRxiv 2023.02.27.530293 (2023) doi:10.1101/2023.02.27.530293.

201. Kimura, M. The Neutral Theory of Molecular Evolution. (Cambridge University Press, 1983). doi:10.1017/CBO9780511623486.

202. Crow, J. and Kimura, M. An Introduction to Population Genetics Theory. 608 (The Blackburn Press, 2009).

203. Harris, C. R. et al. Array programming with NumPy. Nature 2020 585:7825 585, 357–362 (2020).

204. Grant, S. G., Jessee, J., Bloom, F. R. & Hanahan, D. Differential plasmid rescue from transgenic mouse DNAs into Escherichia coli methylation-restriction mutants. Proceedings of the National Academy of Sciences 87, 4645–4649 (1990).

205. Ebright, R. H., Ebright, Y. W. & Gunasekera, A. Consensus DNA site for the Escherichia coli catabolite gene activator protein (CAP): CAP exhibits a 450-fold higher affinity for the consensus DNA site than for the E.coli lac DNA site. Nucleic Acids Res 17, 10295–10305 (1989).

206. Shao, Y., Feldman-Cohen, L. S. & Osuna, R. Biochemical Identification of Base and Phosphate Contacts between Fis and a High-Affinity DNA Binding Site. J Mol Biol 380, 327–339 (2008).

207. Kahramanoglou, C. et al. Direct and indirect effects of H-NS and Fis on global gene expression control in Escherichia coli. Nucleic Acids Res 39, 2073–2091 (2011).

208. Kelly, J. R. et al. Measuring the activity of BioBrick promoters using an in vivo reference standard. J Biol Eng 3, 4 (2009).

209. Meyer, A. J., Segall-Shapiro, T. H., Glassey, E., Zhang, J. & Voigt, C. A. Escherichia coli “Marionette” strains with 12 highly optimized small-molecule sensors. Nature Chemical Biology 2018 15:2 15, 196–204 (2018).

210. Carr, S. B., Beal, J. & Densmore, D. M. Reducing DNA context dependence in bacterial promoters. PLoS One 10.1371/journal.pone.0176013 (2017) doi:10.1371/journal.pone.0176013.

211. Chen, Y. J. et al. Characterization of 582 natural and synthetic terminators and quantification of their design constraints. Nat Methods 10.1038/nmeth.2515 (2013) doi:10.1038/nmeth.2515.

212. Wickham, H. Ggplot2: Elegant Graphics for Data Analysis. (Springer-VerlagNew York, 2016).

